# Maternal CENP-C restores centromere symmetry in mammalian zygotes to ensure proper chromosome segregation

**DOI:** 10.1101/2025.07.23.666394

**Authors:** Catherine A. Tower, Gabriel Manske, Emily L. Ferrell, Dilara N. Anbarci, Kelsey Jorgensen, Binbin Ma, Mansour Aboelenain, Rajesh Ranjan, Saikat Chakraborty, Lindsay Moritz, Arunika Das, Michele Boiani, Ben E. Black, Shawn Chavez, Erica E. Marsh, Ariella Shikanov, Karen Schindler, Xin Chen, Saher Sue Hammoud

**Affiliations:** Department of Human Genetics, University of Michigan, Ann Arbor, MI; Cellular and Molecular Biology Graduate Program, University of Michigan, Ann Arbor, MI; Department of Obstetrics and Gynecology, University of Michigan, Ann Arbor, MI; Department of Anthropology, University of Kansas, Lawrence, KS; Howard Hughes Medical Institute, Department of Biology, John Hopkins University, Baltimore, MD; Department of Genetics, Rutgers University, Piscataway, NJ; Centre for Cell Biology, Institute of Cell Biology, University of Edinburgh EH9 3BF, UK; Department of Theriogenology, Faculty of Veterinary Medicine, Mansoura University, Egypt, 35516; Department of Biomedical Sciences, Cornell University College of Veterinary Medicine, Ithaca, NY; Max Planck Institute for Molecular Biomedicine, Department of Cell & Tissue Dynamics, Muenster, Germany; Department of Biochemistry and Biophysics, University of Pennsylvania, Philadelphia, PA; Division of Reproductive and Developmental Sciences, Oregon National Primate Research Center, Beaverton, OR; Department of Molecular and Medical Genetics, Oregon Health and Science University, Portland, OR; Department of Obstetrics and Gynecology, Oregon Health and Science University, Portland, OR 97239, USA; Department of Biomedical Engineering, Oregon Health and Science University, Portland, OR 97239, USA; Department of Biomedical Engineering, University of Michigan, Ann Arbor, MI; Department of Urology, University of Michigan, Ann Arbor, MI

**Author notes:** Contributed equally (Catherine A. Tower, Gabriel Manske, Emily L. Ferrell).

## Abstract

Across metazoan species, the centromere-specific histone variant CENP-A is essential for accurate chromosome segregation, yet its regulation at the parental-to-zygote transition in mammals is poorly understood. To address this, we developed a CENP-A-mScarlet knock-in mouse model, which revealed sex-specific dynamics: mature sperm retains 10% of the CENP-A levels present in MII-oocytes. However, in zygotes prior to the first mitosis, this difference is resolved, using maternally inherited cytoplasmic-CENP-A. Notably, the increase in CENP-A at paternal centromeres is independent of sensing CENP-A asymmetry or the presence of maternal chromosomes. Instead, CENP-A equalization relies on asymmetric recruitment of maternal CENP-C to paternal centromeres. Depletion of maternal CENP-A decreases total CENP-A in pronuclei without disrupting equalization. In contrast, reducing maternal CENP-C or disruption of its dimerization domains impairs CENP-A equalization and chromosome segregation. Therefore, maternal CENP-C acts a key epigenetic regulator that resets centromeric symmetry at fertilization to preserve genome integrity.

**Highlights:**

● CENP-A asymmetry between sperm and oocyte centromeres is a conserved feature from flies to mammals including mice and humans.
● CENP-A asymmetry between parental centromeres is resolved prior to the first zygotic division via maternally inherited, cytoplasmic CENP-A.
● Zygotic CENP-A levels in zygotes are regulated in a pronucleus-autonomous manner.
● CENP-A equalization relies on asymmetric CENP-C recruitment to the paternal pronucleus and requires CENP-C dimerization.

**Key Terms:** Centromere; CENP-A; CENP-C; sperm; oocyte; zygote; intergenerational; epigenetics; mouse

## Introduction

Genome replication and segregation are fundamental processes that safeguard organismal development. During both mitosis and meiosis, centromeres function as organizing nodes for spindle microtubules and the kinetochore to ensure faithful chromosome segregation. As opposed to relying on the underlying DNA sequence, centromeres are epigenetically defined by the presence of nucleosomes containing the histone H3 variant CENP-A (McKinley and Cheeseman, 2016; Palmer et al., 1991;). CENP-A-containing nucleosomes are extremely stable during the cell cycle (Bodor et al., 2013; Hemmerich et al., 2008; Jansen et al., 2007), a property conferred by CENP-A’s direct binding partners CENP-B, CENP-C, CENP-N, and HJURP (Fachinetti et al., 2015; Falk et al., 2015; Guo et al., 2017; Pentakota et al., 2017; Zasadzińska et al., 2018). These interactions form the foundation of the constitutive centromere-associated network (CCAN), a 16 protein heterooligomeric structure that forms the inner kinetochore (Foltz et al., 2006; Pesenti et al., 2022). After mammalian cell division, pre-existing CENP-A nucleosomes restore CENP-A levels by directing new deposition at the end of telophase and the beginning of G1 (Boyarchuk et al., 2014; Hemmerich et al., 2008; Jansen et al., 2007). This one-to-one mechanism of CENP-A nucleosome incorporation and maintenance ensures the precise propagation of centromeric chromatin, preserving both its genomic location and quantitative nucleosome count after mitotic cell division.

CENP-A deposition and maintenance is highly regulated across the cell cycle. It requires at least four core factors: two MIS18 subunits (MIS18α and MIS18β), the MIS18-binding protein 1 (MIS18BP1, also known as KNL2), and the Holliday Junction Recognition Protein (HJURP), as well as several accessory proteins and regulators (Hu et al., 2011; Pan et al., 2019; Spiller et al., 2017). Although the deposition machinery is present throughout the cell cycle, new CENP-A deposition requires integrated signals from polo like kinase 1 (PLK1) and cyclin-dependent kinases (CDK 1&2). At the end of mitosis and early G1 phase, PLK1 phosphorylation of MIS18BP1 and HJURP promotes their localization to centromeres, while high CDK1&2 kinase activity at all other timepoints phosphorylates and sequesters MIS18BP1 and HJURP away from centromeres (Conti et al., 2024, McKinley and Cheeseman, 2014; Müller et al., 2014; Pan et al., 2017; Parashara et al., 2024, Silva et al., 2012; Spiller et al., 2017; Stankovic et al., 2017). Consistently, MIS18 complex formation, HJURP recruitment to centromeres, and CENP-A incorporation are all reduced in the presence of a small molecule PLK1 inhibitor (McKinley and Cheeseman, 2014), whereas inhibition of CDK1 and CDK2 leads to ectopic deposition of CENP-A in G2 and S-phase (Silva et al., 2012; Stankovic et al., 2017). Importantly, dysregulation of any of these pathways renders the genome vulnerable to aneuploidy and genome instability, a hallmark of aging and many diseases including cancer (Barra and Fachinetti, 2018).

Although mechanisms governing CENP-A dynamics and inheritance in mitotic cells have been defined *in vitro*, our understanding of centromere regulation and maintenance *in vivo* remains limited. Studies in *Drosophila* and plants suggest that CENP-A levels at centromeres are dynamically modulated during gametogenesis, with levels decreasing transiently during meiosis, but are subsequently restored in post-meiotic cells (Dunleavy et al., 2012; Raychaudhuri et al., 2012; Schubert et al., 2014). In contrast, previous studies in mice have suggested that CENP-A levels in female germ cells are maintained without active CENP-A deposition. Indeed, conditional knockout models of *Cenpa* or *Mis18α* in early prophase I-arrested mouse oocytes have no effect on centromere maintenance, oocyte maturation, or female fertility (Das et al., 2023; Smoak et al., 2016). These findings indicate that mouse oocytes do not require new CENP-A deposition during their extended period of quiescence, nor is new transcription needed to resume meiotic divisions or support embryonic development. Additional studies showed that *Cenpa^+/-^* mothers have decreased live birth rate and reduced CENP-A abundance in the germline of F1 male offspring (Das et al., 2022). Thus, weakened centromeres are propagated trans-generationally, but interestingly, the epigenetic memory of reduced CENP-A in F1 mice is specifically erased in the female germline. Importantly, the epigenetic memory of *Cenpa*^+/-^ males can be reset in early embryos between 1-to-4 cell if males are mated to wildtype (WT) *Cenpa*^+/+^ females (Das et al., 2022); therefore, this correction process relies on the maternal *Cenpa* genotype. Together, mammalian centromeres utilize genetic and epigenetic information to maintain and correct aberrations to centromeric chromatin from one generation to the next.

Across species, the physical inheritance and requirement of CENP-A at centromeres is variable (Mellone and Fachinetti, 2021). In *Caenorhabditis elegans*, CENP-A is neither inherited nor required for re-establishment of holocentric chromosomes in the early embryo (Gassmann et al., 2012). A similar transient absence of centromeric CenH3 has also been described in egg cells of *Arabidopsis thaliana*, which have regional CENP-A domains similar to mammalian centromeres (Ingouff et al., 2010). In *Drosophila melanogaster*, centromeric proteins such as CENP-C and CAL1, the ortholog of mammalian HJURP, are not retained in sperm (Dunleavy et al., 2012; Raychaudhuri et al., 2012). However, inheriting the CENP-A ortholog, CID, is necessary for embryonic development and essential for propagating the paternal genome to the next generation (Raychaudhuri et al., 2012). Case studies in humans have reported that the location and strength of neocentromeres, including those on the Y chromosome, can be passed down the paternal lineage (Amor et al., 2004; Bukvic et al., 1996; Tyler-Smith et al., 1999). Finally, in mice, somatic CENP-A levels are propagated faithfully from one generation to the next, but the extent and mechanism by which parental *Cenpa* transcripts and centromeric nucleosomes are required to re-establish functional centromeres in the totipotent embryo is unknown.

Here, by quantifying CENP-A dynamics during male and female gametogenesis, we show that centromeric CENP-A levels are elevated in germ cell precursors relative to somatic cells. However, CENP-A loss is differentially regulated between the sexes, leading to sperm retaining roughly a tenth of the CENP-A levels present in mature oocytes. Following fertilization, CENP-A levels increase and equalize between parental chromosomes using maternally inherited, cytoplasmic CENP-A. Analysis of androgenetic embryos reveals that CENP-A accumulation in the paternal pronucleus is an autonomously regulated program – independent of the maternal pronucleus or any CENP-A asymmetry sensing mechanism, suggesting that sperm chromatin state dictates the upper limit of CENP-A levels that can be deposited on centromeres. This equalization process requires maternal CENP-C and its dimerization capacity. While normalization of parental CENP-A levels is necessary, it is not sufficient for successful embryogenesis. Zygotes with insufficient CENP-A can compensate through elevated CENP-C accumulation, highlighting a maternal safeguard mechanism that ensures faithful chromosome segregation and genome stability at the onset of development.

## Results

### Generation and validation of tagged *Cenpa^+/mScarlet^* mouse model

To monitor CENP-A dynamics during mouse development and gametogenesis, we used CRISPR/Cas9 to tag the endogenous *Cenpa* gene at the C-terminus with an enhanced red fluorescent protein mScarlet-I and a V5 epitope tag, connected by a flexible glycine-serine linker (GS) (**Figure 1A**). Sanger sequencing confirmed successful knock-in in two independent founder lines (F0), which were subsequently backcrossed to C57BL/6J prior to phenotypic characterization. Adult male *Cenpa^mScarlet/+^* mice displayed no overt abnormalities, including normal testes-to-body weight ratios and sperm counts compared to WT littermates (**Figure S1A&B**). However, intercrosses of *Cenpa^mScarlet/+^* mice (het × het) yielded no *Cenpa^mScarlet/mScarlet^* embryos (0/18 embryos), while WT (*Cenpa**^+/+^***, 22.2%) and heterozygous (*Cenpa^mScarlet/+^;* 77.8%) embryos were recovered at near-Mendelian ratios (**Supplemental Methods Table 2**). The homozygous lethality underscores a critical role for the CENP-A C-terminus in likely supporting important centromeric interactions, like CENP-C, which is known to stabilize CENP-A nucleosomes (Carroll et al., 2010; Falk et al., 2015). Consistent with this, salt fractionation experiments in *Cenpa^mScarlet/+^ testes* revealed that although the tagged CENP-A-mScarlet protein incorporates into chromatin, it is less stable than the untagged CENP-A (**Figure 1SC**). Importantly, despite reduced stability *in vitro* with increasing salt concentrations, CENP-A-mScarlet retains proper centromeric localization *in vivo*, as evidenced by its enrichment at DAPI-dense chromocenters in both testes and ovarian follicles, colocalization with CREST or total CENP-A foci, and accumulation at chromosomal ends in spermatocyte spreads—consistent with the telocentric organization of murine chromosomes (Kalitsis et al., 2006) (**Figure S1D**). Importantly, this fluorescently tagged CENP-A protein escapes to waves of chromatin remodeling in spermiogenesis and embryogenesis and allows for parent-of-origin-specific visualization of CENP-A dynamics, offering a powerful tool to dissect inheritance and regulation of centromeric component during mammalian development – a level of insight that would not be possible without a tagged protein.

**Figure 1:**
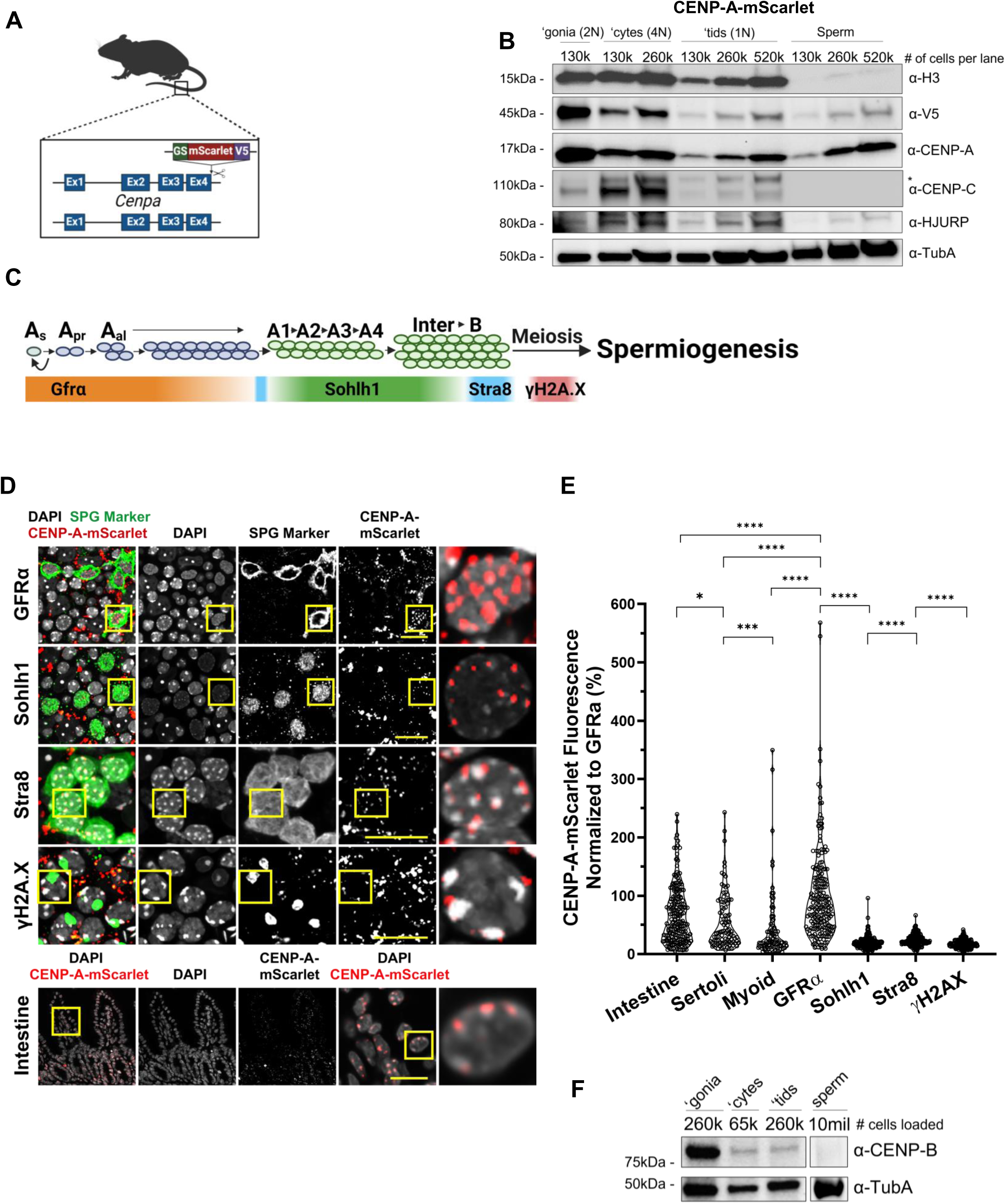
Loss of CENP-A in male germ cells precedes the histone to protamine exchange. A) Schematic of the Cenpa-GS-mScarlet-i-V5 transgene. B) Immunoblot of H3, V5, CENP-A, CENP-C, and HJURP in flow sorted spermatogonia (‘gonia), pachytene/diplotene spermatocytes (‘cytes), round spermatids (‘tids), and mature sperm from *Cenpa^mScarlet/+^* males. Alpha tubulin (TubA) is shown as a loading control. Shown is a representative immunoblot from n = 2 mice. Membrane was stripped and re-blotted for the indicated proteins. C) Schematic overview of germ cell markers across spermatogenesis. GFRα1 marks undifferentiated spermatogonia, SOHLH1 marks differentiating Type A and Intermediate spermatogonia, STRA8 marks differentiating Type B spermatogonia and early preleptotene spermatocytes, and condensed γH2A.X marks mid to late pachytene spermatocytes. D) Immunostaining of *Cenpa^mScarlet/+^* whole mount tubules in specified germ cell types. Representative images from n = 3 staining experiments on n = 3 mice. Scale bars are 20um. E) Quantification of CENP-A-mScarlet direct fluorescence across germ cell stages, somatic cells of the testes, and intestinal cells. n = 200 germ and intestinal cells were quantified for each cell type, split evenly between n = 2 staining experiments and n = 2 mice. n = 100 Sertoli and Myoid cells were quantified for each cell type. Each dot is the sum of all CENP-A-mScarlet puncta in one cell. Average fluorescence of CENP-A-mScarlet normalized to GFRα cells across the cell types are as follows: intestine = 68%, Sertoli = 56%, Myoid = 43%, GFRα = 100%, Sohlh1 = 19% Stra8 = 24%, γH2A.X = 18%. F) Immunoblots of CENP-B protein levels in flow sorted germ cells and mature sperm from C57Bl/6J testes and epididymides. Representative image from n = 2 replicates from n = 2 mice.

### Reduction of CENP-A coincides with the onset of germ line stem cell differentiation, preceding the histone-to-protamine exchange in testes

During spermiogenesis, the male epigenome undergoes a near complete replacement of canonical histones with small basic proteins called protamines (Bao and Bedford, 2016), a process highly conserved across many vertebrates and invertebrates (Lewis et al., 2003). Although retention of CENP-A nucleosomes has been reported in mammals, invertebrates, and even plants (Dunleavy et al., 2012; Milks et al., 2009; Palmer et al., 1990; Raychaudhuri et al., 2012; Schubert et al., 2014), the dynamics of CENP-A throughout both male and female gametogenesis programs remain poorly understood (reviewed in Das et al., 2020 and Štiavnická et al., 2025). To answer this question, we used flow cytometry to enrich for germ cell populations isolated from adult *Cenpa^mScarlet/+^*males. As expected, histone H3 levels are fairly constant between spermatogonia and meiotic spermatocytes but decrease in post-meiotic haploid spermatids and are largely depleted in mature spermatozoa (**Figure 1B and S1E**). In contrast, a significant fraction of tagged CENP-A-mScarlet is lost at the transition from spermatogonia to spermatocytes. Subsequently, a smaller but notable loss occurs again during the transition from spermatocytes to round spermatids, with no significant change between spermatids and mature sperm (**Figure 1B and S1E**). Importantly, similar dynamics were observed when assessing total CENP-A levels in *Cenpa^mScarlet/+^* germ cells across stages (**Figure 1B and S1E**). Overall, these data suggest that tagged and untagged CENP-A proteins behave similarly, and that the loss of CENP-A from paternal chromosomes occurs before the histone-to-protamine exchange in spermatids.

To more precisely define the onset of CENP-A loss, we performed combinatorial staining on whole mount seminiferous tubules to resolve different spermatogonia and germ cell subtypes. Within tubules, the most undifferentiated spermatogonial stem cells known as A_single_ (A_s_) undergo a series of incomplete mitotic divisions to form undifferentiated germ cell clones (**Figure 1C**), consisting of syncytial clones of 2, 4, 8, and 16 cells interconnected by cytoplasmic bridges, referred to as A_paired_ (A_pr_) or A_aligned(al)_ (A_al-4_, A_al-8_, and A_al-16)_. The A_s_, A_pr,_ and A_al_ cells are collectively referred to as A_undifferentiated_ (A_undiff_) spermatogonia, a heterogenous pool marked by two proteins: GFRα (A_s_, A_pr_, A_al-4_, and A_al-8_) and ID4 (A_s_ and A_pr_). The A_al_ differentiate into A1 spermatogonia, which undergo additional transit amplifying divisions to form A2, A3, A4, Intermediate, and Type-B spermatogonia; characterized by SOHLH1 followed by STRA8 expression (**Figure 1C**). To investigate CENP-A dynamics across early stages of germ cell differentiation and to compare CENP-A levels between germline and soma, we quantified fluorescence intensity of *Cenpa^mScarlet/+^* in intestinal villi and testis somatic cells (Sertoli and myoid) and contrasted these data to *Cenpa^mScarlet/+^* levels in various spermatogonia subtypes (GFRα1 and/or ID4) (Buageaw et al., 2005; Helsel et al., 2017), differentiating spermatogonia (SOHLH1 and STRA8) (Ballow et al., 2006; Endo et al., 2015; Zhou et al., 2008), and mid-meiotic prophase spermatocytes (γH2A.X) (Mahadevaiah et al., 2001) (**Figure 1C**). Notably, CENP-A-mScarlet fluorescence intensity was significantly higher in undifferentiated spermatogonia compared to other subtypes including differentiating germ cells, testis somatic cells (Sertoli and myoid cells) or in intestinal crypt cells, indicative of a unique increase in CENP-A density at centromeres of germline stem cells (**Figure 1D&E**). Across germ cells, a 30% decrease in CENP-A levels is immediately observed when comparing stem cells with high-vs. low-self renewal potential (i.e. ID4-high vs. ID4-Low A_s_, A_pr_, and A_al_ spermatogonia (**Figure S1F-I**) (Chan et al., 2014; Helsel et al., 2017; Sun et al., 2015). These findings indicate that the reduction in CENP-A that occurs very early in the undifferentiated stem cells potentially distinguishes spermatogonia with high vs. low stemness potential. Therefore, CENP-A reduction coincides with stem cell differentiation and precedes early chromosome remodeling known to occur during meiotic prophase and global chromatin remodeling in round spermatids.

To assess whether the observed decrease in CENP-A during murine germ cell differentiation is a unique or conserved phenomenon, we turned to *Drosophila melanogaster*, a model system in which germline stem cell hierarchies can be easily identified histologically. We collected adult fly testes and quantified the total fluorescence intensity of CID, the *Drosophila* CENP-A ortholog, in both somatic hub cells and germ cells at increasing distances from the hub, which correspond to different stages of differentiation. Consistent with our findings in the mouse germline, CID levels were elevated in germline stem cells compared to hub cells and progressively decreased during spermatogonia transit-amplification divisions (**Figure S2A-C**). This suggests that CENP-A accumulation followed by a differentiation-coupled reduction is a conserved feature of male germ cell differentiation in both mice and flies.

### Centromere constituents change throughout spermatogenesis

Next, we examined whether other centromeric proteins that directly interact with CENP-A have similar dynamics during spermatogenesis. Briefly, we found that CENP-C levels increase transiently in spermatocytes (with normalization for ploidy), despite significant loss of CENP-A, but decreases in haploid spermatids (**Figure 1B and S1E**). In contrast, CENP-B, which binds to B-box DNA sequences in minor satellites (Fachinetti et al., 2015; Fujita et al., 2015), levels were significantly reduced from spermatogonia to spermatocytes but were maintained in spermatids (**Figure 1F and S1E**). However, in mature sperm, both CENP-B and CENP-C were undetectable, consistent with earlier observations in bull and *Drosophila* sperm (Palmer et al., 1990; Raychaudhuri et al., 2012). For HJURP, the CENP-A-specific chaperone protein, we observed high levels in spermatogonia and spermatocytes which decreased in round spermatids. Unlike CENP-B and CENP-C, low levels of HJURP persisted in mature sperm (**Figure 1B and S1E**). The HJURP present in sperm appears to interact with CENP-A and H2B containing nucleosomes, as evidenced by co-immunoprecipitation (co-IPs) in mouse (**Figure S1J**) or reciprocal co-IPs from human sperm (**Figure S1K**). Taken together, these data indicate that many centromere-associated proteins (in addition to CENP-A) are actively remodeled during germ cell differentiation, and that sperm-retained HJURP either associates with or localizes near CENP-A-containing nucleosomes, suggesting that paternally inherited HJURP may play a role in zygotic development.

### CENP-A density at centromeres increases from PGCs to GV oocytes

Given the observed CENP-A dynamics in the male germline, we next investigated whether similar CENP-A dynamics occur in the female germline during oogenesis. Oogenesis in mice begins in the embryonic gonad, but these oocytes arrest at the dictyate stage of meiotic prophase I shortly after birth, which can last for the entirety of a female’s reproductive lifespan (Larose et al., 2019). Here, we co-stained ovarian tissue cross-sections from OCT4^eGFP^ mice (Jones, 2008; Larose et al., 2019) to quantify total CENP-A or CENP-C levels in primordial germ cells (PGCs; E11.5) and in differentiating oogonia (E13.5, E18.5, P2, and adult) (**Figure S3A**). As in the male germline, the early female germline progenitors, such as the PGCs and oogonia, have a higher level of CENP-A relative to the neighboring somatic cells; however, CENP-A levels at centromeres continue to increase between birth and adult germinal vesicle-intact oocytes (GVs) (with normalization for ploidy) (**Figure S3B**). At the same developmental stages, CENP-C levels followed similar patterns as CENP-A: all assayed female germ cells contained more CENP-C than neighboring somatic cells, and the major increase in CENP-C occurred between birth and adult GV (**Figure S3C**). To monitor changes after meiotic resumption, we isolated GV oocytes from *Cenpa^mScarlet/+^* and WT mice and *in vitro* matured GV oocytes (4n) to generate metaphase I (MI) (4n) and metaphase II (MII) (2n) oocytes. CENP-A levels at centromeres were highest in GV and MI but decreased slightly at the MI-to-MII transition after normalizing for DNA ploidy (**Figure S3D&E**). In contrast, CENP-C levels at centromeres steadily increased during female meiosis, when normalized for DNA ploidy (**Figure S3F**). Thus, unlike in males, centromeric CENP-A levels rise during oocyte maturation but a small proportion of CENP-A is lost from meiosis I to II. This modest decrease in CENP-A is accompanied by an increase in CENP-C levels.

### Germline centromeric remodeling is regulated in a sex-specific manner

Given the differences in CENP-A dynamics in the male and female germlines, we next sought to directly compare CENP-A levels in mature sperm and oocytes, as unequal amounts of CENP-A in gametes may pose a challenge for faithful chromosome segregation during zygotic mitosis. To quantify the extent of this difference between the two gametes, we collected and compared 450 GV and 450 MII oocytes to a titration series of sperm concentrations. After correcting for ploidy, CENP-A levels in sperm are significantly lower, maintaining only ∼17.5% of CENP-A detected in GV oocytes or ∼10.6% of CENP-A present in MII oocytes (**Figure 2A&B**). This marked difference in CENP-A levels was further confirmed by immunofluorescence quantification in MII oocytes and mature sperm (**Figure 2C**). To assess whether these differences are inherited, we collected and examined zygotes from *Cenpa^mScarlet/+^* transgenic males and females immediately post-fertilization. Consistent with gamete measurements, paternal chromosomes in these zygotes had only about 10–15% of the CENP-A-mScarlet fluorescence intensity observed on maternal chromosomes (**Figure 2D&E**). Notably, the differences in centromere composition extended beyond CENP-A, as sperm failed to retain CENP-C, while CENP-C levels on maternal chromosomes increase from GV to MII eggs roughly two-fold (183.7%) (**Figure 2A&B**). To assess whether CENP-A asymmetry is conserved across species, we collected unfertilized MII-stage human oocytes and sperm samples and performed similar immunofluorescence measurements. Strikingly, the differential CENP-A retention between oocytes and sperm was also evident in humans, with human sperm retaining ∼10% of the CENP-A present in human oocytes (**Figure 2F**). We also extended our analysis to *Drosophila melanogaster*, where we quantified endogenous CID-Dendra2 (the fly homolog of CENP-A) fluorescence levels in oocytes and sperm. Although sperm in flies retained higher levels of CID-Dendra2 relative to oocytes (∼60% of oocyte levels), this asymmetry remained significant (**Figure 2G&H**). The less pronounced difference in *Drosophila* gametes likely reflects the known post-meiotic deposition of CENP-A in males (Dunleavy et al., 2012; Raychaudhuri et al., 2012). Collectively, these findings highlight a conserved, yet species-tuned, difference in centromere composition between male and female gametes.

**Figure 2:**
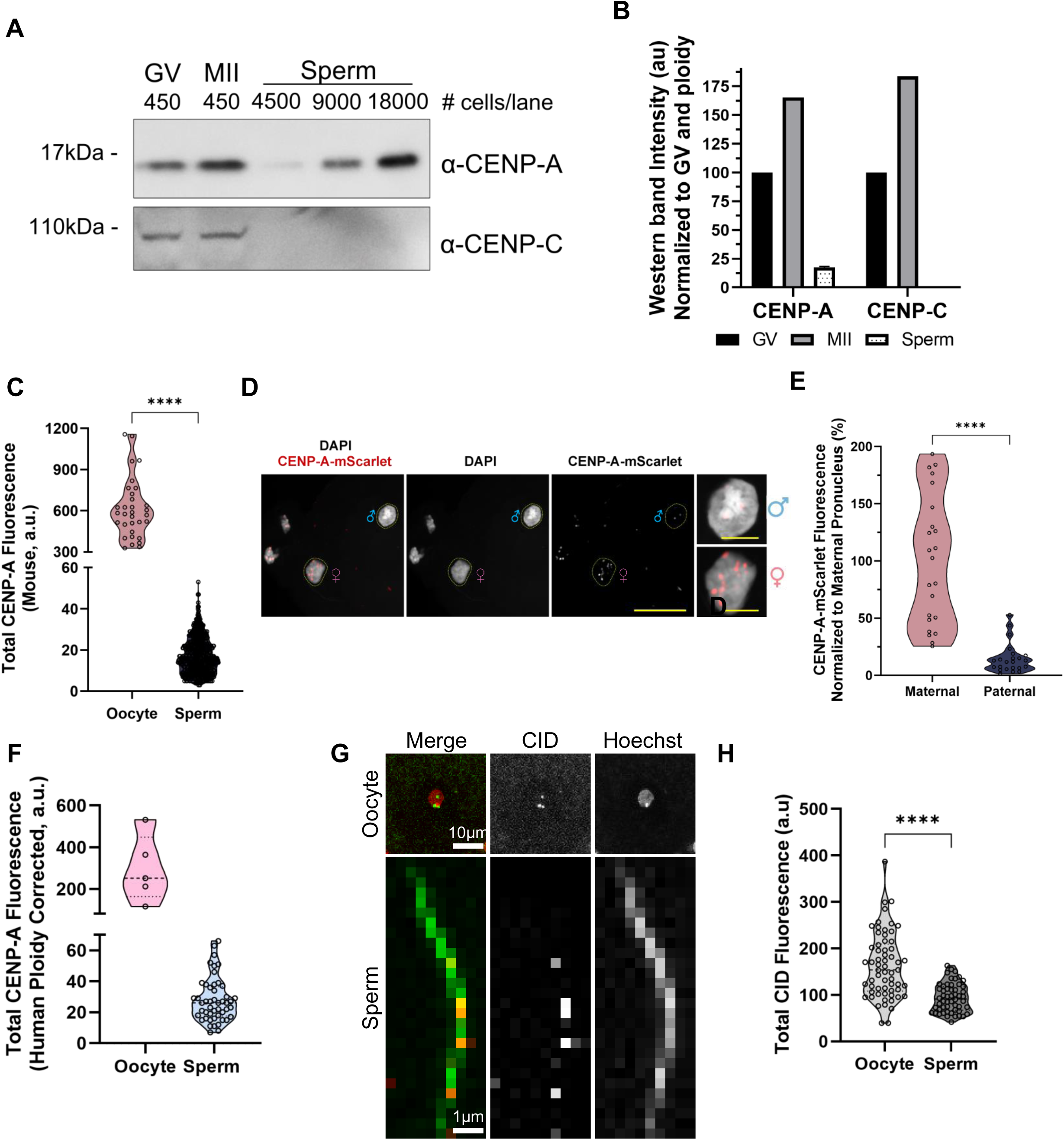
CENP-A asymmetry between oocytes and sperm is a conserved feature of gametogenesis across flies, mice, and humans. A) Immunoblot analysis of CENP-A and CENP-C in GV, MII and increasing sperm concentrations per lane. The membrane was stripped and re-probed with the indicated antibodies. Representative image from n = 2 blots using oocytes from n = 12 females and n = 2 males. B) Quantification of western band intensities from (A). Data is shown normalized to ploidy, cellular input, and GV protein levels. Mean values are as follows: GV CENP-A = 100, MII CENP-A = 165.26, Sperm CENP-A = 17.5, GV CENP-C = 100, MII CENP-C = 183.7, Sperm CENP-C = 0. C) Quantification of total CENP-A immunofluorescence in MII oocytes and mature sperm from C57BL/6J mice. Data shown is from n = 6 females and n = 2 males and are normalized for ploidy. Mean values are as follows: Oocyte = 611 a.u. for n = 32 cells and sperm = 17 a.u. for n = 600 cells. D) Representative image of CENP-A-mScarlet fluorescence inherited from *Cenpa^mScarlet/+^* males and females. Data shown is from n = 23 zygotes from n = 2 technical replicates. D) Quantification of endogenous CENP-A-mScarlet fluorescence in the maternal and paternal pronuclei of zygotes shown in panel (D). Average CENP-A-mScarlet fluorescence, normalized to the maternal average, is as follows: Maternal = 100%, Paternal = 15%. F) Quantification of total CENP-A immunofluorescence in human oocytes and sperm. Mean a.u. values are as follows: Oocyte = 295 a.u. for n = 5 cells and sperm = 28 a.u. for n = 49 cells. Data are from n = 3 female donors and n = 3 male donors and are normalized for ploidy. G) Representative images of *Drosophila* oocyte and sperm with endogenous CID-Dendra2 fluorescence (red), co-stained with Hoechst (green). Scale bars: oocyte = 10um, sperm = 1um. H) Quantification of CID-Dendra2 fluorescence from *Drosophila* gametes shown in (G). Mean a.u. values, normalized for ploidy, are as follows: Oocytes = 163 a.u. for n = 62 and sperm = 94.2 a.u. for n = 94. ****: p < 0.0001.

### Paternal CENP-A-containing nucleosomes are inherited intergenerationally in mouse embryos

To investigate the fate of CENP-A nucleosomes in mouse sperm, we performed *in vitro* fertilization (IVF) using WT females and *Cenpa^mScarlet/+^*transgenic males, collecting embryos at various stages of early embryonic development. (**Figure 3A**). Progression through the first embryonic cell cycle can be grouped into five pronuclear stages (PN1 to PN5) based on the size and relative distance between the maternal and paternal pronuclei (Adenot et al., 1997). Following fertilization, we visualized paternal CENP-A-mScarlet nucleosomes in the decondensing sperm head of PN0 stage embryos, throughout all five pronuclear stages, and in the blastomeres of 2-cell embryos after the first mitotic division (**Figure 3B**). Based on the presence of the tagged protein in 2-cell embryos, we conclude that paternal CENP-A nucleosomes survive the protamine-to-histone exchange, DNA replication, and global genome reorganization, contributing to early embryonic chromatin. To our knowledge, this is the first direct evidence of paternal nucleosome inheritance into early mouse embryos.

**Figure 3:**
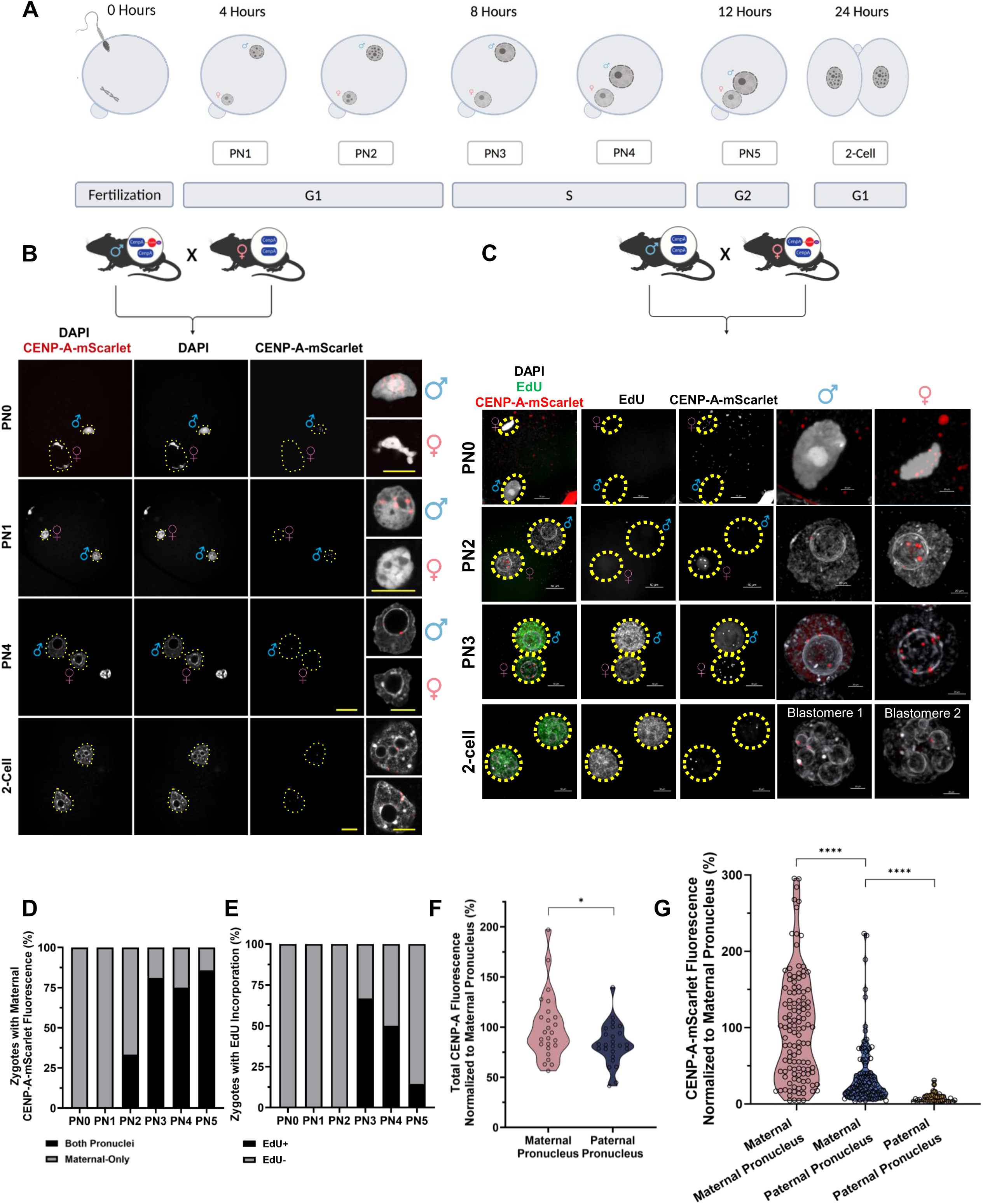
Maternal– and paternal-derived CENP-A nucleosomes are inherited intergenerationally. A) Overview of pronuclear staging during the first cell cycle post-fertilization. PN = pronuclear stage. B) Visualization of CENP-A-mScarlet fluorescence in *in vitro* fertilized embryos, generated using C57Bl/6J females and *Cenpa mScarlet/+* males, collected at the indicated pronuclear stages. Representative images from four IVF and immunostaining experiments using four males. Maternal and paternal pronuclei were identified based on relative size and position in relation to the polar body. Scale bars: 20 μm (main) and 10 μm (insets).C) Imaging of endogenous maternal CENP-A-mScarlet fluorescence at the indicated pronuclear stages. Representative images from four IVF and immunostaining experiments using 12 females. Maternal and paternal pronuclei were identified by their relative size and position in relation to the polar body. EdU, visualized using click chemistry, marked zygotes that had entered S phase. Scale bars: 20 μm (main) and 10 μm (insets). D) Quantification of the percentage of zygotes with maternal CENP-A-mScarlet in both pronuclei or only in the maternal pronuclei at each pronuclear stage. E) Quantification of the percentage of zygotes positive or negative for EdU incorporation (indicating DNA replication). F) Quantification of total CENP-A immunofluorescence in the maternal and paternal pronuclei at stage PN5, each dot is the sum of total CENP-A puncta in a single pronucleus. Fluorescence values were normalized to the mean for the maternal pronucleus. Mean total CENP-A fluorescence in the maternal pronucleus = 100% (n = 25 embryos). Mean total CENP-A fluorescence in the paternal pronucleus = 83% (n = 25 embryos). p < 0.05. G) Comparison of direct fluorescence intensity of maternally vs. paternally derived CENP-A-mScarlet in male and female pronuclei at 8 hours post-fertilization. Only zygotes with maternal CENP-A-mScarlet present in both pronuclei were included. Each dot represents the total CENP-A-mScarlet puncta in a single pronucleus. Values were averaged to the mean fluorescence quantified for maternally inherited mScarlet in the maternal pronucleus. Mean for maternal CENP-A-mScarlet in the maternal pronucleus = 100% from n = 119 embryos. Mean for maternal CENP-A-mScarlet in the paternal pronucleus = 37% from n = 103 embryos. Mean for paternal CENP-A-mScarlet in the paternal pronucleus = 8% from n = 35 embryos. *: p < 0.05, ****: p < 0.0001.

### Parental centromeres begin to equalize prior to DNA replication in zygotes

To track the dynamics of maternally inherited CENP-A in zygotes, we next performed IVF using *Cenpa^mScarlet/+^*oocytes and WT sperm. As expected, maternal CENP-A-mScarlet nucleosomes were visible on MII chromosomes completing meiosis II just post-fertilization (**Figure 3C**). Furthermore, we observed the tagged CENP-A-mScarlet histones throughout all pronuclear stages as well as in the blastomeres of 2-cell embryos (**Figure 3C**). Importantly, these experiments show that maternally inherited CENP-A begins localizing to paternal centromeres prior to the onset of zygotic S-phase (PN2/3; **Figure 3C-E**). By PN3 stage, most zygotes (77%) have maternal CENP-A-mScarlet in the paternal pronucleus, while a smaller fraction (23%) showed localization only in the maternal pronucleus (**Figure 3D**). In late-stage zygotes (PN5), total CENP-A levels were nearly equal between the parental pronuclei, with a paternal/maternal ratio of 0.87 (**Figure 3F**). Moreover, quantification of mScarlet fluorescence in PN3-5 zygotes revealed that the majority of CENP-A present in the paternal pronucleus is maternally derived (**Figure 3G**). These results demonstrate that CENP-A equalization is largely achieved prior to the first cell division.

### Equalization of parental centromere density is independent of zygotic transcription, translation, and DNA replication

Oocytes contain a significant reservoir of mRNA and protein that supports embryonic development prior to zygotic genome activation (ZGA) but are not believed to carry a large pool of free cytosolic CENP-A protein (Smoak et al., 2016). To examine whether minor zygotic genome activation (ZGA) is necessary for deposition of maternal CENP-A-mScarlet onto paternal centromeres we collected MII eggs from superovulated *Cenpa^mScarlet/+^*females and performed IVF using mature sperm from WT male mice. At 1.5 hours after fertilization, zygotes were treated with 5,6-dichloro-1-β-D-ribofuranosyl-benzimidazole (DRB), a reversible RNA polymerase II inhibitor shown to inhibit minor ZGA in mice (Abe et al., 2018), or vehicle control (DMSO). To confirm the efficiency of our DRB treatment, we incubated embryos with 5-ethynyl-uridine (EU), a uridine analog that can be detected using click chemistry reactions (Jao and Salic, 2008), to monitor newly synthesized transcripts (**Figure S4A&D**). As expected, the DRB treatment inhibited transcription in zygotes, as determined by the difference in total EU signal in vehicle and DRB-treated embryos (**Figure S4A**, note that EU has cytoplasmic staining in all assayed cell types and treatments). Importantly, inhibition of the minor ZGA did not alter cell cycle progression (**Figure S4G**) nor did it prevent maternal CENP-A-mScarlet from accumulating in the paternal pronucleus. Consequently, no significant differences in CENP-A-mScarlet intensity were observed between the DRB-treated maternal and paternal pronuclei relative to the controls (**Figure S4A&D**). Altogether, these results indicate that neither zygotic transcription of *Cenpa* or transcription-mediated nucleosome turnover are necessary for maternal CENP-A-mScarlet to be localized to the paternal genome.

Next, we asked whether zygotic translation is required for the increase in CENP-A levels observed in the paternal pronucleus. To test this, we treated IVF-derived embryos containing maternal CENP-A-mScarlet with either cycloheximide (CHX) to inhibit translation or an equal volume of vehicle (DMSO) after 1 hour of fertilization and cultured the zygotes for approximately another seven hours. To verify that the CHX treatment was effective, we added O-propargyl-puromycin (OPP), which incorporates into the nascently synthesized polypeptide chains to monitor translation activity by click chemistry (Liu et al., 2012). As confirmation that CHX successfully inhibited translation, the OPP signal was absent in CHX-treated zygotes (**Figure S4B**). Interestingly, inhibition of protein synthesis did not significantly change the amount of maternal CENP-A-mScarlet levels being deposited into the paternal pronucleus, although there was a statistically significant decrease in the maternal pronucleus (**Figure S4B&E**). To eliminate potential bias from cell cycle differences (**Figure S4H**), we re-analyzed only PN3 embryos and still did not detect any significant reduction of CENP-A-mScarlet among male pronuclei. However, we again found a slight, but statistically significant decrease in the CENP-A-mScarlet fluorescence in maternal pronuclei of CHX-treated embryos (**Figure S4J**). These results suggest that newly made CENP-A protein in zygotes maybe be needed to achieve optimal steady-state CENP-A levels at least in the maternal pronucleus.

Finally, because most zygotes incorporate CENP-A nucleosomes near the time of S-phase, we assessed whether DNA replication contributes to or is necessary for the acquisition of maternal CENP-A-mScarlet into the paternal genome. To this end, we followed a similar IVF schema, but incubated zygotes with aphidicolin (Aph), a reversible DNA polymerase inhibitor that blocks DNA synthesis (Ikegami et al., 1978) or vehicle (DMSO), starting 1.5 hours after fertilization, and assessed incorporation of the thymidine analog 5-Ethynyl-2′-deoxyuridine (EdU) to confirm DNA replication inhibition (**Figure S4C**). Again, inhibition of zygotic DNA replication did not alter maternal CENP-A-mScarlet levels in the paternal pronucleus or drastically change the pronuclear staging of Aph-treated embryos (**Figure S3C, F, I**). Taken together, these experiments suggest that paternal incorporation of maternal CENP-A-mScarlet does not require embryonic transcription, translation, or DNA replication, although some contribution from *de novo* translation is possible.

### CENP-A equalization relies on maternally inherited CENP-A

Since zygotic transcription and translation contributed very little to the equalization of maternal/paternal centromeres, we next examined whether pre-existing, chromatin-bound maternal CENP-A nucleosomes and/or an inherited cytoplasmic CENP-A protein pool contributes to equalization. To test this possibility, we collected *Cenpa^mScarlet/+^* GV oocytes, photobleached the mScarlet fluorescence in the nuclei, and performed *in vitro* maturation from GV to metaphase II (MII) oocytes. The *in vitro* matured MII oocytes were then fertilized with WT sperm, and the resulting zygotes were used to monitor the levels and dynamics of CENP-A-mScarlet and total CENP-A (**Figure S5A**). Our photobleaching protocol depleted ∼80-90% of the CENP-A-mScarlet signal in the GV nuclei while maintaining oocyte integrity and competency (**Figure S5 B & C; see methods**). Furthermore, we did not observe significant photorecovery after holding GV oocytes for either one or 24 hours, nor following their maturation through the remainder of meiosis (**Figure S5 B & C**). This suggests that oocytes arrested in prophase I and progressing through meiosis exhibit minimal to no exchange of chromatin-bound CENP-A. This observation is consistent with previous work suggesting that CENP-A nucleosomes assembled on oocyte chromosomes are extremely stable even in the absence of nascent *Cenpa* transcription or deposition (Das et al., 2023; Smoak et al., 2016).

Given these observations, we next performed IVF using photobleached or unbleached *in vitro* matured oocytes. The resulting zygotes were cultured in the presence of CHX and were collected eight hours after fertilization. To verify that photobleaching did not interfere with CENP-A equalization, we compared total CENP-A levels between photobleached and control embryos using immunofluorescence. Importantly, no significant differences were found between the groups (**Figure S5D&E**), suggesting normal equalization in embryos from photobleached oocytes. As expected, CENP-A-mScarlet levels were significantly reduced in the maternal pronuclei of photobleached embryos compared to controls (32% of WT levels**; Figure S5D&F**). Surprisingly, despite near complete photobleaching (reducing by about 90% compared to WT levels) in GV and MII oocytes, we detected a notable amount of new CENP-A-mScarlet in the maternal pronucleus. This suggests that new CENP-A is incorporated, potentially due to CENP-A turnover in the zygote or a need to accumulate more CENP-A in zygotes. In contrast, the paternal pronucleus accumulates ∼50% of CENP-A-mScarlet levels observed in WT embryos, suggesting that CENP-A deposition in the paternal genome may depend on two sources: recycled maternal CENP-A and a free inherited cytoplasmic pool of CENP-A from oocytes.

To confirm the possibility of inheriting a cytoplasmic pool of CENP-A RNA/protein, we performed RT-qPCR across various developmental stages, and confirmed the presence of Cenpa RNAs in GV, MII, zygotes, 2-cell, and 4-cell embryos. We observed that CENP-A RNA levels decreased from GV oocytes to zygotes but peaked after zygotic genome activation (**Figure S5G**). Additionally, re-analysis of low-input ribosome profiling data (Ribo-lite) from GV, MI, MII oocytes, and multiple embryonic stages, confirmed that CENP-A mRNA is actively engaged with ribosomes during GV-MII and at multiple embryonic stages, suggesting that a cytoplasmic pool of CENP-A protein is synthesized during oocyte maturation and supplied to the embryo (**Figure S5H**) (Xiong et al., 2022). Taken together, our data suggests that increasing CENP-A levels at both parental centromeres likely relies on maternally provided CENP-A protein.

### CENP-A accumulation in zygotes does not rely on sensing CENP-A asymmetry or the presence of maternal pronucleus

Our photobleaching experiments suggested that CENP-A nucleosomes at paternal centromeres likely rely on two sources: a cytoplasmic pool and a recycled maternal nuclear pool. To examine whether the recycled maternal nuclear pool is “special” or is necessary for initiating CENP-A accumulation at paternal centromeres, we generated diploid zygotes using intracytoplasmic sperm injection (ICSI), as well as diploid androgenetic zygotes, which were created by injecting two sperm nuclei into enucleated oocytes (**Figure 4A**). In the ICSI embryos, despite using a different mouse strain for the oocytes, the CENP-A levels at maternal and paternal centromeres equalize, similar to what we observed in IVF-generated zygotes. Interestingly, in the androgenetic zygotes we find that the total CENP-A levels at paternal centromeres reach nearly identical levels to those in the paternal genomes of ICSI-derived embryos (**Figure 4B**). These findings emphasize several important points: First, recycling of a maternal nuclear CENP-A pool is not essential for paternal centromere loading or for directing CENP-A equalization, as the maternal cytoplasm alone is sufficient to support this process in the absence of a maternal pronucleus. Second, CENP-A equalization does not require sensing parental asymmetry, as zygotes containing two sperm-derived pronuclei—each with equally low CENP-A—nonetheless initiate the CENP-A deposition cascade and reach levels comparable to those of a normal paternal pronucleus. Finally, since both sperm nuclei accumulate equivalent levels of CENP-A, this suggests that the oocyte provides a non-limiting of CENP-A protein, but the amount of CENP-A that gets incorporated is dictated by intrinsic properties of the sperm-derived centromeric chromatin, rather than being instructed by the maternally inherited centromeric levels.

**Figure 4:**
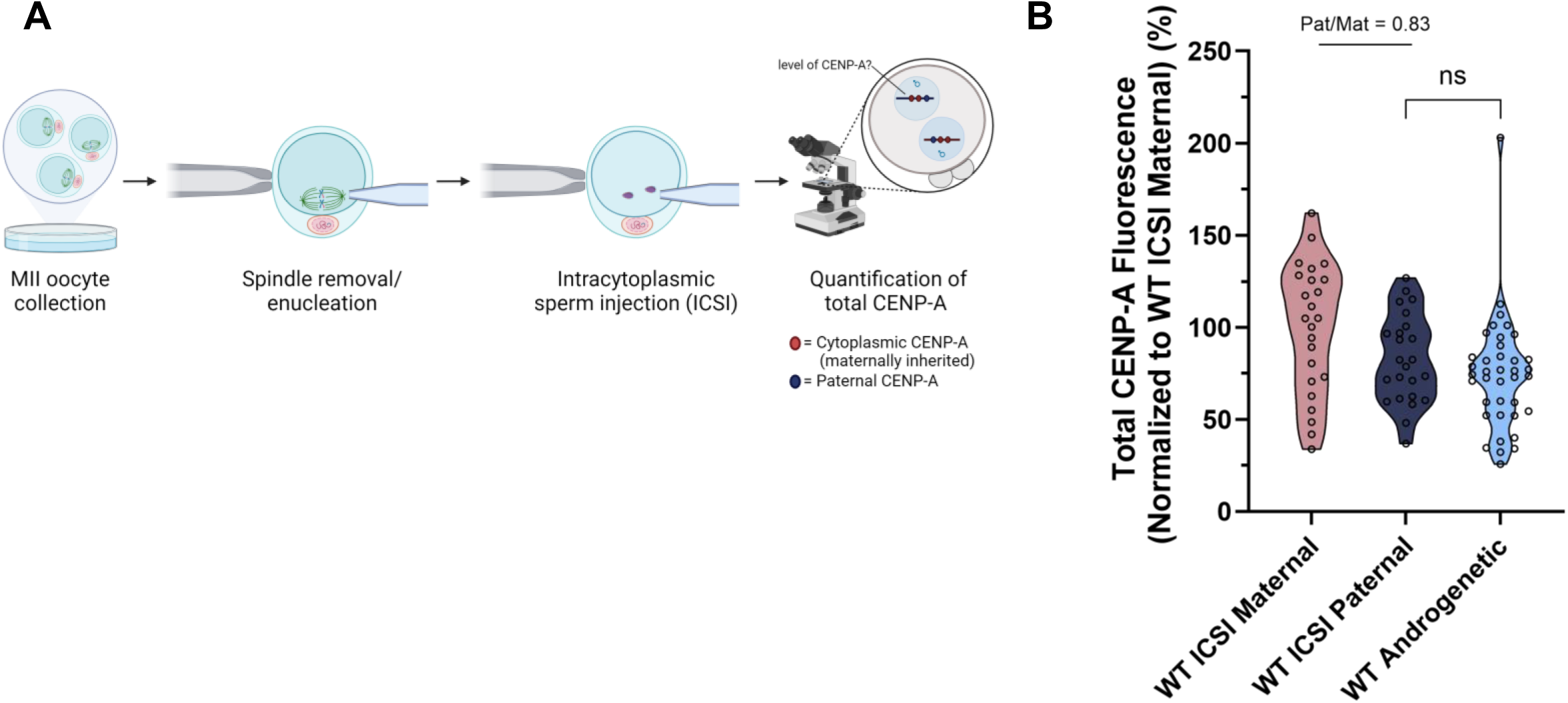
CENP-A incorporation in the paternal pronucleus is autonomously regulated. A) Schematic of androgenetic (diploid zygotes with only paternal DNA) embryo generation. B) Quantification of total CENP-A immunofluorescence in either control or androgenetic zygotes collected 16 hours after ICSI. Embryos were collected and stained from n = 2 independent ICSI experiments. Each dot is the sum of the total puncta in one pronucleus. For androgenetic embryos in which the genomes fused into one pronucleus, the total fluorescence was halved to estimate the total per genome. Values were normalized to the mean for the control maternal pronucleus. Mean fluorescence intensities are as follows: WT ICSI Maternal = 100% from n = 24 pronuclei, WT ICSI Paternal = 83% from n = 24 pronuclei, and WT Androgenetic = 76% from n = 38. ns is not significant.

### CENP-A redistribution in the zygote is licensed by the somatic cell cycle regulators CDK1/2 and PLK1

In cultured human cells, the timing and duration of CENP-A deposition is regulated by two kinases: PLK1 and CDK1. Consistently, small molecule inhibition of CDK1/2 induces ectopic CENP-A deposition during the S and G2 phases, while PLK1 inhibition in mitotic cells prevents CENP-A deposition regardless of CDK activity (McKinley and Cheeseman, 2014; Müller et al., 2014; Pan et al., 2017; Silva et al., 2012; Spiller et al., 2017; Stankovic et al., 2017). To examine the role of these kinases in zygotic centromere remodeling, we tested various inhibitor concentrations in zygotes and found that 5uM of Flavopiridol (CDK1/2 inhibitor) and 10nM BI2536 (PLK1 inhibitor) were sufficient to allow >80% of treated zygotes to develop to the 2-cell stage (data not shown) (Baran et al., 2013; Oqani et al., 2011). CDK inhibition (Flavo-treated) slowed zygotic cell cycle progression in embryos and increased the fraction of PN2 zygotes with CENP-A-mScarlet deposited in both pronuclei (**Figure S6E&F**). Notably, while CDKi did not change the total CENP-A-mScarlet levels in the maternal pronucleus, it increased the total CENP-A-mScarlet levels in the paternal pronucleus (**Figure S6A&C**). These findings suggest that the CDK1/2 activity can modulate the amount of CENP-A deposition in the paternal pronucleus of the zygote. In contrast, PLK1 inhibition in zygotes significantly reduced CENP-A-mScarlet levels in both pronuclei (**Figure S6B&D**), delayed cell cycle progression (**Figure S6G**), and reduced the proportion of zygotes with maternal CENP-A in both pronuclei (**Figure S6H**). In summary, our data suggest that CDK1/2 and PLK1 activity, which normally suppress premature or ectopic CENP-A loading in somatic cells, also regulate CENP-A deposition and levels in zygotes.

### CENP-C and MIS18BP1 are asymmetrically recruited to paternal centromeres in zygotes

Next, to investigate the mechanism regulating CENP-A equalization at paternal centromeres, we examined how the dynamics of known mitotic assembly factors in zygotes. In mitotic mammalian cells, CENP-C associates with CENP-A throughout the cell cycle and directs the MIS18 complex and HJURP to restore CENP-A levels in the subsequent G1 after DNA replication, ensuring the quantitative maintenance of centromeric CENP-A (Falk et al., 2015; Guo et al., 2017; Mitra et al., 2020). However, in zygotes, paternal centromeres inherit only ∼10% of the CENP-A levels present at maternal centromeres (**Figure 2**), requiring a ∼9 –10-fold increase to achieve parity— which challenges the classic one-to-one centromere inheritance model described in mitotic cells. To examine how this equalization might be achieved, we tested whether inner kinetochore components or CENP-A licensing /deposition machinery disproportionately accumulate at paternal vs. maternal centromeres. Two key inner CCAN proteins, CENP-B and CENP-C, are absent in sperm (**Figure 1B and S1E**) but are immediately deposited onto the decondensing sperm genome in PN0-stage embryos, preceding CENP-A deposition (**Figure 5A**). Further, quantification of CENP-B-EGFP (via RNA microinjection; see **methods**) and CENP-C immunofluorescence enrichment in the maternal and paternal pronuclei reveals striking differences. For CENP-B-EGFP zygotes, the amount of CENP-B recruited was comparable in both pronuclei, which is expected given that CENP-B is recruited in a DNA-dependent mechanism (**Figure 5B**). However, when examining CENP-C levels in zygotes we find that paternal centromeres load more CENP-C than maternal centromeres; this bias increases in later-stage zygotes, suggesting that asymmetric recruitment of CENP-C to paternal centromeres may contribute to the asymmetric increase in CENP-A molecules at male centromeres (**Figure 5B&C**). In addition to the increased CENP-C enrichment in the paternal pronucleus, we found that MIS18BP1, a centromere licensing factor that plays a crucial role in the deposition of CENP-A, is preferentially enriched in the paternal pronucleus of zygotes, with this asymmetry again increasing in later-stage zygotes (**Figure 5D&E**). This preferential accumulation of both CENP-C and MIS18BP1 in the paternal pronucleus may contribute to CENP-A equalization at parental centromeres.

**Figure 5:**
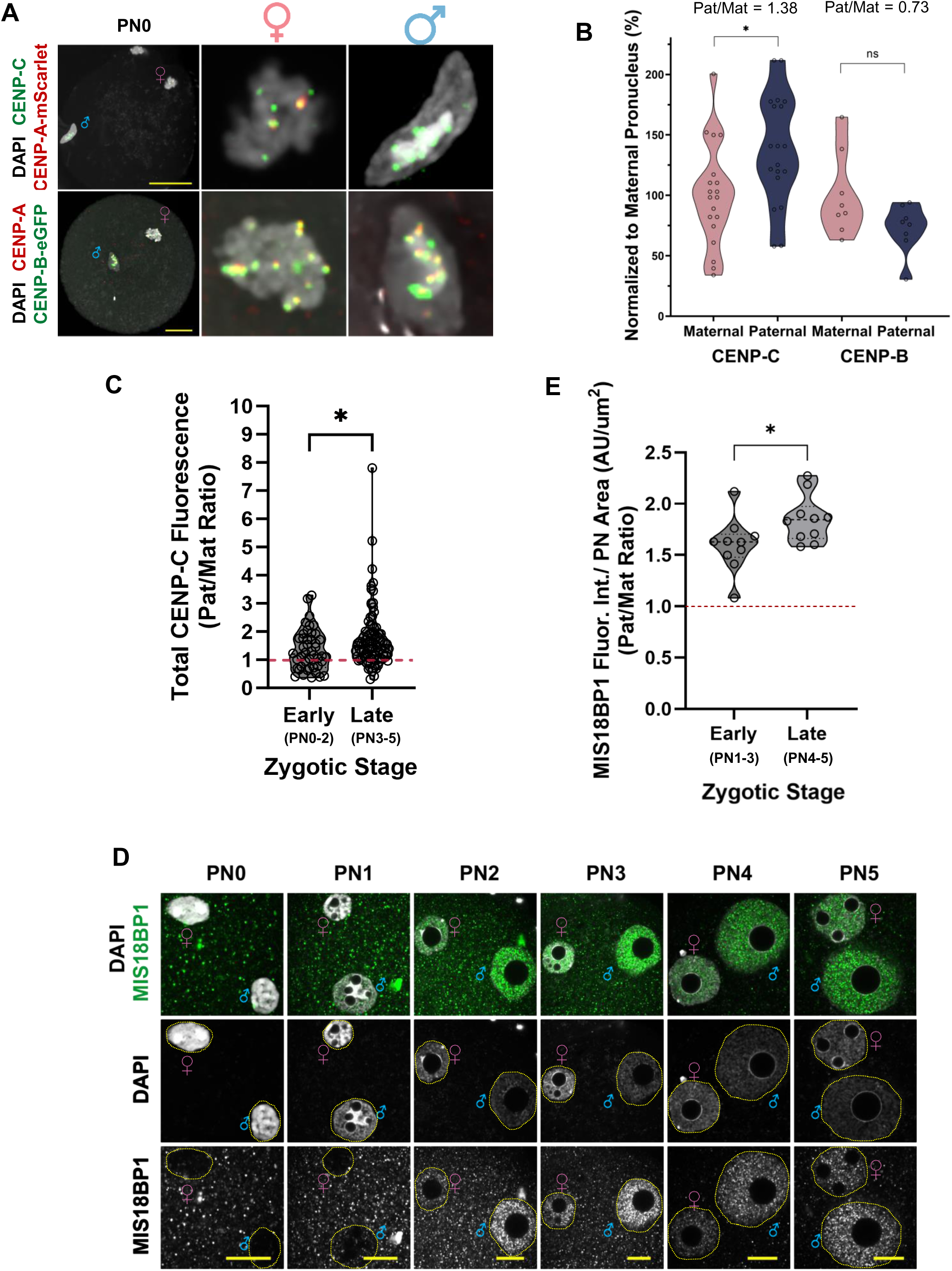
CENP-C and MIS18BP1 are asymmetrically recruited to paternal centromere in zygotes. A) Representative immunofluorescence images of PN0–2 stage zygotes co-stained for CENP-C and maternal CENP-A-mScarlet (top) or total CENP-A and CENP-B-eGFP (bottom). CENP-B-eGFP was expressed by RNA injection due to the lack of a suitable antibody. Images are representative of two IVF experiments using six *Cenpa^mScarlet/+^* or three CF-1 females and one (C57Bl/6J × DBA2)F1 male. Scale bars: 20 µm. B) Quantification of CENP-C immunofluorescence or CENP-B-eGFP fluorescence shown in panel (A), grouped by pronuclei. Each dot is the sum of the total puncta in one pronucleus. Values were normalized to the mean for the maternal pronucleus. Mean fluorescence intensities are as follows: Maternal CENP-C = 100% from n = 34 pronuclei, Paternal CENP-C = 138% from n = 34 pronuclei, Maternal CENP-B = 100% from n = 8 pronuclei, Paternal CENP-B = 73% from n = 8 pronuclei. C) Paternal/maternal ratios of total CENP-C fluorescence in early (PN0–2, *n* = 44) and late (PN3–5, *n* = 90) zygotes. Each dot represents one zygote. The red dashed line represents a paternal/maternal ratio of 1. D) Representative images of MIS18BP1 localization across pronuclear stages, from three IVF experiments using seven females. Scale bars: 20 µm. E) Quantification of MIS18BP1 fluorescence in early (PN 1-3) and late (PN 4-5) zygotes from panel (D). Fluorescence was measured from a central z-slices and normalized to pronuclear area. Shown are paternal/maternal ratios; each dot = one zygote (*n_early_* = 10, *n_late_* = 10). Red dashed line = ratio of 1. *p* < 0.05; ns = not significant.

### CENP-C directs CENP-A equalization and is critical for chromosome segregation during zygotic mitoses

Given the preferential accumulation of CENP-C in male pronuclei, we next tested the importance of CENP-C levels in mediating CENP-A equalization. To this end, we knocked down maternally inherited CENP-C by microinjecting GV oocytes with a pool of *Cenpc* mRNA-targeting siRNAs. As controls, we also monitored non-injected oocytes, as well as oocytes microinjected with a non-targeting siRNA pool (Neg Ctrl). Considering CENP-C’s role in mediating interactions with other kinetochore components during chromosome segregation (Kwon et al., 2007, McKinley et al., 2015; McKinley and Cheeseman, 2016), we titrated the concentration of the siRNA pool to achieve a knockdown efficiency of about 50% (measured by RT-qPCR) to allow the injected oocytes to progress successfully through the remainder of meiosis and fertilization (**Figure 6A**). Interestingly, reducing CENP-C levels in oocytes preferentially impaired CENP-C’s recruitment to the paternal centromeres (**Figure 6B**), and decreased the paternal-to-maternal ratio of CENP-A compared to control zygotes (**Figure 6C-F**). Next, to assess how the reduced maternal CENP-C levels impact the first zygotic division, we allowed *Cenpc* siRNA-injected oocytes to develop to the 2-cell stage. While the knockdown embryos progressed to the 2-cell stage at rates comparable to controls (**Figure 6H**), they had significantly higher incidence of chromosome mis-segregation rates (**Figure 6G&I**). Together, these findings suggest that centromere equalization and faithful chromosome segregation are highly sensitive to CENP-C dosage. This is consistent with prior studies demonstrating a CENP-C–dependent mechanism for CENP-A deposition (Carty et al., 2021; Moree et al., 2011, McKinley and Cheeseman, 2016).

**Figure 6:**
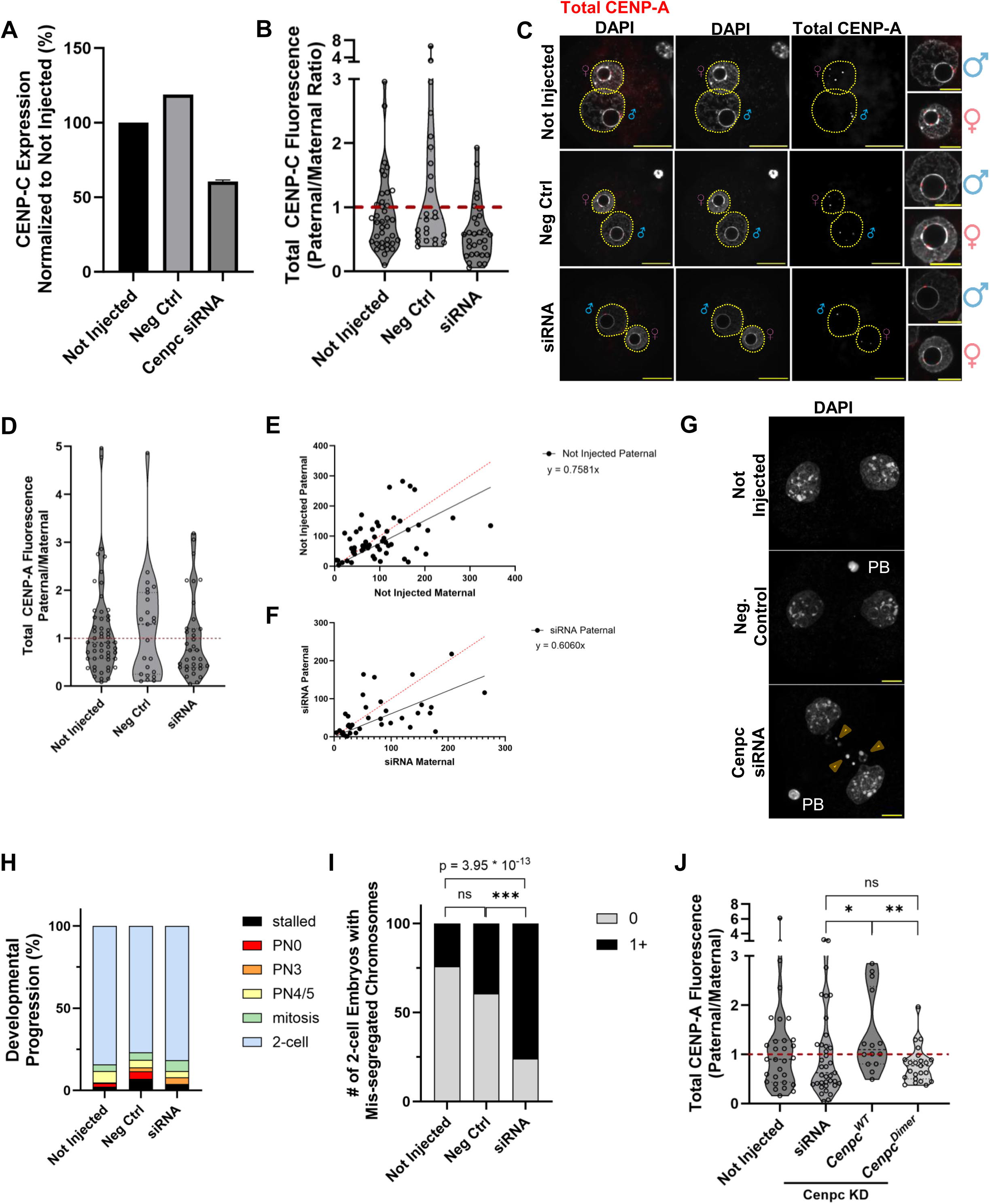
CENP-A equalization in zygotes relies on maternal CENP-C and disruption to its dimerization domain impairs and compromises chromosome segregation fidelity. A) RT-qPCR analysis confirms efficient CENP-C knockdown in meiotic oocytes. Cenpc transcript levels were quantified from uninjected controls, negative control siRNA (20–40 nM), and Cenpc siRNA (20–40 nM) groups. Each sample comprised ∼20 oocytes per n = 2 biological replicates with n = 3 technical replicates. B) Quantification of total CENP-C immunofluorescence in early (PN0–2) zygotes, shown as paternal/maternal fluorescence ratios. The red dashed line denotes a ratio of 1. Data are from n = 3 biological replicates (3 CF-1 females and 1 (C57Bl/6J × DBA2)F1 male per replicate). C) Representative images of CENP-A immunofluorescence in PN4/5 zygotes generate from GV oocytes injected with *Cenpc*, control siRNA pools, or uninjected controls. Data are from n = 6 IVF experiments using n = 3-4 CF-1 females and n = 1 (C57Bl/6J X DBA2)F1 male for each experiment. Zygotes with paternal/maternal total CENP-A ratios above 5 were excluded from further analysis. D) Quantification of CENP-A fluorescence in zygotes from (C), plotted as paternal/maternal ratios. Each dot represents one zygote. Data: uninjected (n = 54), negative control (n = 23), and siRNA-treated (n = 35) zygotes from n = 4 biological replicates. E-F) Scatterplots of CENP-A fluorescence in maternal (x-axis) vs. paternal (y-axis) pronuclei from (D) for uninjected (E) and Cenpc siRNA-treated (F) zygotes. Linear regressions were performed with y-intercepts fixed at 0. Each dot = 1 zygote. G) Representative DAPI images of 2-cell embryos from conditions in (C), cultured for 24 hours post-fertilization (hpf). H) The proportion of embryos from (G) collected at each stage shown in the legend 24hpf. “Stalled” indicates zygotes that failed to form a proper paternal pronucleus. Percentages were calculated by combining all embryos generated across all replicates. Data is representative of n = 227 not injected embryos, n = 43 neg ctrl embryos, and n = 76 siRNA embryos, from at least n = 3 replicates using n = 5 CF-1 females and n = 1 (C57Bl/6J X DBA2)F1 male each. I) The percentage of 2-cell embryos with at least one chromosome segregation defects from (G). Percentages were calculated by combining all embryos generated across all replicates. The number of 2-cell embryos are as follows: Not Injected = 191, neg ctrl = 33, siRNA = 62. J) CENP-A levels are restored in Cenpc siRNA-injected zygotes when co-injected with an siRNA resistant *Cenpc^WT^* RNA but not a CENPC dimerization-deficient RNA *(Cenpc^Dimer^*). Paternal/maternal CENP-A ratios are shown for each group: uninjected (n = 29), siRNA (n = 37), CenpcWT (n = 14), and CenpcDimer (n = 23). Data from n = 3 replicates using 5 CF-1 females and 1 (C57Bl/6J × DBA2)F1 male per replicate. The red dashed line indicates a ratio of 1. *p < 0.05, **p < 0.01, ***p < 0.001; ns = not significant.

This requirement for CENP-C raised a new question: how does the paternal genome, despite having less CENP-A at fertilization, accumulate disproportionately higher levels of CENP-C? This is especially intriguing given that CENP-C is typically recruited in a CENP-A–dependent manner (Carty et al., 2021; Moree et al., 2011, McKinley and Cheeseman, 2016). Prior studies have shown that CENP-C can dimerize or oligomerize, and mutations in residues important for dimerization impair CENP-C’s centromeric localization as well as its binding to HJURP (Carroll et al., 2010; Hara et al., 2023; Flores Servin et al., 2023). To test whether CENP-C dimerization contributes to CENP-A equalization, we again knocked down maternal CENP-C in oocytes, but co-injected siRNA-resistant *Cenpc* mRNA encoding either the wildtype protein (*Cenpc^WT^*) or a dimerization-deficient mutant mRNA (*Cenpc^Dimer^*; Carroll et al., 2010; Flores Servin et al., 2023). Zygotes were cultured to the PN4/5 stage and assessed for CENP-A equalization. Notably, zygotes injected with the *Cenpc^Dimer^* RNA failed to equalize CENP-A levels, phenocopying the siRNA-only condition, whereas *Cenpc^WT^* RNA rescued the CENP-A equalization defect observed in *Cenpc* siRNA-injected zygotes (**Figure 6J**). This finding indicates that CENP-C dimerization is required for effective CENP-A equalization.

Next, to examine whether reduced CENP-A levels in oocytes can similarly impair CENP-A equalization, we depleted the maternally inherited *Cenpa* RNA pool to approximately 20% of control levels (**Figure 7A**). This led to a global reduction in CENP-A at both parental centromeres (**Figure 7B&C**), yet equalization between pronuclei remained intact; suggesting that CENP-A equalization in zygotes relies on CENP-C levels. Curiously, in the *Cenpa* siRNA treated embryos, CENP-C levels were elevated in both pronuclei suggesting that additional CENP-C recruitment helps stabilize weaker centromeres in zygotes (**Figure 7D**). Zygotes from *Cenpa^+/-^* females, which have comparably reduced maternal CENP-A (**Figure S7A**), showed similar results: reduced total centromeric CENP-A protein levels yet maintained a roughly equalized maternal-paternal ratio, along with increased CENP-C accumulation—particularly in the paternal pronucleus (**Figure S7B&C**). Although these effects are not as extreme as what was observed in our knockdown zygotes, this likely reflect the mixed genotype of oocytes generated from *Cenpa^+/-^* females. Interestingly, zygotes from *Cenpa* siRNA-treated oocytes or maternal *Cenpa^+/−^* females showed no significant differences in 2-cell progression or chromosome mis-segregation rates compared to their respective controls (**Figure 7F&G**), suggesting that reduced CENP-A levels can be tolerated if compensated by increased CENP-C recruitment.

**Figure 7:**
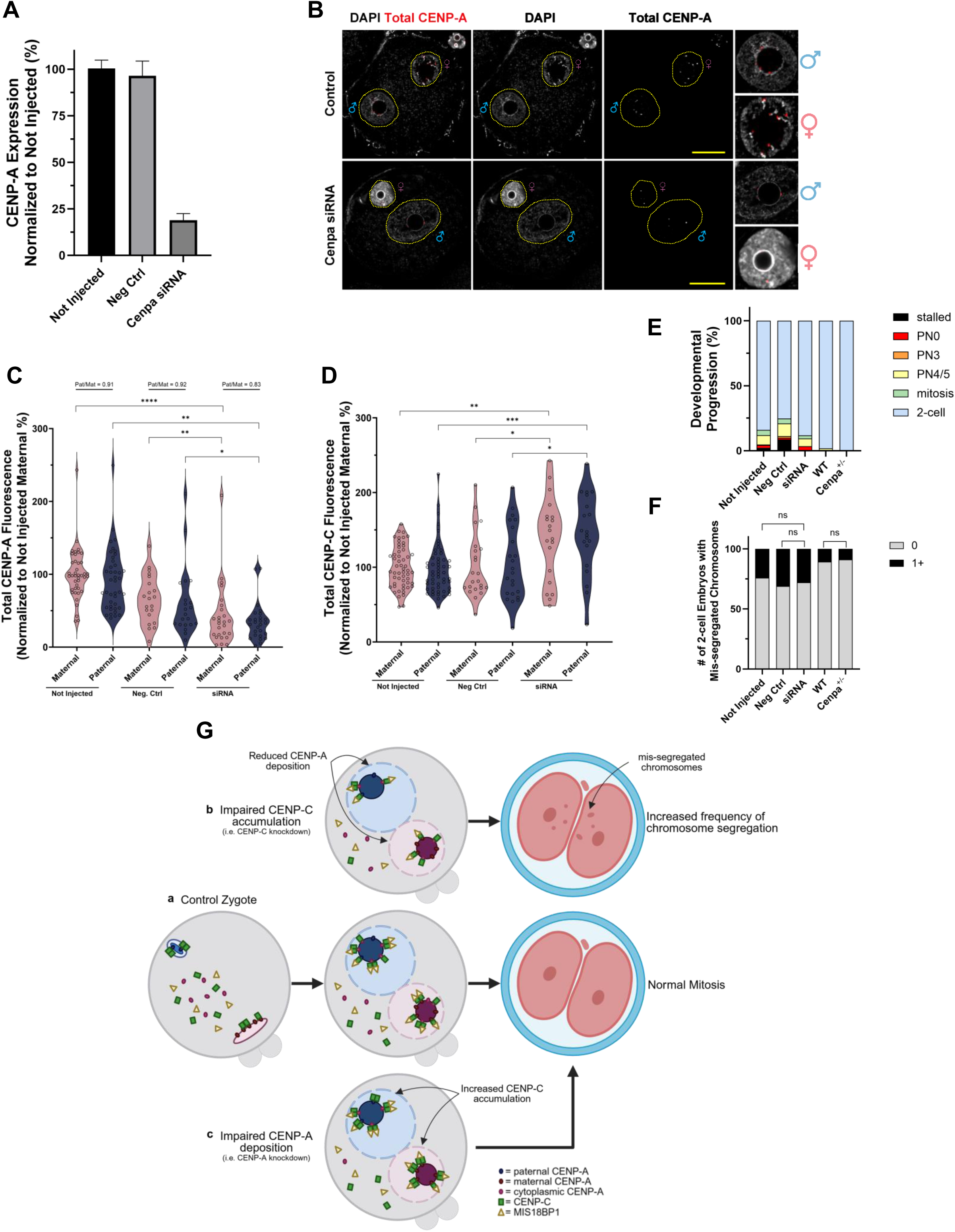
CENP-C compensates for acute CENP-A loss to preserve faithful chromosome segregation. A) RT-qPCR quantification of *Cenpa* transcripts in uninjected oocytes, negative control siRNA-injected (100– 150 nM), and *Cenpa* siRNA-injected oocytes (100–150 nM). Data represent n = 6 biological replicates, each with RNA from 10–20 oocytes, analyzed across three technical replicates. B) Representative images of CENP-A immunofluorescence in PN3–PN5 zygotes from control and *Cenpa* siRNA-injected oocytes. Scale bars: 20 µm. Images are from n = 5 IVF experiments using 4–5 CF-1 females and one (C57Bl/6J × DBA2)F1 male per experiment. C) Quantification of total CENP-A immunofluorescence in either maternal or paternal pronuclei from (B). Fluorescence measurements are normalized to the mean total maternal fluorescence in uninjected control oocytes. Shown are the average paternal-to-maternal ratios for each sample. Mean normalized fluorescence intensities are as follows: uninjected maternal = 100% for n = 41 zygotes, uninjected paternal = 91% for n = 41 zygotes, negative control maternal = 66% for n = 21 zygotes, negative control paternal = 60% for n = 21 zygotes, siRNA maternal = 42% for n = 26 zygotes, and siRNA paternal = 35% for n = 26 zygotes. D) Quantification of total CENP-C in PN2/3 zygotes from *Cenpa* siRNA-treated zygotes generated as described for (C) from n = 3 IVFs. Mean normalized fluorescence intensities are as follows: Not injected maternal = 100% for n = 54 zygotes, Not injected paternal = 98% for n = 54 zygotes, negative control maternal = 101% for n = 23 zygotes, negative control paternal = 104% for n = 23 zygotes, siRNA maternal = 138% for n = 20 zygotes, and siRNA paternal = 145% for n = 20 zygotes. E) The proportion of *Cenpa* siRNA-treated embryos or embryos derived from *Cenpa^+/-^* females (“*Cenpa^+/-^*”) collected at each stage shown in the legend. Embryos were allowed to develop for 24hrs. Not injected, neg ctrl, and siRNA groups were generated as described for (B-D). “Stalled” indicates zygotes that failed to form a proper paternal pronucleus. Percentages were calculated by combining all embryos generated across all replicates. Data is representative of n = 227 not injected embryos, n = 81 neg ctrl embryos, and n = 85 siRNA embryos, from at least n = 3 replicates using n = 5 CF-1 females and n = 1 (C57Bl/6J X DBA2)F1 male each. Data also includes n = 56 WT and n = 56 *Cenpa^+/-^* from n = 2 replicates cumulatively using n = 2-4 females of each genotype and n = 2 (C57Bl/6J X DBA2)F1 males. Not injected embryos are the same as in Fig. 6H. F) The percentage of 2-cell embryos with at least one chromosome segregation defects from (E). Percentages were calculated by combining all embryos generated across all replicates. The number of 2-cell embryos are as follows: Not Injected = 191, neg ctrl = 61, siRNA = 75, WT = 55, and *Cenpa^+/-^* = 56. G) The model for centromere remodeling and its impact on 2-cell embryo outcomes. Briefly, maternal and paternal centromeres in zygotes have very different CENP-A levels, but equalization is achieved prior to the first mitosis using maternal cytoplasmic pools of CENP-A, CENP-C, and MIS18BP1. In control zygotes (**a**), paternal genomes preferentially accumulate both CENP-C and MIS18BP1, which allows zygotes to resolve this inherited CENP-A asymmetry, allowing for a normal first mitosis. Zygotes with impaired CENP-A deposition but intact equalization (**b**) can compensate for weakened centromeres by recruiting additional CENP-C to stabilize them, allowing for normal mitosis. However, zygotes with reduced CENP-C levels have lower centromeric CENP-C and impaired CENP-A equalization, both of which contribute to increased chromosome mis-segregation during the first embryonic division. *: p < 0.05, **: p < 0.01, ***: p < 0.001, ****: p < 0.0001. ns indicates not significant.

Taken together, these findings highlight a critical role for CENP-C in mediating CENP-A equalization. While maternal CENP-A levels set the baseline for centromere identity, it is the availability and functional state of CENP-C – including its ability to dimerize – that governs the equalization process between parental genomes.

## Discussion

Centromeres are epigenetically defined chromosomal regions marked by the histone H3 variant CENP-A and are essential for kinetochore assembly, microtubule attachment, and faithful chromosome segregation. Despite all centromeres having this conserved function, the underlying DNA sequences are rapidly evolving, and the mechanisms regulating CENP-A are highly context-, cell type-, and –species-dependent. Here, we uncover how centromere identity is dynamically regulated during mammalian gametogenesis and early embryogenesis, revealing distinct sex-specific strategies and identifying CENP-C as a key mediator of CENP-A equalization in the zygote.

Firstly, we show that in the male germline, CENP-A levels progressively decline during spermatogenesis, with the most pronounced drop occurring at the onset of differentiation and continuing through meiosis (**Figure 1D&E and S1F-I**). This stepwise reduction precedes the global chromatin remodeling associated with spermiogenesis. Although this finding seems to contradict prior studies that have suggested that CENP-A is robustly retained throughout spermatogenesis (Brinkley et al., 1986; Das et al., 2022; Palmer et al., 1990; Palmer et al., 1991), their conclusions were based on late-stage spermatids and thus failed to capture early CENP-A dynamics in spermatogonia and differentiating cells. Our findings reveal a more nuanced view, uncovering dynamic regulation of CENP-A during early stages of male germ cell development—a pattern that is conserved in *Drosophila*, indicating evolutionary conservation of this regulatory strategy. However, unlike in mice and humans, CENP-A is redeposited post-meiosis in *Drosophila* and in plants (Dunleavy et al., 2012; Raychaudhuri et al., 2012; Schubert et al., 2014). Given these differences, it is not surprising that when we compare CENP-A levels between oocytes and sperm we find that mammalian gametes have more dramatic CENP-A asymmetry compared to *Drosophila* (**Figure 2**).We speculate that the recovery in flies occurring in the male germline may be needed due to rapid cell divisions in *Drosophila* embryos, which does not provide sufficient time to overcome large centromeric asymmetry.

Despite the remarkably low levels of CENP-A retained in sperm, centromere identity is preserved; suggesting that this residual amount of CENP-A maintained in sperm likely represents the minimal threshold required to epigenetically define a centromere. Supporting this, we observe limited variability in CENP-A intensity across individual sperm nuclei in humans and mice. In fact, the ∼10% of CENP-A retained in sperm – relative to levels in either spermatogonia or oocytes – is comparable to the lowest levels tolerated in somatic cells before chromosome segregation errors are observed (Fachinetti et al., 2013; Hoffmann et al., 2016). Such low abundance likely necessitates specialized stabilization mechanisms. Notably, among the examined CENP-A-associated factors, HJURP is the only centromeric protein retained in mature sperm (**Figure 1B and S1E**). Previous studies have shown that HJURP binding to CENP-A nucleosomes during passage of the replication fork maintains centromeric localization during mitotic divisions (Huang et al., 2015; Nechemia-Arbely et al., 2019; Zasadzińska et al., 2018). We propose that paternally inherited HJURP may mediate a similar stabilizing effect on CENP-A during extensive chromatin remodeling of the paternal genome – both during spermatogenesis as well as in the early zygote.

In contrast to males, CENP-A levels in the female germline increase markedly from primordial germ cells to prophase I arrested germinal vesicle (GV) oocytes (**Figure S3B**), followed by a modest decrease between GV to MII oocytes (**Figure S3E**). While centromeric CENP-A levels decrease slightly in MII oocytes, total CENP-A protein continues to increase – even after correcting for ploidy –(**Figure 2A&B**) suggesting the presence of a free cytoplasmic pool of CENP-A in oocyte. This interpretation is supported by our immunoblot analysis of multiple stage oocytes, photobleaching experiments, and reanalysis of previously published ribosome profiling data (Xiong et al., 2022) (**Figure S5**). This cytoplasmic pool of CENP-A is particularly intriguing because we show that CENP-A in mammalian oocytes is highly stable in GV and in vitro matured MII oocytes (**Figure S5B&C**). These findings align with prior genetic studies using conditional knockouts of *Cenpa* or *Mis18α*, a key licensing factor for CENP-A deposition, in growing oocytes which showed that CENP-A levels are largely preserved during oocyte maturation, even in the absence of new deposition (Das et al., 2023; Smoak et al., 2016). Together, these results suggest that the cytoplasmic pool of CENP-A may serve as a stable reservoir of protein needed to support centromeric remodeling in zygotic divisions.

Given the differences in centromere maintenance in both the male and female germlines, mammalian gametogenesis leads to the formation of two terminally differentiated gametes that contain drastically different levels of CENP-A (**Figure 2**). These differences in CENP-A maintenance may be due to differences in CENP-A post-translational modifications (PTMs) in male and female germline and/or, cell type-specific centromere-associated factors that can modify centromeric chromatin composition and structure. Indeed, CENP-A PTMs have been identified in mammalian cells, but their functions are either unexplored or currently under debate, and many of the PTMs identified in humans are not conserved even within the mammalian clade (Srivastava and Foltz, 2018). Importantly, this asymmetry in CENP-A is inherited by the mammalian zygote (**Figure 2D&E**). Although indirect methods have suggested that paternal nucleosomes are inherited and can affect early embryonic development (Hammoud et al., 2009; van der Heijden et al., 2008; Van De Werken et al., 2014), to our knowledge, our study is the first physical demonstration and direct visualization of intergenerational epigenetic inheritance of CENP-A in mice. Conceivably, our observations of intergenerational CENP-A inheritance may extend to other types of histones retained in sperm, suggesting that paternal nucleosomes can serve as epigenetic memory carriers from father to offspring.

To investigate how zygotes resolve parental asymmetry at centromeres, we tracked the temporal recruitment of key CENP-A-associated proteins – CENP-B, CENP-C, and MIS18BP1 – following fertilization. CENP-B and CENP-C were rapidly recruited to centromeres, while MIS18BP1 appeared later and was diffusely distributed throughout the nucleus, rather than centromere-restricted. Surprisingly, despite paternal centromeres having lower CENP-A levels, the paternal pronucleus disproportionately accumulated higher levels of CCAN and CENP-A deposition machinery, particularly CENP-C and MIS18BP1 (**Figure 5**). The pronounced enrichment of CENP-C at paternal centromeres is unexpected, given that its recruitment depends on CENP-A and CENP-B (Fachinetti et al., 2015; Falk et al., 2015, McKinnley and Cheeseman, 2016), which are present at equal or lower levels in the paternal genome compared to the maternal. This paradoxical asymmetry suggests that centromere assembly is shaped by intrinsic chromatin features specific to the parental genomes. Consistent with this, androgenetic embryos – comprising only paternal genomes – accumulate CENP-A to levels indistinguishable from those of normal zygotes, indicating that CENP-A equalization, including the establishment of its upper limit, occurs independently of maternal centromeric input or asymmetry sensing.

Importantly, maternal depletion of CENP-C impairs CENP-A equalization and leads to chromosome segregation errors at the first mitotic division (**Figure 6D-I**), underscoring the essential role of maternal CENP-C in zygotic centromere remodeling. Mechanistically, this process depends on CENP-C dimerization, which has been shown to recruit HJURP independently of the MIS18 complex (Flores Servin et al., 2023), raising the possibility that zygotic CENP-A loading proceeds via a non-canonical, MIS18-independent pathway. Unlike CENP-C knockdown, reducing maternally inherited CENP-A does not impair CENP-A equalization but does lower the overall CENP-A levels at both sets of parental centromeres. However, embryos with reduced CENP-A have a significantly higher amount of centromeric CENP-C levels (**Figure 7D and S7C**); suggesting a compensatory mechanism that may help stabilize centromeres. Others have previously proposed a similar mechanism in which they suggest that centromeric CENP-C can be stabilized in a CENP-B-dependent manner, allowing cells to compensate for diminished levels of CENP-A (Fachinetti et al., 2015). A similar inverse relationship between CENP-A and CENP-C was also observed in spermatocytes, where decreasing CENP-A levels coincided with increased CENP-C accumulation, reinforcing the idea that upregulation of CENP-C supports weaker centromere function and allows chromosome segregation in both spermatocytes and zygotes (**Figure 1B and S1E**) These observations are consistent with prior work showing that chromosome segregation defects in CENP-A–depleted cell lines emerge only after concomitant loss of CENP-C (Fachinetti et al., 2013; Hoffmann et al., 2016), and with reports that *Cenpc*-null embryos exhibit earlier developmental arrest than *Cenpa*-null embryos (Howman et al., 2000; Kalitsis et al., 1998).

While further work is needed to pinpoint inherited factors underlying sex-specific differences in centromeric chromatin states in gametes, our data suggest that asymmetric CENP-C recruitment in the zygote is key to the exponential increase in CENP-A observed in the paternal genome (**Figure 7H**). Proper regulation of centromeric chromatin during gametogenesis and early embryonic development is critical, as disruptions can lead to mitotically derived aneuploidies, which are common in mammalian embryos (Chuang et al., 2021; van Echten-Arends et al., 2011; Vázquez-Diez & FitzHarris, 2018). Notably, we also found that CENP-A and its associated proteins vary significantly, and these proteins are predominantly maternally inherited. Therefore, we propose that oocytes must maintain a sufficient pool of these proteins for successful embryogenesis, with variations in their levels potentially serving as biomarkers for oocyte quality and female fertility.

### Limitations of the study

All experiments using CENP-A-mScarlet were conducted with heterozygous *Cenpa^mScarlet/+^* mouse, as *Cenpa^mScarlet/mScarlet^* mice die between E8.5 and E15.5 (**see methods**). This lethality is likely due to weakened interactions between tagged CENP-A and CCAN components (Carroll et al., 2010). However, we do not anticipate that using this model has vastly affected our conclusions about centromere dynamics in gametogenesis and embryogenesis for multiple reasons: 1) Heterozygous animals are viable and fertile, and CENP-A-mScarlet faithfully localizes to chromocenters and colocalizes with CREST, with signal enrichment restricted only to chromosome ends. 2) Previous findings have elegantly shown that not all CENP-A nucleosomes need to interact with downstream kinetochore components to maintain a functional centromere (Bodor et al., 2014); indeed, only about 50% of CENP-A molecules are required to associate with kinetochores which should not be a problem in our model. 3) Any potential effects of the mScarlet tag on CENP-A’s biophysical behavior would apply equally to maternal and paternal genomes and therefore would not bias the comparative dynamics we report. However, to ensure validity of our conclusions, we tracked both tagged and endogenous CENP-A wherever feasible, consistently observing similar localization patterns and dynamic behavior during both gametogenesis and zygotic development. Finally, for robustness, all knockdown experiments were performed exclusively in wildtype oocytes, allowing us to assess the consequences of depleting fully functional, untagged CENP-A.

## Acknowledgements

We thank Drs. Richard Schultz, Paula Stein, and Carmen Williams, for their invaluable scientific discussions on *in vitro* maturation of oocytes and embryos, and the Hammoud Laboratory members for discussions. Additionally, we thank Mashiat Rabbani, Wenxin Xie, and Dominic Bazzano for their help editing the manuscript. We also thank Dr. Jon Oatley for sharing the Id4-eGFP mouse line, Dr. Iain Cheeseman for sharing the custom anti-mouse CENP-C antibody, and Dr. Alex Vargo for his help re-analyzing the Ribo-lite data. This work would not be possible without the assistance of Dr. Thomas Saunders in designing targeting vectors and members of the University of Michigan Transgenic Animal Model Core in the generation of the transgenic mouse model. Schematics were created with BioRender.com. This research was supported by National Institute of Health (NIH) grants 1DP2HD091949-01 (S.S.H.), R01HD104680 01 (S.S.H), R01HD058730 (B.E.B.), 1F31HD117648-01 (C.A.T.), F31HD100124 (G.M.), R35GM127075 (X.C), R35GM136340 (M. A., K.S.), training grants T32GM145470 (G.M.), T32GM007544 (C.A.T.), Rackham Predoctoral Fellowship (G.M.), and Open Philanthropy Grant 2019-199327 (5384) (S.S.H.). X.C. is a Howard Hughes Medical Institute Investigator.

## Author contributions

C.A.T., G.M., and S.S.H. conceived and designed the experiments. Experiments were carried out by C.A.T., G.M., E.L.F., D.A., K.J., B.M., M.A., R.R., S.C., and L.M. G.M. and S.S.H. received experimental and model system expertise from K.S. and X.C. respectively. Technical expertise as well as reagents were provided by A.D. and B.E.B., who have previously published a related study (Das et al., 2022). S.C., E.E.M., and A.S. also provided reagents. C.A.T., G.M., and S.S.H. wrote the manuscript and received input from all co-authors.

## Supplemental Figure Legends

**Supplemental Figure 1:**
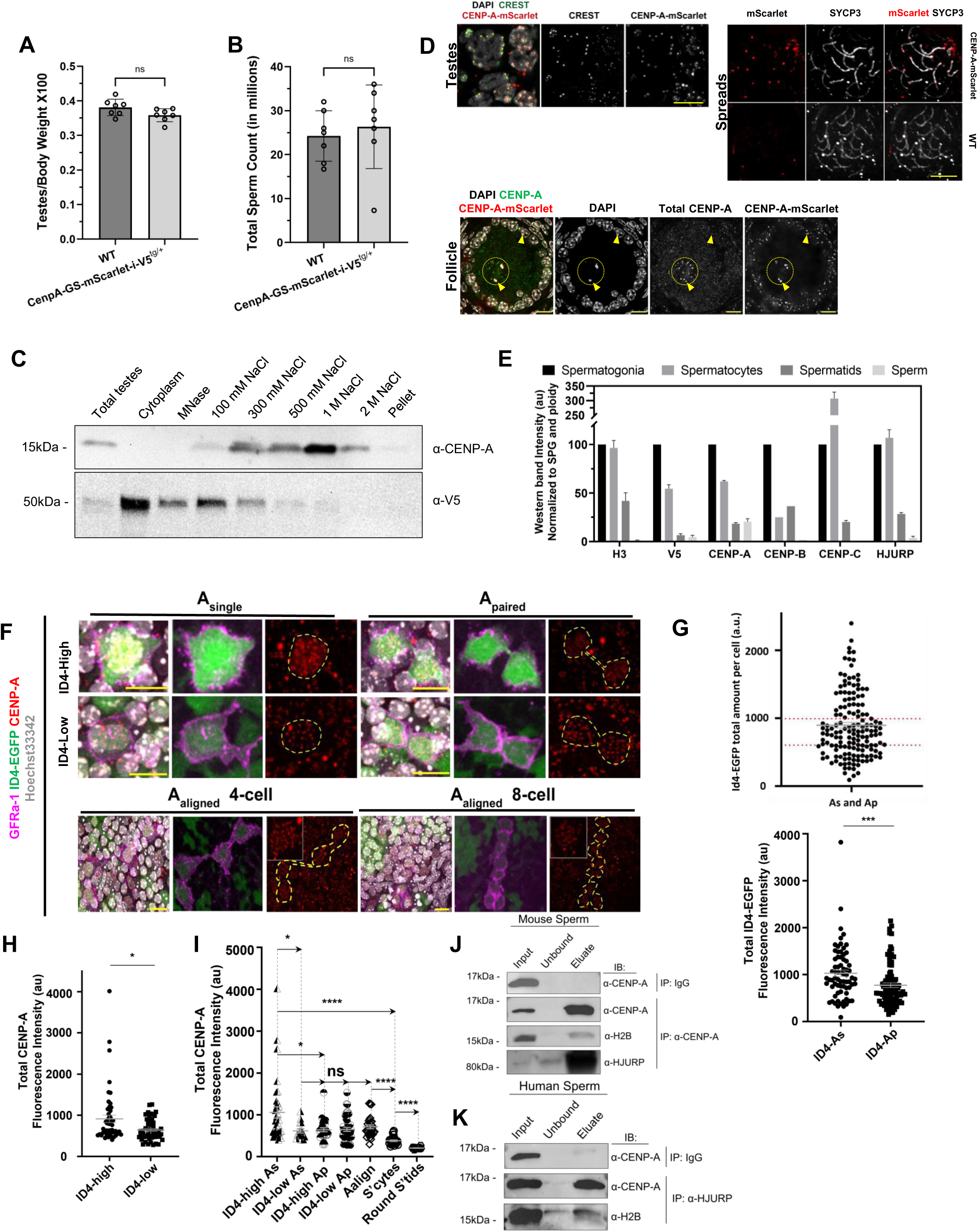
CENP-A loss coincides with germline stem cells differentiation. A&B) Testes-to-body weight ratio (A) and total sperm count (B) are maintained in *Cenpa^mScarlet/+^* mice. Each dot is the measurement from one mouse. ns indicates not significant. Mean values for testes/body weight ratio are as follows: WT = 0.3812 from n = 7 mice and *Cenpa^mScarlet/+^*= 0.3585 from n = 7 mice. Mean values for sperm count are as follows: WT = 24.24 million from n = 7 mice and *Cenpa^mScarlet/+^*= 26.32 million from n = 7 mice C) Representative immunoblots of high-salt fractionation of *Cenpa^mScarlet/+^* testes. Blots were stripped and re-probed for the indicated antibodies. D) Top left: CENP-A-mScarlet fluorescence colocalizes with CREST in *Cenpa^mScarlet/+^*tubule squashes. Representative images from n=3 staining experiments and n = 2 mice. Scale bar is 20um. Top right: *Cenpa^mScarlet/+^* enriches at chromosomal ends in meiotic spreads. Scale bar is 10um. Bottom: CENP-A-mScarlet colocalizes with total CENP-A in *Cenpa^mScarlet/+^*ovarian sections. Scale bar is 10um. E) Quantification of western band intensities from Fig 1B, normalized to ploidy and graphed as a percentage of spermatogonial protein levels. Mean values for H3 are as follows: spermatogonia = 100%, spermatocyte = 96.46%, spermatid = 41.77%, sperm = 1.26%. Mean values for V5 are as follows: spermatogonia = 100%, spermatocytes = 54.50%, spermatids = 6.50%, sperm = 4.59%. Mean values for total CENP-A are as follows: spermatogonia = 100%, spermatocytes = 61.77%, spermatids = 18.28%, sperm = 20.38%. Mean values for CENP-B are as follows: spermatogonia = 100%, spermatocytes = 24.94%, spermatids = 36.46%, sperm = 1.76%. Mean values for CENP-C are as follows: spermatogonia = 100%, spermatocytes = 307.02%, spermatids = 20.15%, sperm = 0%. Mean values for HJURP are as follows: spermatogonia = 100%, spermatocytes = 106.83%, spermatids = 28.35%, sperm = 3.88%. F) Quantification of CENPA levels in ID4-EGFP whole mount tubules stained with anti-GFRα1 antibody. Representative images from n = 3 staining experiments on n = 4 mice. Scale bar is 10 µm. G) Top: Quantification of endogenous ID4-EGFP fluorescence in all (n = 157) A_s_ or A_pr_ EGFP+ cells. ID4-EGFP high and low cutoffs are marked as horizontal dotted lines and were defined as the cells in the top or bottom 33 percentiles of fluorescence, the top cutoff is ≥992.4au and the bottom is ≤ 601.51au. Bottom: Quantification of endogenous ID4-EGFP fluorescence _single_ (As) and A_paired_ (Ap) cells. n_As_ = 71, n_Ap_ = 86. Mean fluorescence of ID4-EGFP in As and Ap are: As = 1027.11, Ap = 773.31. H) Quantification of total CENP-A immunofluorescence in ID4-EGFP high and ID4-EGFP low cells in As and Ap populations. n_Id4 high_ = 52, n_Id4 low_ = 52. Mean fluorescence of CENP-A in ID4-EGFP high and ID4-EGFP low is as follows: CENP-A_Id4 high_ = 912.94, CENP-A_Id4 low_ = 644.36. I) Quantification of total CENP-A immunofluorescence across all stages of ID4-High or ID4-Low undifferentiated spermatogonia, with spermatocytes (SPT) and round spermatids (RS) as controls. Mean fluorescence of CENP-A across the cell types is as follows: CENP-A_Id4 high As_ = 1059.53, CENP-A_Id4 low As_ = 617.45, CENP-A_Id4 high Ap_ = 645.07, CENP-A_Id4 high Ap_ = 653.15, CENP-A_Aalign_ = 645.42, CENP-A_SPT_ = 381.84, CENP-A_RS_ = 206.79. J) Immunoblot from CENP-A immunoprecipitation in mouse sperm, blotted for H2B and HJURP. Shown is a representative blot from n = 2 IPs on n = 2 mice. Membrane from the anti-CENP-A IP was stripped and re-blotted for the proteins indicated. K) Immunoblot from HJURP immunoprecipitation in human sperm, blotted for H2B and CENP-A. Representative blots from n = 3 HJURP immunoprecipitations on n = 3 deidentified human sperm samples. Membrane from anti-HJURP immunoprecipitation was stripped and re-blotted for H2B. IP = Immunoprecipitation, IB = Immunoblot. *: p < 0.05, **: p < 0.01, ***: p < 0.001, ****: p < 0.0001. ns indicates not significant.

**Supplemental Figure 2:**
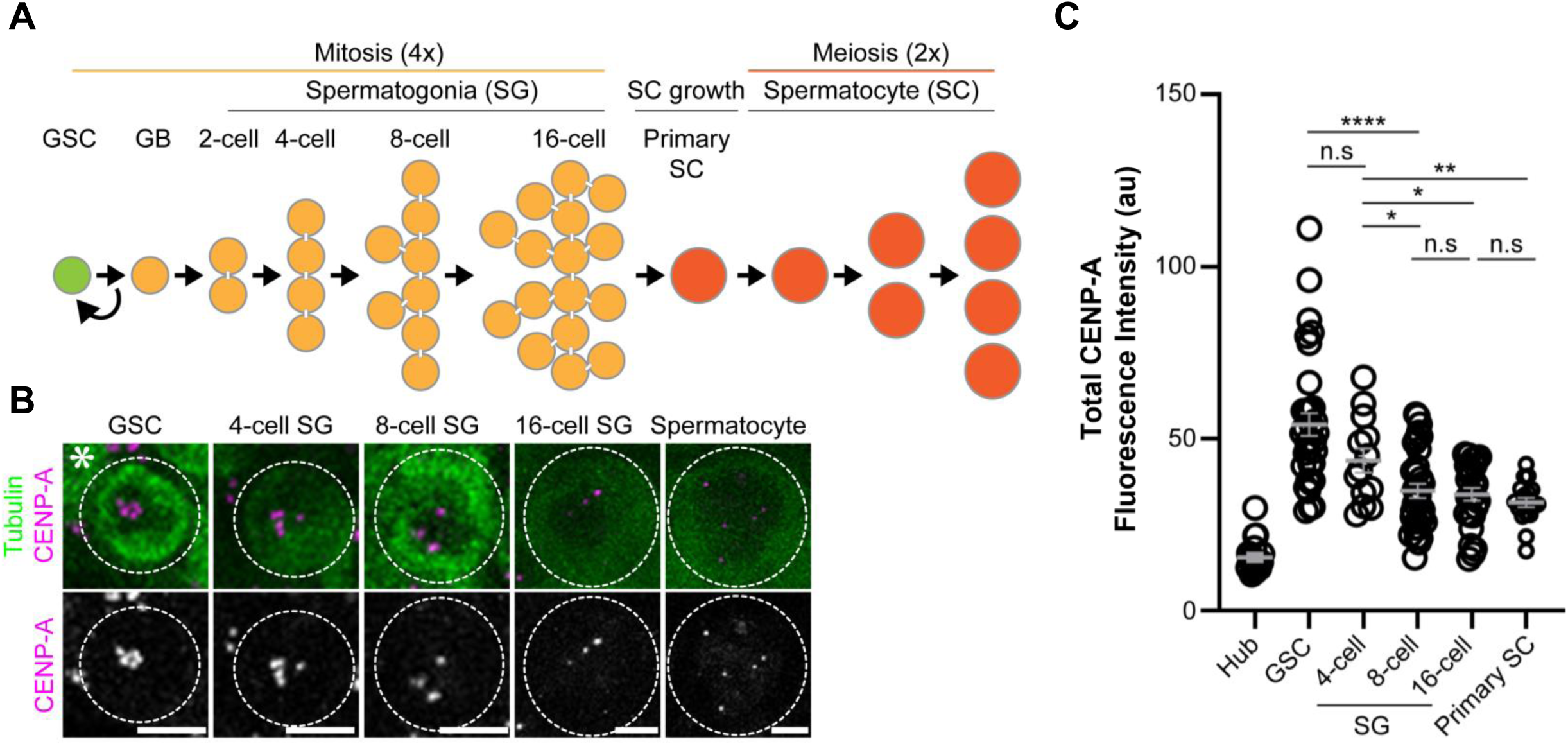
CENP-A loss during spermatogenesis is conserved in *Drosophila*. D) A schematic diagram of *Drosophila melanogaster* spermatogenesis. B) Representative images of each step of spermatogenesis. Germ cells expressing Tubulin-GFP and immunostained with anti-CENP-A^CID^ antibody. Asterisk: hub. Scale bars are 5um. C) Quantification of total fluorescence of CENP-A^CID^ across the stages of spermatogenesis. *: p < 0.05, **: p < 0.01, ****: p < 0.0001. ns indicates not significant.

**Supplemental Figure 3:**
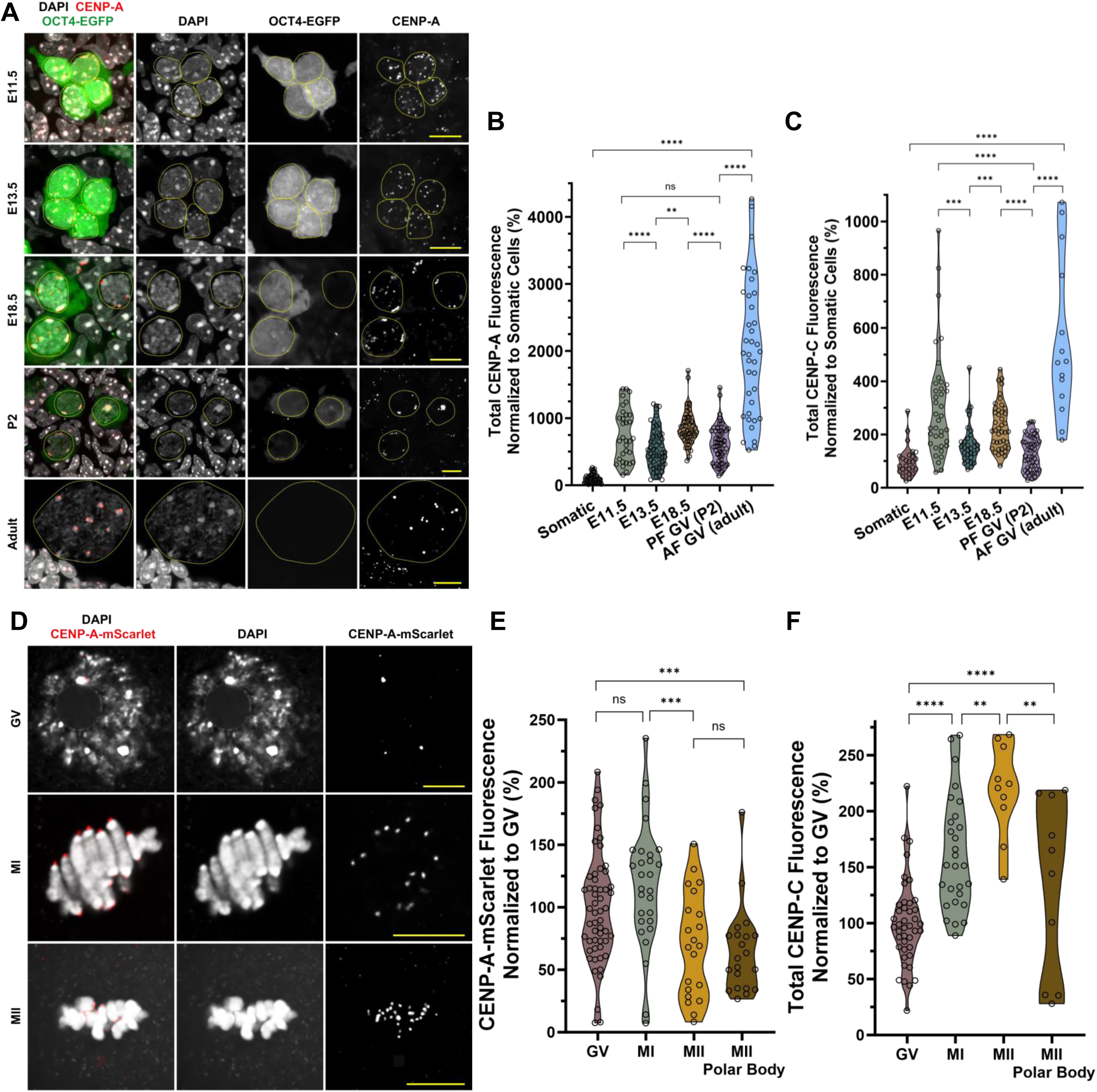
CENP-A levels increase during meiotic prophase I and are largely maintained during oogenesis. A) Representative images of total CENP-A immunofluorescence in OCT4-EGFP fetal gonads collected at the indicated embryonic and adult timepoints (eg. E11.5 is 11.5 days post conception). Scale bars are 20um, oogonia/oocytes are circled in yellow. Images are representative of n = 3 technical replicates from two gonads collected at each embryonic timepoint from one pregnant female mouse. B&C) Quantification of (B) total CENP-A and (C) CENP-C immunofluorescence at the indicated embryonic and adult timepoint. Each dot is the sum of CENP-A or CENP-C puncta in one cell. Mean intensity normalized to the mean somatic total CENP-A fluorescence is as follows: Somatic = 100%, n = 41 cells, E11.5 = 755% from n = 32 PGCs, E13.5 = 545% from n = 51 oogonia/oocytes, E18.5 = 871% from n = 50 oocytes, PF GV (P2) = 614% from n = 60 oocytes, AF GV (adult) = 2023% from n = 37 oocytes. Mean intensity of the total CENP-C fluorescence normalized to the mean somatic fluorescence is as follows: Somatic = 100% from n = 26 cells, E11.5 = 244% from n = 48 PGCs, E13.5 = 165% from n = 31 oogonia/oocytes, E18.5 = 232% from n = 42 oocytes, PF GV (P2) = 129% from n = 50 oocytes, AF GV (adult) = 553% from n = 14 oocytes. PF = primordial follicle and AF = activated follicle. D) Representative images of endogenous CENP-A-mScarlet fluorescence in geminal vesicle (GV), metaphase I (MI) and metaphase II (MII) stage eggs. Oocytes were collected from n = 6 *Cenpa^mScarlet/+^* females, and *in vitro* matured in n = 3 replicates. Scale bar is 10um. E&F) Quantification of endogenous (E) CENP-A-mScarlet fluorescence or (F) CENP-C immunofluorescence at each of the oocyte stages. Each dot is the sum of CENP-A-mScarlet or CENP-C puncta in one cell. Mean intensity of the CENP-A-Scarlet fluorescence is as follows: GV = 100% from n = 55 oocytes, MI = 117% from n = 28 oocytes, MII = 70% from n = 21 oocytes, MII polar body = 65% from n = 21 polar bodies. Mean intensity of the CENP-C immunofluorescence is as follows: GV = 100% from n = 42 oocytes, MI = 162% from n = 28 oocytes, MII = 219% from n = 10, MII polar body = 134% from n = 10 oocytes. *: p < 0.05, **: p < 0.01, ***: p < 0.001, ****: p < 0.0001. ns indicates not significant.

**Supplemental Figure 4:**
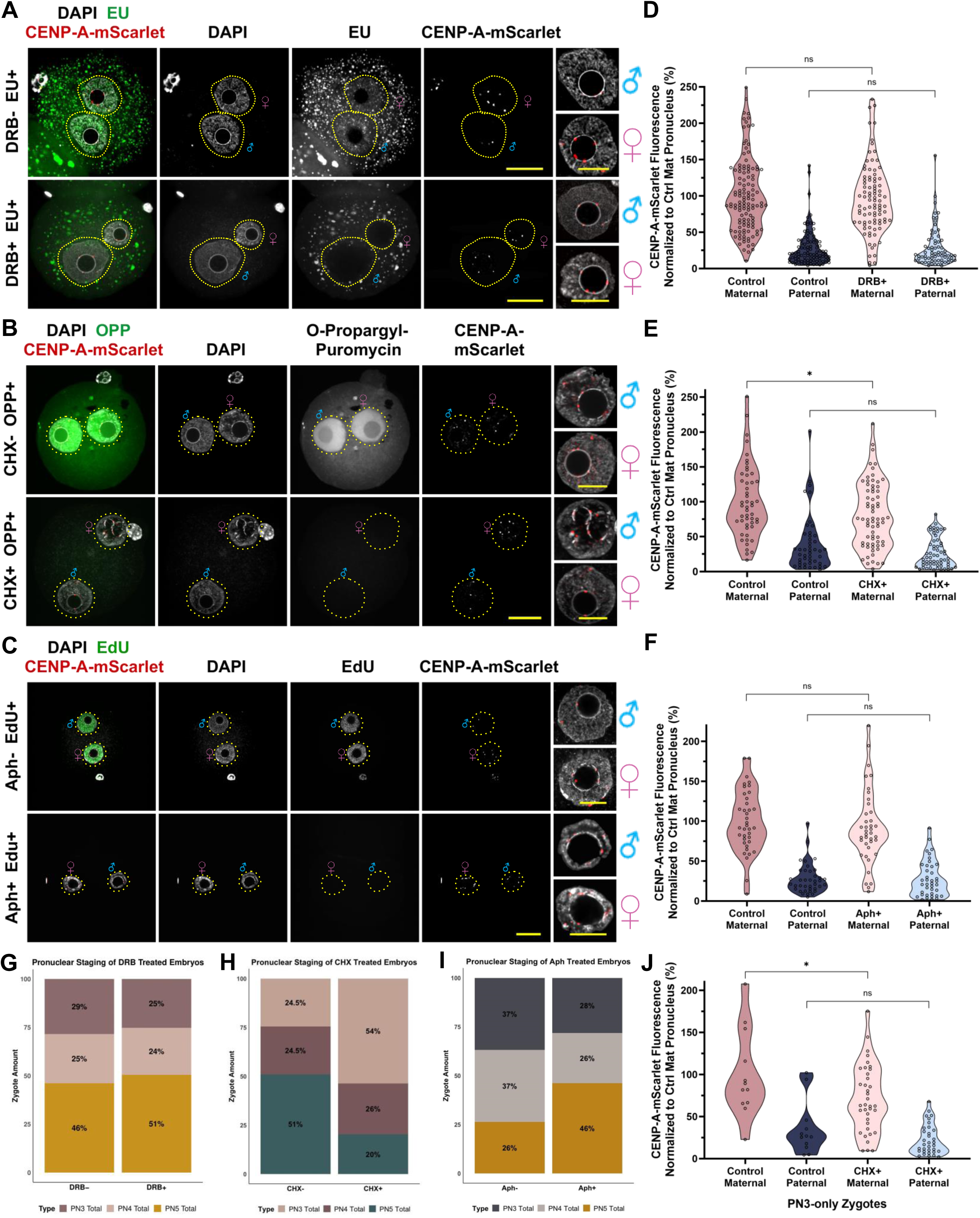
CENP-A deposition is independent of zygotic transcription, translation, and DNA replication. A-C) Representative images of zygotes collected from: A) Embryos treated with 200uM 5,6-dichloro-1-β-D-ribofuranosylbenzimidazole (DRB). 5-ethynyl uridine (EU) was used to monitor RNA synthesis and drug treatment efficiency. Embryos were collected at 10hpf from n = 5 IVFs with n = 13 *Cenpa^mScarlet/+^* females. B) Embryos treated with 100ug/mL cycloheximide (CHX). O-propargyl puromycin (OPP) was added to the culture media to monitor nascent peptide synthesis. Embryos were collected at 8.5hpf from n = 3 IVF experiments with n = 9 *Cenpa^mScarlet/+^* females. C) Embryos treated with 10ug/mL Aphidicoline (Aph). 5-ethynyl deoxyuridine (EdU) was used to monitor DNA synthesis. Embryos were collected at 8.5hpf from n = 3 IVFs with n = 9 *Cenpa^mScarlet/+^*females. Scale bars are 20um in main figure and 10um at the insets. D-F) Quantification of CENP-A-mScarlet direct fluorescence from embryos treated with (D) DRB, (E) CHX, (F) Aph. Each dot is the sum of CENP-A-mScarlet puncta in each pronucleus. Fluorescence measurements were normalized to mean fluorescence intensity in control maternal pronuclei. Mean fluorescence intensities shown in D) are as follows: Control maternal = 100% from n = 119 embryos, Control paternal = 28% from n = 98 embryos, DRB+ maternal = 95% from n = 91 embryos, DRB+ paternal = 29% from n = 71 embryos. Mean fluorescence intensities shown in E) are as follows: Control maternal = 100% from n = 51 embryos, Control paternal = 39% from n = 45 embryos, CHX+ maternal = 81% from n = 69 embryos, CHX+ paternal = 26% from n = 61 embryos. Mean fluorescence intensities shown in F) are as follows: Control maternal = 100% from n = 38 embryos, Control paternal = 28% from n = 38 embryos, Aph+ maternal = 92% from n = 39 embryos, Aph+ paternal = 28% from n = 39 embryos. ns indicates not significant. G-I) Quantification of pronuclear stages in zygotes treated with either (D) DRB, (E) CHX, or (F) Aph. J) Quantification of CENP-A-mScarlet direct fluorescence from only PN3 from stage embryos treated with cycloheximide (from panel E). Each dot is the sum of CENP-A-mScarlet puncta in one pronucleus. Fluorescence measurements were normalized to mean fluorescence intensity in control maternal pronuclei. Mean fluorescence intensities shown in J) are as follows: Control maternal = 100% from n = 12 embryos, Control paternal = 35% from n = 12 embryos, CHX+ maternal = 71% from n = 37 embryos, CHX+ paternal = 23% from n=33 embryos. *: p < 0.05, ns indicates not significant.

**Supplemental Figure 5:**
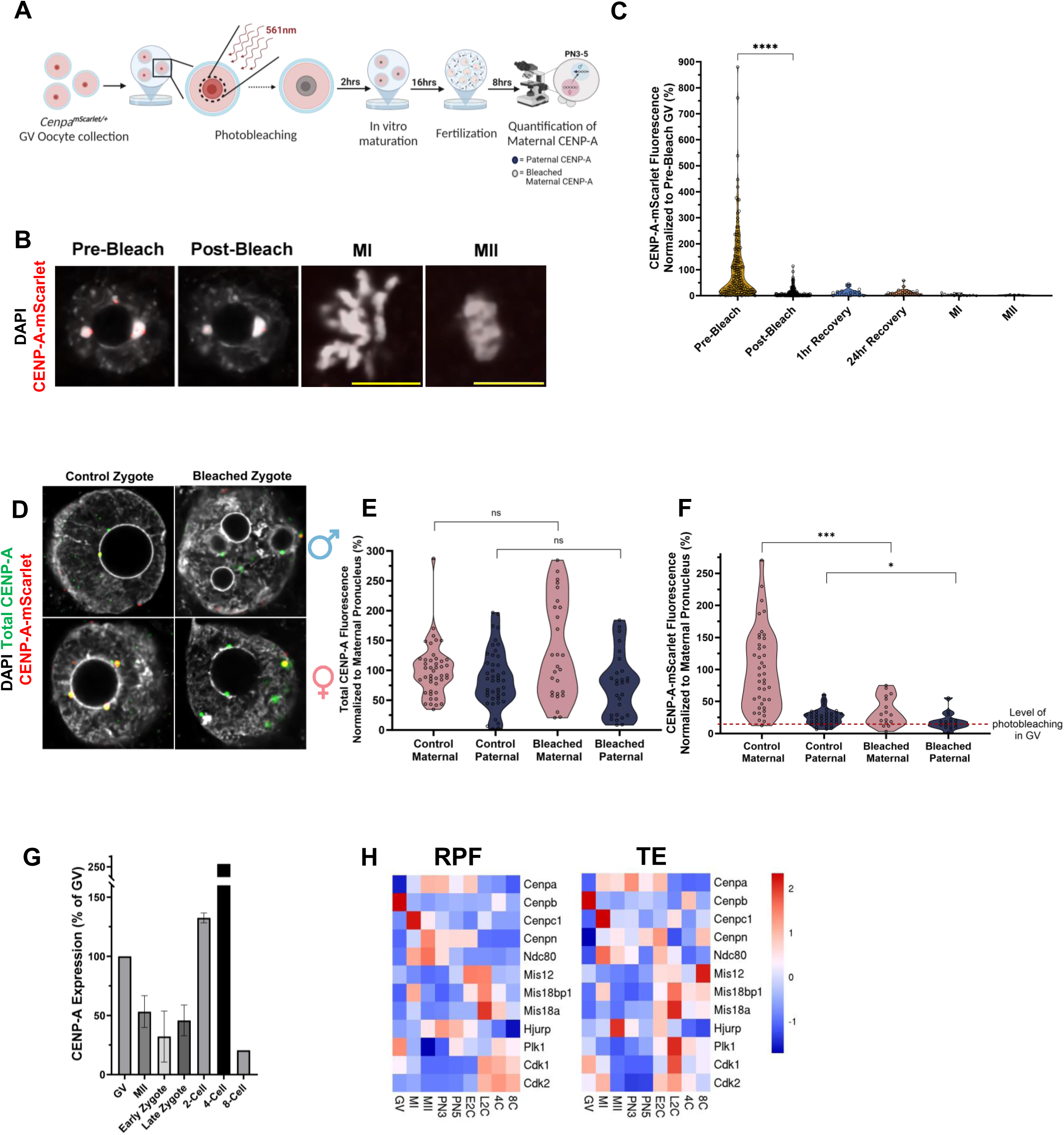
Maternally inherited cytoplasmic CENP-A is deposited at parental centromeres in the zygote. A) Overview of maternal CENP-A-mScarlet photobleaching experiments. B) Representative images of endogenous CENP-A-mScarlet fluorescence in oocytes before photobleaching, immediately after photobleaching, as well as after meiotic maturation to MI or MII stages. C) Quantification of CENP-A-mScarlet direct fluorescence in GV oocytes before bleaching, immediately after bleaching, 1– and 24-hours post-recovery, and in MI and MII oocytes. Each dot represents the total CENP-A-mScarlet puncta in one oocyte, normalized to the mean fluorescence intensity of pre-bleached GVs. Mean fluorescence intensities are as follows: Pre-bleach = 100% from n = 161 oocytes, post-bleach = 12% from n = 161 oocytes, 1hr recovery = 12% from n = 17 oocytes, 24hr recovery = 12% from n = 20 oocytes, MI = 3% from n = 14 oocytes, and MII = 3% from n = 5 oocytes. D) Representative images of endogenous and total CENP-A fluorescence in maternal and paternal pronuclei of PN3 zygotes from either control or photobleached experiments. All zygotes were treated with 100 µg/mL cycloheximide as in Fig. S4A. Images are representative of n = 5 independent photobleaching experiments using a total of 10 *Cenpa^mScarlet/+^* females. E&F) Quantification of either total CENP-A immunofluorescence (E) or direct CENP-A-mScarlet fluorescence (F) from the PN3-PN5 embryos represented in (D). Each dot is the sum of CENP-A-mScarlet or total CENP-A puncta in one pronucleus. Average fluorescence intensities, normalized to the mean control maternal total fluorescence shown in (E) are as follows: Control maternal = 100% from n = 46 embryos, Control paternal = 87% from n = 46 embryos, bleached maternal = 132% from n = 30 embryos, bleached paternal = 76% from n = 30 embryos. Average fluorescence intensities, normalized to the mean control maternal CENP-A-mScarlet fluorescence shown in (F) are as follows Control maternal = 100% from n = 41 embryos, Control paternal = 25% from n = 41 embryos, bleached maternal = 35% from n = 16 embryos, bleached paternal = 18% from n = 16 embryos. G) Quantification of *Cenpa* RNA transcripts from *in vitro* matured GV and MII eggs, early (6hpf) and late (12hpf) zygotes, and 2-, 4-, and 8-cell embryos. RNA levels are shown as a percentage of GV: GV = 100%, MII = 53.23%, Early Zygote = 32.13%, Late Zygote = 45.83%, 2-cell = 132.5%, 4-cell = 256.92%, 8-cell = 20.43%. n = 2 RT-qPCRs were run on oocytes and embryos from n = 6 mice. H) Heatmaps showing the relative expression of various centromere/kinetochore associated factors across various oocyte and embryo stages. E2C = early 2 cell L2C = late 2 cell 4C = 4 cell and 8C = 8 cell. RPF = ribosome protected fragment and TE = translation elongation. Heatmaps contain data reanalyzed from (Xiong et al., 2022). *: p < 0.05, ***: p < 0.001, ****: p < 0.0001. ns indicates not significant.

**Supplemental Figure 6:**
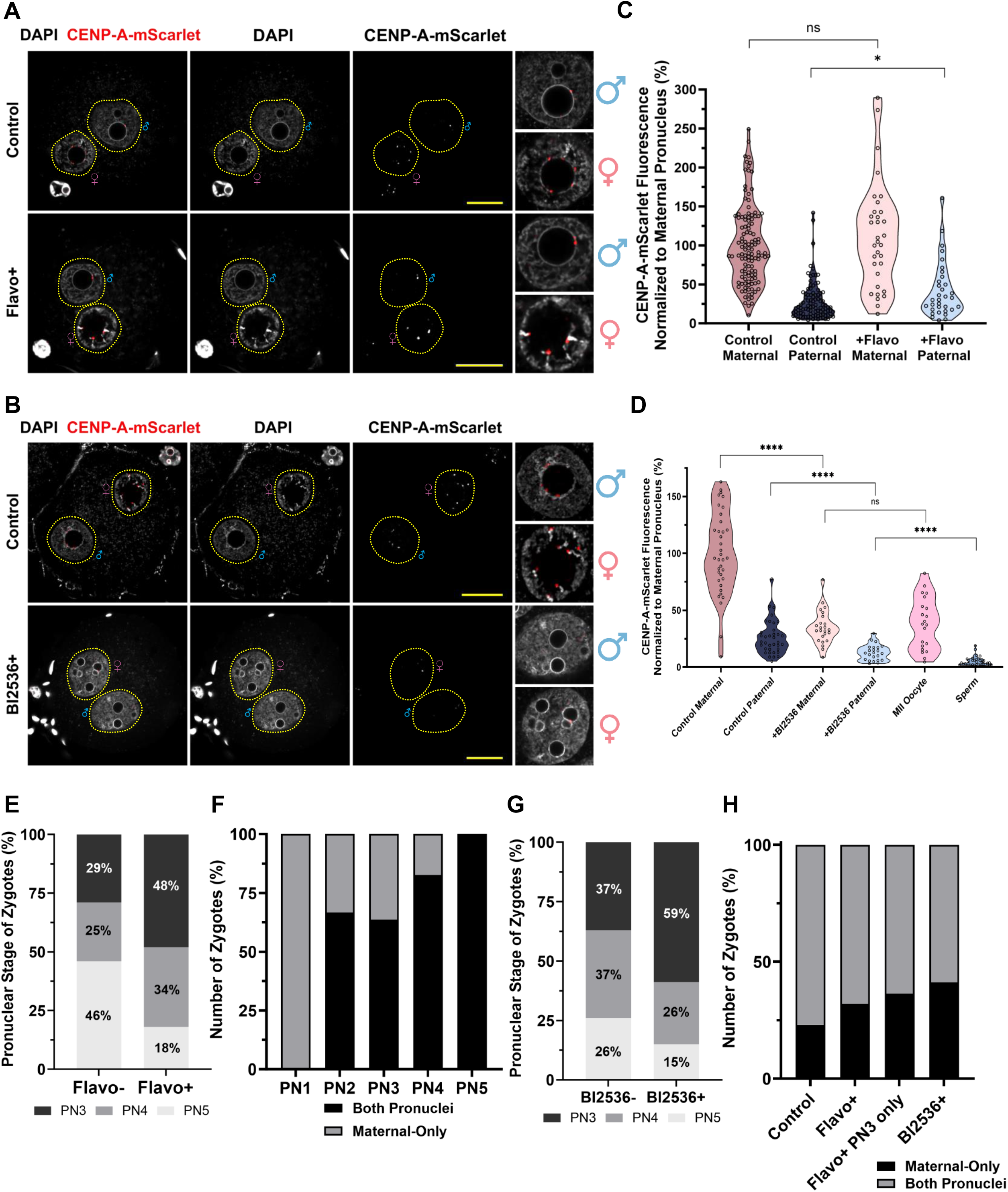
CDK1/2 and PLK1 kinase activity regulates CENP-A redistribution in the zygote. A&B) Representative images of zygotes collected from IVF experiments in which the embryos were treated with (A) 5uM Flavopiridol (Flavo) or (B) 10nM Bl2536. For each drug treatment, embryos were collected at 8.5hpf from n = 2 IVF experiments with n = 4 *Cenpa^mScarlet/+^* females. Scale bars are 20um or 10um in insets. C&D) Quantification of CENP-A-mScarlet direct fluorescence from embryos treated with Flavo (C) or Bl2536 (D). Each dot is the sum of CENP-A-mScarlet puncta in each pronucleus. Mean fluorescence intensities normalized to the average fluorescence quantified in the control maternal pronucleus shown in C) are as follows: Control maternal = 100% from n = 69 embryos, Control paternal = 28% from n = 98 embryos, Flavo+ maternal = 112% from n = 33 embryos, Flavo+ paternal = 42% from n = 33 embryos. Mean fluorescence intensities normalized to average control maternal fluorescence shown in D) are as follows: Control maternal = 100% from n = 119 embryos, Control paternal = 26% from n = 36 embryos, BI2536+ maternal = 35% from n = 24 embryos, BI2536+ paternal = 13% from n = 24 embryos, MII eggs = 38% from n = 21 oocytes (CENP-A-mScarlet in MII from **Figure S2E**), and sperm (paternally inherited CENP-A-mScarlet, see Fig. 3F) = 5% from n = 35 embryos. E) Quantification of pronuclear stages in zygotes treated with Flavopiridol. F) Quantification of the percentage of zygotes treated with Flavopiridol with maternal CENP-A-mScarlet deposited in both pronuclei or only in the maternal pronuclei at each pronuclear stage. G) Quantification of pronuclear stages in zygotes treated with BI2536. H) Quantification of the percentage of zygotes with maternal CENP-A-mScarlet in both pronuclei or only in the maternal pronuclei after 8hrs of fertilization in control, Flavo-treated, Flavo-treated PN3-only, and BI2536-treated embryos. *: p < 0.05, ****: p < 0.0001, ns indicates not significant

**Supplemental Figure 7:**
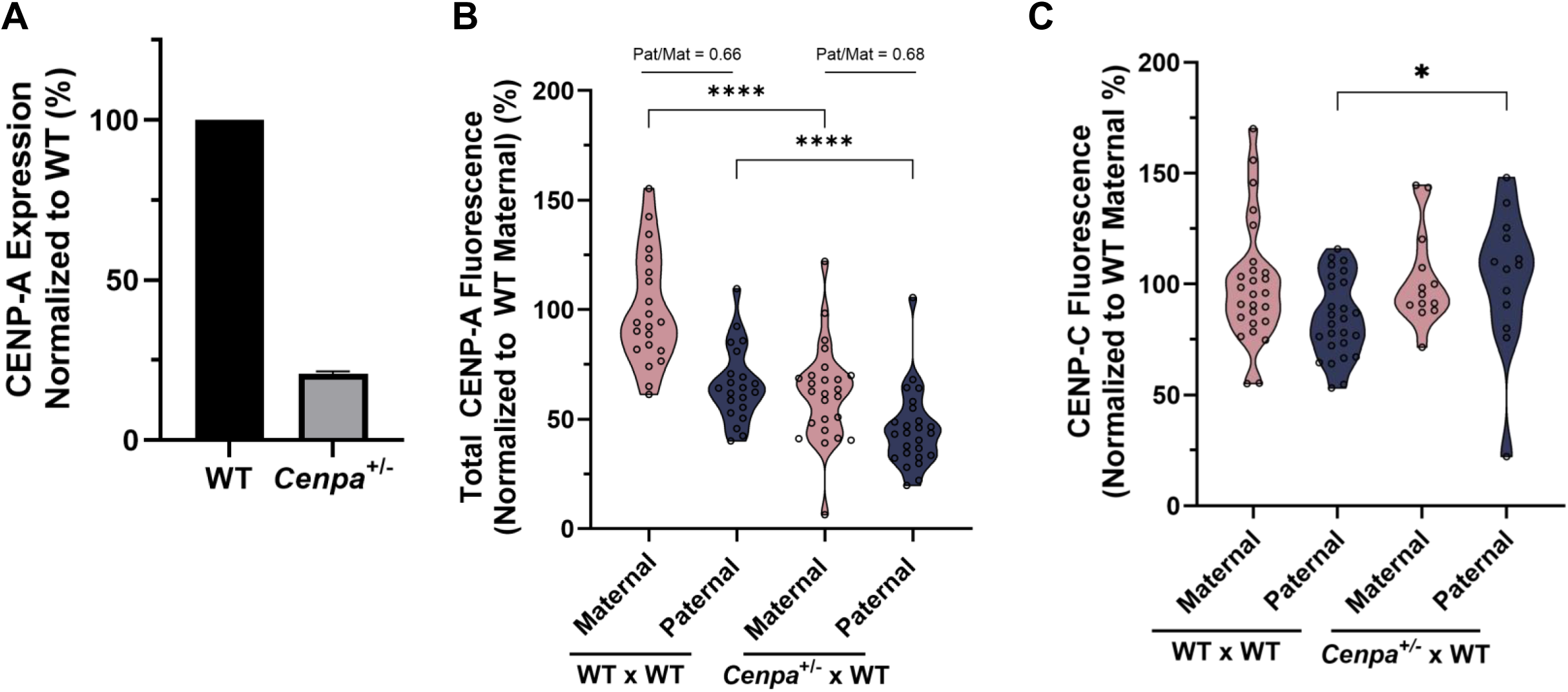
Zygotes derived from *Cenpa*^⁺/⁻^ females have reduced yet equalized CENP-A levels, along with increased CENP-C accumulation at paternal centromeres. A) Quantification of *Cenpa* RNA transcripts from WT or *Cenpa^+/-^* oocytes. Graph is representative of n = 2 replicates, with each replicate containing RNA from n = 10 meiotic oocytes for each sample, split across 3 technical replicates. B) Quantification of total CENP-A immunofluorescence in PN4/5 zygotes generated from either WT or *Cenpa^+/-^* females. Graph is representative of zygotes from n = 2 females per genotype fertilized with sperm from n = 1 WT (C57Bl/6J X DBA2)F1 male. Each dot is the sum of the total puncta in one pronucleus. Fluorescence intensities were normalized to the mean total fluorescence intensity of a WT x WT maternal pronucleus. Mean fluorescence intensities are as follows: WT x WT Maternal = 100% from n = 21 pronuclei, WT x WT Paternal = 66% from n = 21 pronuclei, *Cenpa^+/-^* x WT Maternal = 62% from n = 25 pronuclei, *Cenpa^+/-^* x WT Paternal = 45% from n = 20 pronuclei. C) Quantification of total CENP-C in PN2/3 stage zygotes that were generated as in (B). Data is representative of n = 2 females per genotype. Each dot is the sum of the total puncta in one pronucleus. Fluorescence intensities were normalized to the mean total fluorescence intensity of a WT x WT maternal pronucleus. Mean fluorescence intensities are as follows: WT Maternal = 100% from n = 26 pronuclei, WT Paternal = 85% from n = 26 pronuclei, *Cenpa^+/-^*Maternal = 102% from n = 13 pronuclei, *Cenpa^+/-^* Paternal = 103% from n = 13 pronuclei. *: p < 0.05, ****: p < 0.0001

## METHODS

### EXPERIMENTAL MODEL AND SUBJECT DETAILS

#### Mice

All experiments utilizing animals in this study were approved by the Institutional Animal Care and Use Committees of the University of Michigan (Protocols: PRO00006047, PRO00008135, PRO00010000, PRO00011691), John Hopkins University (Protocol: MO22A71), and Rutgers University (Protocol: 201702497) and was performed in accordance with the National Institutes of Health Guide for the care and use of laboratory animals. Briefly, mice were housed in an environment controlled for light (12 hours on/off) and temperature (21 to 23C) with *ad libitum* access to water and food (Lab Diet #5008 for breeding mice, #5LOD for non-breeding animals).

*Cenpa^mScarlet/+^* knock-in mice were generated on the C57BL/6N background using CRISPR/Cas9-mediated genome editing by the University of Michigan’s Transgenic Animal Model Core (Bindels et al., 2016). The sgRNA and donor oligo were designed as previously described (Haeussler et al., 2016). The guide RNA target sequence was selected according to the on– and off-target scores provided by the web tool CRISPOR (Haeussler et al., 2016) (http://crispor.tefor.net) and proximity to the target site. Ribonucleoprotein (RNP) complexes were formed by mixing the sgRNA (2.5 ng/uL) with Cas9 protein (IDT, 5 ng/uL) in Opti-MEM (ThermoFisher) and incubating at 37C for 10 minutes, at which time the donor oligo (IDT, 10 ng/uL) containing the intended transgene was added. The paternal pronucleus of zygotes generated from super-ovulated C57BL/6N females were microinjected with the RNP/donor oligo mix using a Femtojet 4i microinjector (Eppendorf). After injection, zygotes were moved to KSOM AA medium (Sigma), matured to the 2-cell stage, and transferred to oviductal ampullas of pseudopregnant CD-1 females. All animal procedures were carried out in accordance with the Institutional Animal Care and Use Committee and approved protocol of the University of Michigan. Offspring were genotyped for the transgene insertion by extraction of genomic DNA from a small ear biopsy. Mouse lines from two founding CRISPR/Cas9 transgene insertions were maintained separately for *Cenpa^mScarlet/+^* and utilized interchangeably in all experiments. Transgenic and control males were used for all experiments at least 8 weeks of age. Transgenic females were used 3-4 or 6-16 weeks.

The *Cenpa^+/-^* mouse line was generated for a previous publication (Smoak et al., 2016). Mice from this line were transferred to the University of Michigan animal facility where they are maintained on the C57Bl/6J genetic background for the experiments presented here.

#### Collection and preparation of human sperm and oocytes

For the human sperm collected for immunoprecipitation, the University of Michigan Institutional Review Board determined this study did not fit the definition of research involving human subjects (U.S. Department of Health & Human Services regulations at 45CFR46.102) because the research was intended to contribute to generalizable knowledge, the researchers did not interact with human subjects, nor obtained identifiable private information or identifiable biospecimens.

Discarded and de-identified semen samples were obtained from the University of Michigan Center for Reproductive Medicine. Briefly, semen samples were collected via masturbation from men presenting for routine semen analysis. Samples were allowed to liquefy at room temperature in a sterile container for 30 minutes before being processed for storage. The total semen samples were washed three times in PBS with centrifugation at 200g to remove seminal fluid and somatic cells in the resulting pellet were lysed in PBS (Thermo Fisher) + 0.1% SDS (Sigma) + 0.5% Triton-X-100 (Sigma) for 30 minutes on ice. The samples were centrifuged at 200g for 10 minutes and the final sperm pellet was snap frozen in liquid nitrogen and stored at –80C.

Human sperm and oocytes for immunofluorescence analysis were collected from the University of Michigan Reproductive Clinic. The University of Michigan Institutional Review Board approved the collection of these gametes and data (IRB00001996 approved 12/13/24, study ID HUM00264272). Oocytes were retrieved, fixed, and stored for staining on the same day according to the fixation protocol detailed below for mouse oocytes. Fresh ejaculates were collected from patients presenting in clinic for fertility treatment and were either processed fresh or washed, pelleted, and flash frozen in liquid nitrogen for later processing (see details below).

#### *Drosophila* gamete collection

Gametes were collected from knock-in *Drosophila melanogaster* (fly) strains. In collaboration with Fungene Inc. (Beijing, China), the following fly lines were generated using the CRISPR-Cas9 technology: CG613329 (*cid*) with Dendra2 tag at the internal site (between 118th – 119th codon), in order to generate the following fusion protein: CID N term-Dendra2-CID C term (Ranjan et al., 2019).

### METHOD DETAILS

#### Phenotypic assessment of *Cenpa^mScarlet/+^* mice

All phenotyping was carried out in males between 8 and 12 weeks of age. All weight measurements were recorded less than 5 minutes after euthanasia. Sperm were counted using a Makler chamber (Cooper Surgical SEF-MAKL) and performed as n=3 independent technical replicates per mouse (n=2-3 mice per founding line).

Timed matings were set up for *Cenpa^mScarlet/+^ x Cenpa^mScarlet/+^* mice to collect embryonic samples to assess homozygous lethality. Females (age at least 6 weeks) were plug checked every morning after pairing with a male (at least 6 weeks). After plugging females were separated into a different cage until the time of collection. Pregnant females were euthanized at E6.5, E7.5/8.5, or E15.5 to count the number of pups per litter as well as check for reabsorption sites. Time points were determined based on previously literature indicating *Cenpa^-/-^* embryos are viable until E8.5 (Howman et al., 2000). Litter sizes and pup genotypes at E15.5 are reported in the table below. Primer sequences for genotyping *Cenpa^mScarlet/+^* mice are as follows: wildtype allele forward 5’-GGCAGCTTAGGGGAGTCC-3’, mScarlet allele forward 5’-CTGAAAAGAACAGCCAATGTGTAAC-3’, and shared reverse primer 5’-TCCATATGCACTTTAAACCTCATAAATTC-3’.

Supplemental Methods Table 1.

| Embryonic Day | Average Litter Size (n litters) |
| --- | --- |
| E6.5 | 7(3) |
| E7.5 | 9(2) |
| E8.5 | 8(2) |
| E15.5 | 6(4) |

Supplemental Methods Table 2.

Supplemental Methods Table 2
| Embryonic Day | *Cenpa*<sup>+/+</sup>(%) | *Cenpa*<sup>+/mScarlet</sup>(%) | *Cenpa*<sup>mScarlet/mScarlet</sup>(%) |
| --- | --- | --- | --- |
| E15.5 | 4/18(22.2%) | 14/18(77.8) | 0/18(0) |

#### Microscopy

All fixed mouse samples were imaged on a Nikon A1R HD25 point scanning confocal microscope with a 60X 1.4NA oil objective (Nikon MRD71600) and motorized stage (Nikon TI2-S-SE-E) with 405nm, 488nm, 561nm, and 647nm solid state laser lines detected using a 1-channel GaAsP spectral detector (Nikon A1 DUV-B HD). The microscope platform was controlled in Nikon NIS Elements (Version 5.21.03). Images were acquired with 0.75um z-steps and ∼140nm xy pixel resolution.

Fixed human samples were imaged on a Leica Stellaris 5 point scanning confocal microscope using a 63x objective (HC PL APO 63×/1.40 oil C52).

#### Immunofluorescence on cryosectioned mouse testes

Testes were collected from WT C57BL/6J or *Cenpa^mScarlet/+^*males between 8-12 weeks of age. Mice were euthanized their testes were immediately removed and snap frozen in liquid nitrogen. The frozen testes were then embedded in OCT medium (Leica 39475237) on dry ice and stored at –80C in a sealed container. After embedding, the tissue was cryosectioned onto glass coverslips (Thermo Fisher) at a thickness of 10um and stored at –80C. To prepare for staining the coverslips were removed from storage at –80C and dried for 10-20 minutes at room temperature. The tissue sections were then fixed in 4% paraformaldehyde (Millipore Sigma) for 10 minutes, washed in PBS (Thermo Fisher) two times for 5 minutes each, and permeabilized in PBS + 0.5% TritonX-100 (Thermo Fisher) for 30 minutes at room temperature. Next, the tissue sections were washed twice in PBS, then blocked for 1 hour at room temperature in PBS + 3% BSA + 0.1% TritonX-100 filtered through a 0.2um filter (Millipore Sigma) before incubation with primary antibodies diluted in PBS + 3% BSA + 0.1% TritonX-100 filtered through a 0.2um filter at 4C overnight: rabbit α CENP-A at 1:250 dilution (Cell Signaling Technologies 2048), rabbit α CENP-C at 1:2000 dilution (gift from Ben E. Black). The next day, the tissue sections were washed four times for 15 minutes each at room temperature in PBS + 0.1% TritonX-100 then labelled with PNA Lectin-FITC (Vector Laboratories) diluted 1:1000, DAPI (Sigma) diluted 1:1000, and secondary antibodies diluted 1:1000 in PBS + 3% BSA + 0.1% TritonX-100 filtered through a 0.2um filter for either 1.5-2 hours at room temperature or overnight at 4C: Alexa Fluor 568 donkey α rabbit (Molecular Probes), Alexa Fluor 647 donkey α rabbit (Molecular Probes). Finally, the secondary antibodies were washed out with PBS three times for 10 minutes each, covered in a drop of VectaShield (Vector Laboratories) and a 25×50mm coverslip (Thermo Fisher), and sealed with nail polish.

#### Immunofluorescence on whole mount testes

Testicular tubules from adult *Cenpa^mScarlet/+^* male mice were prepared as previously published with some modifications (Smith and Braun, 2012). Briefly, the tunica albuginea was removed from the testes of euthanized males and the seminiferous tubules were teased apart in PBS. Interstitial cells were removed by incubating the tubules in 0.5mg/mL collagenase (Sigma) at room temperature (RT) for 5 minutes. The tubules were then rinsed three times with PBS, and fixed in 2% paraformaldehyde (Electron Microscopy Sciences) for 6 hours at 4°C. Paraformaldehyde-fixed tubules were then rinsed three times with PBS, and permeabilized with 0.25% NP-40 (US Biologicals) in PBST (PBS + 0.05% Tween (Thermo Fisher) for 25 minutes at RT. The tubules were rinsed three times for 5 minutes each with PBST and blocked using 5% normal donkey serum (Jackson Immuno Research) in PBST either 2 hours at RT or over-night at 4°C. The tubules were incubated with one of the following primary antibodies diluted in PBST + 5% normal donkey serum at 4C overnight: rabbit α CENP-A at 1:250 (Cell Signaling Technologies 2048), rabbit α CENP-A-Ser18ph at 1:100 (Active Motif 61483), Gt GFRa at 1:250 (R&D Systems AF429), Rb Sohlh1 at 1:200 (gift from Aleksandar Rajkovic), Rb Stra8 at 1:250 (Abcam ab49602), or mouse gH2A.X at 1:500 (EMD Millipore 05-636). The next day, the tubules were washed three times for 10 minutes each in PBST, then stained with DAPI at a 1:1000 dilution and an experiment-dependant combination of the following secondary antibodies at room temperature for 1 hour in PBST + 5% normal donkey serum: Alexa Fluor 488 donkey α goat at 1:500 (Molecular Probes), Alexa Fluor 488 donkey α mouse at 1:1000 (Molecular Probes), Alexa Fluor 568 donkey α rabbit at 1:1000 (Molecular Probes), Alexa Fluor 568 donkey α mouse at 1:1000 (Molecular Probes), Alexa Fluor 647 donkey α rabbit at 1:1000 (Molecular Probes), and/or Alexa Fluor 647 donkey α mouse at 1:1000 (Molecular Probes). After secondary antibody incubation, the tubules were rinsed three times for 10 minutes each with PBST. The tubules were then gently spread onto a coverslip with a paintbrush, dried briefly, covered in a drop of VectaShield (Vector Laboratories) and a 25×50mm coverslip (Thermo Fisher), and sealed with nail polish.

#### Preparation of spermatocyte spreads

Spermatocytes were mounted and spread onto glass slides as previously published with some modifications (Zelazowski et al., 2017). Briefly, a single testis from an adult C57Bl/6 or *Cenpa^mScarlet/+^* male was decapsulated by shaking at 150rpm for 15 minutes at 32C in 2mL testis isolation medium (TIM) comprised of 104 mM NaCl (Sigma) + 45 mM KCl (Sigma) + 1.2 mM MgSO4 (Sigma) + 0.6 mM KH2PO4 (Sigma) + 0.1% (w/v) glucose (Sigma) + 6 mM sodium lactate (Sigma) + 1 mM sodium pyruvate pH 7.3 (Sigma) with an additional 2mg/mL collagenase (Worthington) added for this step only. The tubules were washed three times in 15mL of TIM at room temperature before being resuspended in 2 mL of TIM + 0.7 mg/mL Trypsin (Sigma) + 4 ug/mL DNaseI (Roche 104159) and shaken at 150rpm for 15 minutes at 32C. The trypsin digest was stopped by diluting adding 1mL of fetal bovine serum (FBS) (Thermo Fisher) + DnaseI to 6.67ug/mL in the resulting single cell suspension and cell clumps were separated by gentle pipetting with a plastic transfer pipet and filtering through a 100um nylon mesh filter (Fisher Scientific). After filtering, add TIM to 15 mL and spin 200g for 5 min. The cells were then washed twice in TIM + 15uL of 400ug/mL DnaseI and once in 12mL PBS + 15uL of 400ug/mL DnaseI with centrifugation at 200g. The dissociated cells were then separated into twelve equal aliquots and two aliquots were spun at 200g for 5 min spread onto glass slides in turns. Each aliquot was resuspended in pre-warmed to 37C 80uL 0.1 M sucrose (Sigma) and incubated for 3-5 minutes at room temp., then 20uL of the aliquot was slowly spread along the length of a positively charged and precleaned glass slide. Each aliquot was spread onto four glass slides, which were each pre-covered with 65uL of prewarmed 1% paraformaldehyde pH 9.2 (EMD Millipore) + 0.1% TritonX-100 during the aliquot’s 3 to 5-minute incubation in 0.1 M sucrose. After spreading all of the aliquots of the testicular single cell suspension onto glass slides, the slides were allowed to dry slowly at room temperature for 3.5 hours. Finally, the slides were rinsed one in H2O and twice for five minutes each in PBS + 0.004% Photo-Flo 200 (Kodak 1464510), air dried, and stored at –80C.

#### Immunofluorescence staining of spermatocyte spreads

Spermatocytes spreads were immunostained following a previously published protocol with some modifications (Zelazowski et al., 2017). Briefly, the desired number of slides were removed from storage at –80C and dried for 10-20 minutes at room temperature, then immediately incubated with lambda phosphatase (New England Biolabs, P0753S) for 2 hours at 30C. Slides were then blocked for 30 minutes at room temperature in PBS (Thermo Fisher) + 1% fetal bovine serum (FBS) (Thermo Fisher) + 3% bovine serum albumin (BSA) (Sigma) + 0.05% Triton X-100 (Sigma) and incubated with primary antibodies diluted in PBS + 1% FBS + 3% BSA + 0.05% Triton X-100 overnight at room temperature: mouse α Sycp3 at 1:200 (Abcam ab97672) and rabbit α CENP-A at 1:250 (Cell Signaling Technologies 2048) or rabbit α CENP-C at 1:2000 (Das et al., 2022 and Iain Cheeseman). The next day, the spreads were washed with PBS + 1% FBS + 3% BSA + 0.05% Triton X-100 four times for 1 minute, 5 minutes, 10 minutes, and 15 minutes respectively, all at room temperature. Next, the spreads were stained with DAPI diluted 1:500 and secondary antibodies diluted 1:200 in PBS + 1% FBS + 3% BSA + 0.05% Triton X-100 for 45 minutes at room temp: Alexa Fluor 647 donkey α mouse (Molecular Probes), and Alexa Fluor 488 donkey α rabbit, respectively (Molecular Probes). The spreads were then washed again with PBS + 1% FBS + 3% BSA + 0.05% Triton X-100 four times for 1 minute, 5 minutes, 10 minutes, and 15 minutes respectively, all at room temperature. All incubations were done in a humified chamber to prevent evaporation. Finally, they were washed twice for five minutes each in PBS + 0.4% Photo-Flo 200 (Kodak 1464510), dried briefly, covered in a drop of VectaShield (Vector Laboratories) and a 25×50mm coverslip (Thermo Fisher), and sealed with nail polish.

#### Flow cytometry

Testes from adult *Cenpa^mScarlet/+^* male mice were dissociated, stained, and flow sorted as previously published with some modifications (Green et al., 2018). Briefly, the tunica albuginea was removed from the testes of euthanized males and the seminiferous tubules were transferred to 10ml of Advanced DMEM:F12 media (Thermo Fisher) + 200ul of 10 mg/ml Collagenase IA (Sigma) + 100ul of 20mg/mL Dnase I (Sigma). Tubules were dispersed by shaking at 35C for 5min at 215rpm and allowed to settle for 1 minute at room temperature. Excess media was removed with a sterile pipette, leaving ∼2mL with the settled tubules. The tubules were then dissociated at 35C with horizontal shaking at 215 rpm for 5 minutes in an additional 10mL of Advanced DMEM:F12 media (Thermo Fisher) + 200ul of 20mg/ml DnaseI (Sigma) + 5mg of Trypsin (Thermo Fisher). The trypsin was quenched with the addition of 3mL of fetal bovine serum (FBS) (Sigma). The resulting single cell suspension was filtered through a 100um strainer (Thermo Fisher), washed in PBS (Thermo Fisher), pelleted at 300g for 5 minutes, and re-suspended in 6mL MACS buffer containing 0.5% BSA (Miltenyi Biotec). The suspension was stained with Hoechst 3342 (Thermo Fisher) and propidium iodide (PI) (Thermo Fisher) as previously published with no modifications (Gaysinskaya et al., 2014). Spermatogonia (2n), spermatocytes (4n, mostly pachytene and diplotene), and round spermatids (1n) were isolated from the stained live single cell suspension based on DNA content and DNA compaction using a FACS ARIA II/III flow cytometer (BD Biosciences) and Bigfoot Spectral Cell Sorter (Thermo Fisher) available at the University of Michigan’s Flow Cytometry Core. Gates and sorting conditions were adapted from previously published methods and optimized for each biological replicate (Gaysinskaya et al., 2014; Green et al., 2018). Sorted cell populations were immediately pelleted at 600g and snap frozen in liquid nitrogen before storage at –80C.

#### Subcellular and high-salt fractionation of tagged CENP-A testes

Cells were fractionated using a high-salt gradient, as we have published previously (Moritz et al., 2023). Briefly, flash frozen testes were homogenized (Dounce homogenizer) and then washed with PBS. Cells was incubated with 0.1% CTAB for 5 min on ice to remove tails from the spermatids and then washed with 50mM Tris-HCl pH 8.0 several times. Cells were lysed for 15 min on ice and then spun down (800 g for 15 min). The supernatant (cytoplasmic fraction) was collected, and the nuclei were digested with 1U of MNase at 37C for 30min. The digestion was quenched with EGTA, then spun down (400g for 10min). The supernatant was collected (MNase fraction) and the remaining sample was fractionated via sequential incubations (30 min each) with varying NaCl concentrations (100 mM, 300 mM, 500 mM, 1 M, and 2 M). Salt fractions were incubated with 20% TCA overnight at –20C to precipitate the proteins and then resuspended in water before immunoblotting. Immunoblotted membranes were incubated with antibodies against CENP-A at 1:250 (Cell Signaling Technologies 2048), V5 at 1:1000 (BioRad MCA1360). Primary antibodies hosted in mouse were detected with HRP-conjugated goat α mouse IgG at 1:10,000 (Abcam ab6721) and primary antibodies hosted in rabbit were detected using HRP conjugated mouse α rabbit IgG light chain at 1:5,000 (Abcam ab99697).

#### Sperm chromatin immunoprecipitations

Mouse sperm was collected in a swim up from the caudal epididymis and vas deferens as previously described (Brykczynska et al., 2010), and mature human sperm was collected and snap frozen as detailed above. Chromatin from both human and mouse sperm was immunoprecipitated using a previously published protocol with some modifications (Yoshida et al., 2018). Briefly, 30 million mature sperm were resuspended in 1mL of PBS and cross-linked by 1% formaldehyde (Sigma) for 10 minutes at room temperature before quenching the fixation by adding Tris pH 7.5 to a final concentration of 0.2 M and incubating for another 10 minutes at room temperature. The fixed sperm was then washed twice in sperm decondensation buffer: 5 mM HEPES pH 8.0 (Sigma) + 1 mM PMSF (Sigma) + 0.2% NP-40 (Sigma) + 10 mM EDTA (Sigma) + 5 mM NaCl (Sigma) + 1.2 M urea (Sigma) + 10 mM DTT (Sigma) + 2X complete protease inhibitor cocktail (Sigma). After washing, the sperm was resuspended in sperm decondensation buffer with 1 mg/mL heparin sodium salt (Sigma) at a concentration of 15 million sperm per 3 mL decondensation buffer. The mouse sperm was then decondensed by incubating for 5 hours at 42C, while human sperm was decondensed by incubating for 2 hours at 42C. The decondensed sperm was then washed twice in lysis buffer: 50 mM HEPES pH 7.5 (Sigma) + 140 mM NaCl (Sigma) + 1 mM EDTA (Sigma) + 10% glycerol (Sigma) + 0.5% NP-40 (Sigma) + 0.25% Triton X-100 (Sigma) + and 1X complete protease inhibitor cocktail (Sigma). After washing, decondensed mouse sperm was treated with lambda phosphatase (NEB) by resuspending the sperm in 1X NEBuffer for Protein MetalloPhosphatases supplemented with 1 mM MnCl2 (NEB) and incubating for 30 minutes at 37C. The decondensed human sperm was not treated with lambda phosphatase. Both mouse and human sperm were then resuspended in 50 mM Tris-HCl pH 8.0 (Sigma) + 10 mM EDTA (Sigma) + 1% SDS (Sigma) + 1X complete protease inhibitors (Sigma) + 1 mM PMSF and sonicated using four cycles of 30 seconds on/off. The sonicated sperm was diluted 1:10 in 10 mM Tris-HCl pH 8.0 (Sigma) + 100 mM NaCl (Sigma) + 1 mM EDTA (Sigma) + 0.5 mM EGTA (Sigma) + 1% Triton X-100 (Sigma) + 0.1% sodium deoxycholate (Sigma) + and 1X complete protease inhibitor cocktail (Sigma) to dilute SDS to a final concentration 0.1%. Next, the sperm chromatin was spun down for 10 minutes at 20,000g and the insoluble fraction was saved for immunoblot analysis. The soluble chromatin was precleared by gently rotating with magnetic Protein A beads (Invitrogen 10002D) for 1 hour at 4C, pre-clearing beads were removed and 10% of the soluble chromatin was saved as input for immunoblot analysis. Chromatin was immunoprecipitated by incubating the soluble chromatin with the desired antibody pre-bound to magnetic Protein A beads for at least 14 hours at 4C with gentle rotation. Antibodies used for immunoprecipitation include rabbit α CENP-A at 1:200 (Cell Signaling Technologies 2048) for mouse sperm, rabbit α HJURP at 1 ug per 10 million sperm (Abcam ab100800) for human sperm, and rabbit IgG isotype control at either 1:200 or 1 ug per 10million sperm (Invitrogen 10500C) for mouse or human sperm respectively. The unbound fraction was saved for immunoblot analysis, and the immunoprecipitation was washed four times in 50 mM HEPES pH 7.0 (Sigma) + 0.5 M LiCl (Sigma) + 1 mM EDTA (Sigma) + 0.7% sodium deoxycholate (Sigma) + 1% NP-40 (Sigma), then twice in 10 mM Tris-HCl pH 8.0 (Sigma) + 1 mM EDTA (Sigma), and the immunoprecipitation was finally eluted in 10 mM Tris-HCl pH 8.0 (Sigma) + 1 mM EDTA (Sigma) + 1% SDS (Sigma) + 1X complete protease inhibitor cocktail (Sigma). The input (10% of IP), 10% of the unbound, and 100% of the elution were immunoblotted from each experiment and antibody. Immunoblotted membranes were incubated with either rabbit α HJURP at 1:500 (Abcam ab100800), rabbit α CENP-A at 1:250 (Cell Signaling Technologies 2048), mouse α CENP-A at 1:1000 (GeneTex GTX13939), mouse α H2B at 1:500 (Abcam ab52484), or rabbit α H2B at 1:1000 (Cell Signaling Technologies 12364). Primary antibodies hosted in mouse were detected with HRP-conjugated goat α mouse IgG at 1:10,000 (Abcam ab6721) and primary antibodies hosted in rabbit were detected using HRP conjugated mouse α rabbit IgG light chain at 1:5,000 (Abcam ab99697).

#### Human sperm immunofluorescence

Fresh ejaculate samples were centrifuged at 2,000 rpm for 6 minutes. The supernatant was discarded, and the resulting pellet was resuspended in 500 µL of PBS. Samples were sonicated at 40% power for 20 seconds, rested for 15 seconds, and then sonicated for an additional 5 seconds to remove sperm tails. Following sonication, samples were centrifuged again at 2,000 rpm for 6 minutes, and the supernatant was discarded. The pellet was resuspended in a Percoll solution composed of 350 µL Percoll (Sigma P1644) and 150 µL DMEM (Gibco) and centrifuged at 2,000 rpm for 5 minutes. The pellet was then resuspended in DMEM, and sperm count was assessed. Samples were centrifuged once more at 2,000 rpm for 5 minutes and resuspended in PBS at a concentration of 25,000 sperm/mL. Cells were cytocentrifuged onto microscope slides using a Cytospin device (Hettich Rotofix 32 A). Sperm were permeabilized in 0.5% CTAB (EMD Millipore; 219374) in PBS for 30 minutes at room temperature. Slides were washed in 50 mM Tris buffer (pH 8.0) for 2 minutes. Sperm heads were then decondensed using 25 mM DTT in 500 mM NaCl for 45 minutes. Samples were fixed in 4% paraformaldehyde for 15 minutes, then treated with lambda phosphatase (New England Biolabs, P0753S) for 2 hours at 30°C. Following treatment, samples were blocked in 3% BSA (Sigma) 0.1% Triton X-100 in PBS for 1 hour at room temperature. For immunostaining, samples were incubated overnight at 4°C with human CENP-A antibody (1:250; Cell Signaling, 2186) diluted in blocking solution. Slides were then washed three times in PBS 10 minutes each wash. Secondary staining was performed using Alexa Fluor 594-conjugated secondary antibody (1:500; Invitrogen A-11012) and DAPI (concentration/source needed) for 2 hours at room temperature. Slides were washed again three times in PBS, 10 minutes each, and mounted with Vectashield.

#### CID quantification in Drosophila gametes

Mature oocytes and mature sperm were isolated from dissected adult *Drosophila melanogaster* females and imaged using a Leica SPE confocal microscope. Quantification of the total CID signal per gamete was performed following a previously established method (Ranjan et al., 2019; Ranjan et al., 2022). Fixed samples were immunostained and imaged for endogenous CID-Dendra2 fluorescence along with Hoechst staining. For quantification, un-deconvolved raw images were collected as 2D Z-stacks and saved as un-scaled 16-bit TIFF files. Fluorescence signals were manually quantified using Fiji (ImageJ). A circular region of interest (ROI) was drawn around the entire fluorescence signal for each gamete (either CID-Dendra2 or Hoechst signal). An identical-sized circle was drawn in an adjacent background area. The raw integrated density (RawIntDen), representing the sum of pixel gray values within each ROI, was determined for both signal and background regions. The net fluorescence signal (Foreground signal) was calculated by subtracting the background signal from the raw signal. The total fluorescence per gamete was then determined by summing the background-corrected fluorescence from all Z-stacks. All CID quantifications were conducted using this approach.

#### In vitro fertilization

For mouse zygotes and early embryos collected by standard in vitro fertilization (IVF), oocytes collected from either C57Bl/6J females (Jax) or *Cenpa^mScarlet/+^* females. Females were superovulated with an intraperitoneal injection of 100 uL HyperOva (Card KYD-010-EX-X5) and 5-7.5 IU of human chorionic gonadotropin (hCG) (Sigma CG5) at 60 and 14-16 hours respectively before oocyte collection. See “Oocyte collection and maturation” for the isolation of oocytes from *in vitro* maturation (IVM). Sperm was collected from either (C57Bl/6J X DBA2)F1 males (Jax), or *Cenpa^mScarlet/+^*males. (C57Bl/6J X DBA2)F1 and *Cenpa^mScarlet/+^* males were housed individually for three to seven days prior to IVF. Males were 8-12 weeks of age and females 3-4 or 6-10 weeks of age at the time of gamete collection.

Media for sperm capacitation and IVF incubation, and if applicable later stage embryo maturation, was allowed to equilibrate in an incubator at 37C with 5% CO2 the night before IVF. The day of IVF, males were euthanized 30 minutes to 1.25 hours before oocyte collection. Mature sperm was collected by mincing the epididymis and vas deferens and capacitating in 500 uL Research Vitro Fert Media (Cook Medical K-RVF-50) or Human Tubule Fluid (HTF, Sigma MR-070-D) for 10 to 15 minutes at 37C and 5% CO2. The epididymis and vas deferens tissue were then removed, sperm concentration and percent motility were counted using a Makler sperm counter (Cooper Surgical SEF-MAKL). One million motile sperm were then transferred into a new 500uL well of Research Vitro Fert Media or HTF in preparation for oocytes/cumulus masses.

For standard IVF, females were euthanized 14-16 hours post hCG injection and cumulus-oocyte complexes were extracted from the oviduct in pre-warmed MEM media (Thermo Fisher 12360038) and immediately transferred into well containing 1 million motile sperm. For oocytes generated through IVM), oocytes were removed from MEMa + Glutamax, rinsed at least once through HTF, and transferred to the well of HTF containing sperm. Except for the experiments with drug treatments, zygotes were allowed to incubate with the sperm until the time of collection. For extended embryo culture, fertilization was allowed to continue for 4-6 hours before the zygotes were removed from the well containing sperm and washed 5 times and incubated in pre-equilibrated KSOM AA (Millipore Sigma) at 37C and 5% CO2 and allowed to develop to morula or blastocyst stage embryos.

#### Drug treatments of zygotes

For inhibitor treatments, fertilization was allowed to continue for 4-6 hours before the zygotes were removed from the well containing sperm and placed in either, another pre-equilibrated 500 uL of Research Vitro Fert Media. Transcription-inhibited zygotes were treated with 5,6-dichloro-1-β-D ribofuranosyl-benzimidazole (Sigma) diluted 1:200 to 200 uM and 5-ethynyl uridine (Sigma) diluted 1:500 to 500 nM, both added to the IVF media 1.5 hours after fertilization, embryos were collected 10 hours after fertilization. Translation-inhibited zygotes were treated with cycloheximide (Sigma) diluted 1:500 to 100 ug/mL starting 1 hour after fertilization and O-propargyl-puromycin (Click Chemistry Tools 1493) diluted 1:500 to 20 uM in the IVF media 1 hour before collecting the zygotes 8.5 hours after fertilization. Zygotes with DNA replication inhibited were treated with aphidicolin (Sigma) diluted 1:500 to 10 ug/mL starting 1.5 hours after fertilization and 5-Ethynyl 2’-deoxyuridine (Sigma) diluted 1:500 to 20 uM added 1 hour before collection at 8.5 hours after fertilization. Zygotes cultured for controls were collected and fertilized alongside every replicate of the drug treatment experiments, and were stained with either EU, OPP, or EdU in the same manner. However, the control zygotes had dimethyl sulfoxide (Sigma) added to their IVF media at the same dilutions and time course as those zygotes treated with a small molecule inhibitor.

#### Immunofluorescence on oocytes and early embryos

For immunofluorescence, embryos or oocytes were collected at the indicated timepoints after washing in PBS treated briefly (30 seconds-1 minute) with Acidic Tyrodes solution (EMD Millipore) to remove the zona pellucida, and fixed in 4% paraformaldehyde (EMD Millipore) + PBS (Thermo Fisher) + 0.04% TritonX-100 (Sigma) + 0.3% Tween-20 (Sigma) + 0.2% sucrose (Sigma) for 10-15 minutes at 37C. Fixed embryos or oocytes were stored under mineral oil (Sigma) in PBS at 4C for no longer than one week. All staining took place in nine-well depression Pyrex plates (Millipore Sigma CLS722085) and utilized buffers which were made fresh and filtered through 0.45 um PES filters (Whatman 6780-2504). On the first day of immunofluorescence staining the embryos or oocytes were permeabilized in PBS + 0.5% TritonX-100 (Sigma) + 3% bovine serum albumin (BSA) (Sigma) at room temperature for 1 hour. If the embryos were incubated with EdU, 5-EU, or OPP (see above), these reagents, which each contain a terminal alkyne group, were next conjugated to a AZDye 488 picolyl azide following the Click Chemistry Tools kit protocol (CCT #1493). If no additional immunostaining was necessary the treated embryos were stained with DAPI (Sigma) diluted at 1:200 in PBS + 0.1% Triton + 3% BSA for 15-30 minutes at room temperature, washed twice for 10 minutes each with PBS + 0.1% Triton + 3% BSA, briefly dried onto a glass slide, then covered in Vectashield (Vector Laboratories) and a 22×22mm no. 1.5H coverslip (Thermo Fisher). After permeabilization and optional click-chemistry staining and/or phosphatase treatment (see above), embryos or oocytes were blocked in PBS + 0.1% Triton + 3% BSA + 10% fetal bovine serum (Thermo Fisher) at room temperature for 1-2 hours and stained with primary antibodies diluted in blocking buffer at 4C overnight: rabbit α CENP-A at 1:250 dilution (Cell Signaling Technologies 2048), rabbit α Mis18BP1 at 1:100 (Sigma HPA006504), rabbit α CENP-C at 1:1000 dilution (gift from Ben Black), or rabbit α CENP-C at 1:2000 dilution (gift from Iain Cheeseman). Embryos or oocytes were washed the following day in PBS with 0.1% Triton and 3% BSA five times for at least 15 minutes each, followed by incubation with DAPI at 1:200 and secondary antibodies diluted 1:500 in PBS + 0.1% Triton + 3% BSA + 10% fetal bovine serum for 2 hours at room temperature or overnight at 4C: Alexa Fluor 488 donkey α rabbit (Molecular Probes), Alexa Fluor 568 goat α rabbit (Molecular Probes). The stained embryos or oocytes were washed thrice for 10 minutes each with PBS + 0.1% Triton + 3% BSA, briefly dried onto a glass slide, then covered in Vectashield and a 22×22 mm no. 1.5H coverslip. Staining strategy was also used for human oocytes.

#### Oocyte collection and maturation

Oocytes were collected and matured *in vitro* as described previously (Stein & Schindler, 2011). Germinal vesicle (GV) stage mouse oocytes were collected from either CF-1 females (Charles River Laboratories), C57Bl/6J females (Jax), or *Cenpa^mScarlet/+^*at 3-4 or 6-16 weeks of age. The females were each injected with 5-7.5 IU pregnant mare’s serum gonadotropin (Sigma) or 100 uL HyperOva (Card KYD-010-EX-X5) intraperitoneally 44-48 hours prior to sacrifice. MEMa with GlutaMAX (Thermo Fisher 32561037) medium for oocyte collection and maturation were placed in an incubator to equilibrate to 37C and 5% CO2 the day before oocyte collection. Milrinone (Sigma, M4659-10mg) and fetal bovine serum (Thermo Fisher) were diluted to 2.5 uM and 5% respectively in both oocyte and collection media about an hour before oocyte collection. Females were euthanized and ovaries were dissected out and placed in pre-warmed MEM (Thermo Fisher 12360038) + 2.5 uM milrinone + 5% FBS + 3mg/mL Polyvinylpyrrolidone (Sigma, PVP10-100G). Oocyte-cumulus cell complexes were isolated by repeatedly poking the ovaries’ antral follicles with an insulin syringe needle. Oocytes were washed through several drops of MEM + 2.5 uM milrinone + 5% FBS and cumulus cells were mechanically detached by repeated mouth pipetting the complexes up and down. Cleaned GV oocytes were then washed and allowed to recover in MEMa with GlutaMAX (Thermo Fisher 32561037) + 2.5 uM milrinone + 5% FBS for several hours in 37C and 5% CO2. For maturation, the GV oocytes were washed away from milrinone by transferring them through at least 5 drops of MEMa with GlutaMAX (Thermo Fisher 32561037) + 5% FBS, at which point they were then cultured at 37C and 5% CO2. Metaphase I (MI) oocytes were collected 8 hours after release from milrinone and metaphase II (MII) oocytes were collected 16 hours after release from milrinone.

#### Oocyte microinjection

GV oocytes were collected as described above and microinjected according to previously published protocols (Stein and Schindler, 2011). Briefly, after letting GV oocytes recover from collection, the oocytes were transferred into MEM (Thermo Fisher 12360038) + 2.5 uM milrinone (Sigma) + 5% FBS (Thermo Fisher) and injected in groups of 20-30 with a FemtoJet 4i microinjector (Eppendorf). Holding micropipettes (Cooper Surgical MPH-MED-35) were cleaned with 70% ethanol (Sigma) and ddH2O, and injecting needles (Sutter B100-75-10) were pulled with a Sutter P97 or Sutter P2000 micropipette puller fresh before every injection. Injection pressure was adjusted for each needle, and injection volumes were approximately the same size as the GV (typically around 500 fPa).

Pools of siRNAs were purchased from Horizon Discovery and reconstituted in RNase-free dH2O according to the product manual. siRNA pools were stored in aliquots at –80C. On the day of injection, individual aliquots were thawed on ice, diluted to the appropriate concentration with RNase-free dH2O, and spun down at 20,000g for 30min at 4C. Pools used for this study are: CENP-A (J-044345-05-0002 through J-044345-08-0002, combined or L-044345-00-0005 pre-pooled), CENP-C (L-044347-00-0005), and negative control (D-001810-10-05).

In vitro transcribed RNA was prepared for injection by either linearizing the pIVT:: CENP-B-EGFP plasmid with PvuI (NEB) or double digesting pGEMHE: CENP-C plasmids with SbfI and PvuI (both NEB). 100-1000ng of linearized plasmid was then utilized as the template for a T7 in vitro transcription reaction with the mMessage mMachine T7 kit (Thermo Fisher AM1344), depending on the construct. RNA was purified with Qiagen’s RNeasy MinElute Cleanup Kit (74204) or with phenol chloroform per the instructions in the mMESSAGE mMachine T7 kit, aliquoted, and stored at –80C. While the collected GV oocytes were recovering, RNA integrity was checked by denaturing gel electrophoresis and good quality IVT RNA was combined, if necessary, with siRNA pools. RNA was diluted to reported concentrations in RNase-free dH2O, and spun down at 20,000g for 30min at 4C. Injected GV oocytes were allowed to recover in MEMa with GlutaMAX (Thermo Fisher 32561037) + 2.5 uM milrinone + 5% FBS for 1-6 hours at 37C and 5% CO2. The injected oocytes were matured and fertilized as described above.

siRNA-resistance was conferred in *Cenpc^WT^* IVT plasmid by introducing synonymous mutations within the seed regions for each siRNA in the targeting pool. The *Cenpc^Dimer^* IVT was generated from the *Cenpc^WT^* IVT plasmid, with point mutations derived from previous literature introduced into the corresponding mouse sequence residues (F829A and F901).

#### Oocyte photobleaching

GV oocytes were collected as described above and imaged and photobleached according to previously published protocols (Christodoulou et al., 2019; Ooga and Wakayama, 2017). For photobleaching, after recovering from collection, the GV oocytes were stained with 10ug/mL Hoechst 3324 (Sigma) for 30 minutes. Stained oocytes were then incubated in droplets of MEMa with GlutaMAX (Thermo Fisher 32561037) + 2.5 uM milrinone (Sigma) + 5% FBS (Thermo Fisher) pre-equilibrated to 37C at 5% CO2 under oil (NidOil, NO-400K). Droplets of oocytes were imaged and bleached in a MatTek 35mm, 1.5 coverslip, 10mm glass bottom dish (P35GC-1.5-10-C). The GV oocytes were incubated in a OkoLab live-cell imaging chamber (H301) on a Nikon Nikon A1R with a 60X 1.2NA long range water-immersion objective available at the University of Michigan’s Center for Live Cell Imaging through the Microscopy and Image Analysis Laboratory. Nikon NIS Elements was used to photobleach the mScarlet fluorescence in the nucleus of each GV oocyte with a circular ROI, the area adjusted to the size of each nucleus, with a 561nm laser line at 90uW of power and a 19.9uSec pixel dwell time. After photobleaching, the oocytes were allowed to recover for at least 30 minutes before being in-vitro matured and subsequently in-vitro fertilized as described above. All zygotes from photobleached, in-vitro matured, and in-vitro fertilized oocytes were collected after 8.5 hours of fertilization.

#### Generation of androgenetic zygotes by intracytoplasmic sperm injection (ICSI)

Ovulation was induced by intraperitoneal injection (i.p.) of equine chorionic gonadotropin (Pregmagon, IDT) and human chorionic gonadotropin (Ovogest, Intergonan). Lean B6C3F1 females aged 8 weeks and weighing approx. 25 g were injected i.p. using a 27G needle, at 5pm, with 10 I.U. eCG and 10 I.U. hCG 48 h apart.

Cumulus-oocyte complexes were collected from oviducts 14 hours after hCG at 7 am. The cumulus cells were dispersed in 50 U/mL hyaluronidase dissolved in Hepes-buffered Chatot, Ziomek and Bavister medium (HCZB) (Chatot et al., 1989) with bovine serum albumin (BSA) replaced by polyvinylpyrrolidone (PVP, 40 kDa) 0.1% w/v, at room temperature (28 °C). The cumulus-free MII oocytes were cultured in 500 µL of α-MEM medium (Sigma, M4526) supplemented with 0.2% (w/v) BSA and 50 mg/mL gentamicin, in 4-well plates (Nunc), in a 6% CO 2 incubator at 37 °C until further processing.

Diploid androgenetic embryos were generated by stripping the oocytes of their meiotic spindle followed by injection of two sperm heads, on the stage of a NikonTE2000 inverted microscope fitted with Nomarski optics, at room temperature (28 °C). To remove the meiotic spindle, oocytes were placed in a 300 µL elongated drop of HCZB medium supplemented with 0.1 % PVP and 5 µM Latrunculin B (cat. no. 428020, Merck Millipore, Darmstadt, Germany), on a glass-bottomed dish (Figure 2). A hole was drilled in the zona pellucida using a mercury-filled, piezo-driven needle (inner diameter 10-12 µm), and the spindle was gently aspirated (Figure 3). Resultant ooplasts were allowed to recover in α-MEM medium for at least 2 hours. For ICSI, ooplasts were placed in a 300 µL elongated drop of HCZB medium supplemented with 0.5 % (w/v) PVP on a glass-bottomed dish as described. Each ooplast was injected with two sperm heads simultaneously using a mercury-filled, piezo-driven needle (inner diameter 7-8 µm) (Figure 4). Ten minutes after the ICSI the oocytes were transferred to 500 mL of Potassium (K) simplex optimization medium (KSOM) in 4-well plates, in a 6% CO2 incubator at 37 °C. KSOM was prepared in house as per the original recipe (Summers et al., 2000) and contained free aminoacids both essential and non-essential, 0.2% (w/v) BSA and gentamicin (50 mg/mL). Controls for the diploid androgenetic embryos were generated by ICSI of single sperm heads in intact oocytes. Both groups were cultured in KSOM medium to allow for oocyte activation and pronuclear formation, until further processing.

After 16 hours of cultures, the embryos were fixed in 3.7% PFA for 1 hour at room temperature, then permeabilized in 0.5% Tritonx-100 for 20 minutes, and then blocked/stored for transport 2% BSA + 2% glycine + 0.1% Tween-20 + 5% Donkey Serum. Subsequent immunofluorescence staining was performed as described above, beginning at the phosphatase treatment.

### QUANTIFICATION AND STATISTICAL ANALYSIS

#### Quantification of centromere signals

Quantification of centromere fluorescence intensity was made in Fiji/ImageJ software by drawing circles of constant diameter around individual fluorescent puncta associated with DNA stains across all Z-planes of the cell. The total intensity was calculated for each punctum after subtracting the background of the image, which was obtained from un-stained regions of the cell nucleus for each cell and then summing the total puncta fluorescence intensities for each pronuclei or cell being studied (see **Supplemental Video 1**). For each statistical test, multiple independent technical and biological replicates were quantified to obtain a minimum of about ten embryos per experiment.

#### Quantification of MIS18BP1 staining in zygotes

Quantification of centromere fluorescence intensity was made in Fiji/ImageJ software by drawing circles around the maternal and paternal pronuclei at the z-slice in which each pronucleus was largest. Raw integrated intensity measurements are normalized by pronuclear area and plotted as the paternal/maternal ratio where reported.

#### Quantification of mis-segregated chromosomes

For 2-cell embryos that were scored for mis-segregated chromosomes, embryos were collected 24hpf to allow sufficient time for nuclear envelope formation to occur in both blastomeres. Mis-segregated chromosome counts were determined by counting the number of DAPI-stained structures outside of the nuclear envelopes in both blastomeres. Individual chromosome counts were not determined.

#### Statistical testing

All statistical tests for significance were performed in either R or GraphPad Prism 9 or 10 software. P-values were calculated at a significance level of 0.05 or 95% confidence interval for all two tailed statistical tests. Figures were made in R and GraphPad Prism 9 software. The number of replicates for each experiment is supplied in the figure legends. Tests for normal distribution of the data were performed in R using both the shapiro.test() and leveneTest() functions. Normally distributed paired data was tested using the t.test() and non-parametric data was tested using the wilcox.test(). In GraphPad, the Shapiro-Wilk normality test was applied to determine if the data was normally distributed, and the appropriate normal or non-parametric t-test was applied to the data accordingly.

Statistical testing for 2-cell developmental progression and 2-cell mis-segregation rates were performed in R. P-values were calculated at a significance level of 0.05 using the prop.test() function.

#### Representative Images

Representative images were acquired primary using Fiji/ImageJ software to create maximum intensity projections using selective z-slices from zygotes to collapse both pronuclei into one image. Additional images were acquired using ImarisViewer 10.1.0. Images were imported and opened in the “Section” view with extended settings to create maximum intensity projections.

#### Representative Video

A video demonstrating the identification and quantification of puncta was recorded using Apple’s built-in screen recording feature on a device running iOS 18.4. The recorded footage was subsequently edited using iMovie version 10.4.3

## References

1. Abe, K., Funaya, S., Tsukioka, D., Kawamura, M., Suzuki, Y., Suzuki, M.G., Schultz, R.M., and Aoki, F. (2018). Minor zygotic gene activation is essential for mouse preimplantation development. Proc. Natl. Acad. Sci. 115, 201804309.

2. Adenot, P.G., Mercier, Y., Renard, J.P., and Thompson, E.M. (1997). Differential H4 acetylation of paternal and maternal chromatin precedes DNA replication and differential transcriptional activity in pronuclei of 1-cell mouse embryos. Development 124, 4615–4625.

3. Amor, D.J., Bentley, K., Ryan, J., Perry, J., Wong, L., Slater, H., and Choo, K.H.A. (2004). Human centromere repositioning “in progress.” Proc. Natl. Acad. Sci. U. S. A. 101, 6542–6547.

4. Ballow, D., Meistrich, M.L., Matzuk, M., and Rajkovic, A. (2006). Sohlh1 is essential for spermatogonial differentiation. Dev. Biol. 294, 161–167.

5. Bao, J., and Bedford, M.T. (2016). Epigenetic regulation of the histone-to-protamine transition during spermiogenesis. Reproduction 151, R55–R70.

6. Baran, V., Solc, P., Kovarikova, V., Rehak, P., and Sutovsky, P. (2013). Polo-like kinase 1 is essential for the first mitotic division in the mouse embryo. Mol. Reprod. Dev. 80, 522–534.

7. Barra, V., and Fachinetti, D. (2018). The dark side of centromeres: types, causes and consequences of structural abnormalities implicating centromeric DNA. Nat. Commun. 9.

8. Bindels, D.S., Haarbosch, L., Weeren, L. Van, Postma, M., Wiese, K.E., Mastop, M., Aumonier, S., Gotthard, G., Royant, A., Hink, M.A., et al. (2016). mScarlet: a bright monomeric red fluorescent protein for cellular imaging. Nat. Publ. Gr. 14, 53–56.

9. Bodor, D.L., Valente, L.P., Mata, J.F., Black, B.E., and Jansen, L.E.T. (2013). Assembly in G1 phase and long-term stability are unique intrinsic features of CENP-A nucleosomes. Mol. Biol. Cell 24, 923–932.

10. Bodor, D. L., Mata, J. F., Sergeev, M., David, A. F., Salimian, K. J., Panchenko, T., Cleveland, D. W., Black, B. E., Shah, J. V., & Jansen, L. E. (2014). The quantitative architecture of centromeric chromatin. eLife, 3, e02137. 10.7554/eLife.02137

11. Boyarchuk, E., Filipescu, D., Vassias, I., Cantaloube, S., and Almouzni, G. (2014). The histone variant composition of centromeres is controlled by the pericentric heterochromatin state during the cell cycle. J. Cell Sci. 127, 3347–3359.

12. Brinkley, B. R., Brenner, S. L., Hall, J. M., Tousson, A., Balczon, R. D., & Valdivia, M. M. (1986). Arrangements of kinetochores in mouse cells during meiosis and spermiogenesis. Chromosoma, 94(4), 309–317. 10.1007/BF00290861

13. Brykczynska, U., Hisano, M., Erkek, S., Ramos, L., Oakeley, E.J., Roloff, T.C., Beisel, C., Schübeler, D., Stadler, M.B., and Peters, A.H.F.M. (2010). Repressive and active histone methylation mark distinct promoters in human and mouse spermatozoa. Nat. Struct. Mol. Biol. 17, 679–687.

14. Buageaw, A., Sukhwani, M., Ben-Yehudah, A., Ehmcke, J., Rawe, V.Y., Pholpramool, C., Orwig, K.E., and Schlatt, S. (2005). GDNF family receptor alpha1 phenotype of spermatogonial stem cells in immature mouse testes. Biol. Reprod. 73, 1011–1016.

15. Bukvic, N., Susca, F., Gentile, M., Tangari, E., Ianniruberto, A., and Guanti, G. (1996). An unusual dicentric Y chromosome with a functional centromere with no detectable alpha-satellite. Hum. Genet. 97, 453–456.

16. Carroll, C. W., Milks, K. J., & Straight, A. F. (2010). Dual recognition of CENP-A nucleosomes is required for centromere assembly. The Journal of cell biology, 189(7), 1143–1155. 10.1083/jcb.201001013

17. Carty, B.L., Dattoli, A.A., Carty, B.L., Dattoli, A.A., and Dunleavy, E.M. (2021). CENP-C functions in centromere assembly, the maintenance of CENP-A asymmetry and epigenetic age in Drosophila germline stem cells. PLoS Genet. 17, 1–23.

18. Chan, F., Oatley, M.J., Kaucher, A. V, Yang, Q., Bieberich, C.J., Shashikant, C.S., and Oatley, J.M. (2014). Functional and molecular features of the Id4 + germline stem cell population in mouse testes. Genes Dev. 28, 1351–1362.

19. Chmátal, L., Gabriel, S.I., Mitsainas, G.P., Martínez-Vargas, J., Ventura, J., Searle, J.B., Schultz, R.M., and Lampson, M.A. (2014). Centromere strength provides the cell biological basis for meiotic drive and karyotype evolution in mice. Curr. Biol. 24, 2295–2300.

20. Christodoulou, N., Weberling, A., and Zernicka-Goetz, M. (2019). Live imaging of mouse embryos during pre-implantation and peri-implantation development. 1–10.

21. Chuang, T. H., Chang, Y. P., Lee, M. J., Wang, H. L., Lai, H. H., & Chen, S. U. (2021). The Incidence of 1 Mosaicism for Individual Chromosome in Human Blastocysts Is Correlated With Chromosome Length. Frontiers in genetics, 11, 565348. 10.3389/fgene.2020.565348

22. Conti, D., Verza, A. E., Pesenti, M. E., Cmentowski, V., Vetter, I. R., Pan, D., & Musacchio, A. (2024). Role of protein kinase PLK1 in the epigenetic maintenance of centromeres. Science (New York, N.Y.), 385(6713), 1091–1097. 10.1126/science.ado5178

23. Das, A., Black, B. E., & Lampson, M. A. (2020). Maternal inheritance of centromeres through the germline. Current topics in developmental biology, 140, 35–54. 10.1016/bs.ctdb.2020.03.004

24. Das, A., Iwata-Otsubo, A., Destouni, A., Dawicki-McKenna, J.M., Boese, K.G., Black, B.E., and Lampson, M.A. (2022). Epigenetic, genetic and maternal effects enable stable centromere inheritance. Nat. Cell Biol. 24, 748–756.

25. Das, A., Boese, K. G., Tachibana, K., Baek, S. H., Lampson, M. A., & Black, B. E. (2023). Centromere-specifying nucleosomes persist in aging mouse oocytes in the absence of nascent assembly. Current biology: CB, 33(17), 3759–3765.e3.

26. Dattoli, A.A., Carty, B.L., Kochendoerfer, A.M., Morgan, C., Walshe, A.E., and Dunleavy, E.M. (2020). Asymmetric assembly of centromeres epigenetically regulates stem cell fate. J. Cell Biol. 219.

27. Dunleavy, E.M., Beier, N.L., Gorgescu, W., Tang, J., Costes, S. V., and Karpen, G.H. (2012). The Cell Cycle Timing of Centromeric Chromatin Assembly in Drosophila Meiosis Is Distinct from Mitosis Yet Requires CAL1 and CENP-C. PLoS Biol. 10, 1–16.

28. van Echten-Arends, J., Mastenbroek, S., Sikkema-Raddatz, B., Korevaar, J. C., Heineman, M. J., van der Veen, F., & Repping, S. (2011). Chromosomal mosaicism in human preimplantation embryos: a systematic review. Human reproduction update, 17(5), 620–627. 10.1093/humupd/dmr014

29. Endo, T., Romer, K.A., Anderson, E.L., Baltus, A.E., de Rooij, D.G., and Page, D.C. (2015). Periodic retinoic acid–STRA8 signaling intersects with periodic germ-cell competencies to regulate spermatogenesis. Proc. Natl. Acad. Sci. 112, E2347–E2356.

30. Fachinetti, D., Folco, H. D., Nechemia-Arbely, Y., Valente, L. P., Nguyen, K., Wong, A. J., Zhu, Q., Holland, A. J., Desai, A., Jansen, L. E., & Cleveland, D. W. (2013). A two-step mechanism for epigenetic specification of centromere identity and function. Nature cell biology, 15(9), 1056–1066. 10.1038/ncb2805

31. Fachinetti, D., Han, J.S., McMahon, M.A., Ly, P., Abdullah, A., Wong, A.J., and Cleveland, D.W. (2015). DNA Sequence-Specific Binding of CENP-B Enhances the Fidelity of Human Centromere Function. Dev. Cell 33, 314–327.

32. Falk, S.J., Guo, L.Y., Sekulic, N., Smoak, E.M., Mani, T., Logsdon, G.A., Gupta, K., Jansen, L.E.T., Van Duyne, G.D., Vinogradov, S.A., et al. (2015). CENP-C reshapes and stabilizes CENP-A nucleosomes at the centromere. Science (80-.). 348, 699–703.

33. Flores Servin, J. C., Brown, R. R., & Straight, A. F. (2023). Repression of CENP-A assembly in metaphase requires HJURP phosphorylation and inhibition by M18BP1. The Journal of cell biology, 222(6), e202110124. 10.1083/jcb.202110124

34. Foltz, D.R., Jansen, L.E.T., Black, B.E., Bailey, A.O., Yates, J.R., and Cleveland, D.W. (2006). The human CENP-A centromeric nucleosome-associated complex. Nat. Cell Biol. 8, 458–469.

35. Fujita, R., Otake, K., Arimura, Y., Horikoshi, N., Miya, Y., Shiga, T., Osakabe, A., Tachiwana, H., Ohzeki, J.I., Larionov, V., et al. (2015). Stable complex formation of CENP-B with the CENP-A nucleosome. Nucleic Acids Res. 43, 4909–4922.

36. García del Arco, A., Edgar, B.A., and Erhardt, S. (2018). In Vivo Analysis of Centromeric Proteins Reveals a Stem Cell-Specific Asymmetry and an Essential Role in Differentiated, Non-proliferating Cells. Cell Rep. 22, 1982–1993.

37. Gassmann, R., Rechtsteiner, A., Yuen, K.W., Muroyama, A., Egelhofer, T., Gaydos, L., Barron, F., Maddox, P., Essex, A., Monen, J., et al. (2012). An inverse relationship to germline transcription defines centromeric chromatin in C. elegans. Nature 484, 534–537.

38. Gaysinskaya, V., Soh, I.Y., van der Heijden, G.W., and Bortvin, A. (2014). Optimized flow cytometry isolation of murine spermatocytes. Cytom. Part A 85, 556–565.

39. Giunta, S., and Funabiki, H. (2017). Integrity of the human centromere DNA repeats is protected by CENP-A, CENP-C, and CENP-T. Proc. Natl. Acad. Sci. U. S. A. 114, 1928–1933.

40. Green, C.D., Ma, Q., Manske, G.L., Shami, A.N., Zheng, X., Marini, S., Moritz, L., Sultan, C., Gurczynski, S.J., Moore, B.B., et al. (2018). A Comprehensive Roadmap of Murine Spermatogenesis Defined by Single-Cell RNA-Seq. Dev. Cell 46, 651–667.e10.

41. Guo, L.Y., Allu, P.K., Zandarashvili, L., McKinley, K.L., Sekulic, N., Dawicki-McKenna, J.M., Fachinetti, D., Logsdon, G.A., Jamiolkowski, R.M., Cleveland, D.W., et al. (2017). Centromeres are maintained by fastening CENP-A to DNA and directing an arginine anchor-dependent nucleosome transition. Nat. Commun. 8, 15775.

42. Haeussler, M., Schönig, K., Eckert, H., Eschstruth, A., Mianné, J., Renaud, J.B., Schneider-Maunoury, S., Shkumatava, A., Teboul, L., Kent, J., et al. (2016). Evaluation of off-target and on-target scoring algorithms and integration into the guide RNA selection tool CRISPOR. Genome Biol. 17, 1–12.

43. Hammoud, S. S., Nix, D. A., Zhang, H., Purwar, J., Carrell, D. T., & Cairns, B. R. (2009). Distinctive chromatin in human sperm packages genes for embryo development. Nature, 460(7254), 473–478. 10.1038/nature08162

44. Hara, M., Ariyoshi, M., Sano, T., Nozawa, R. S., Shinkai, S., Onami, S., Jansen, I., Hirota, T., & Fukagawa, T. (2023). Centromere/kinetochore is assembled through CENP-C oligomerization. Molecular cell, 83(13), 2188–2205.e13. 10.1016/j.molcel.2023.05.023

45. van der Heijden, G.W., Ramos, L., Baart, E.B., van den Berg, I.M., Derijck, A.A., van der Vlag, J., Martini, E., and de Boer, P. (2008). Sperm-derived histones contribute to zygotic chromatin in humans. BMC Dev. Biol. 8, 34.

46. Helsel, A.R., Yang, Q.E., Oatley, M.J., Lord, T., Sablitzky, F., and Oatley, J.M. (2017). Id4 levels dictate the stem cell state in mouse spermatogonia. Dev. 144, 624–634.

47. Hemmerich, P., Weidtkamp-Peters, S., Hoischen, C., Schmiedeberg, L., Erliandri, I., and Diekmann, S. (2008). Dynamics of inner kinetochore assembly and maintenance in living cells. J. Cell Biol. 180, 1101–1114.

48. Hoffmann, S., Dumont, M., Barra, V., Ly, P., Nechemia-Arbely, Y., McMahon, M. A., Hervé, S., Cleveland, D. W., & Fachinetti, D. (2016). CENP-A Is Dispensable for Mitotic Centromere Function after Initial Centromere/Kinetochore Assembly. Cell reports, 17(9), 2394–2404. 10.1016/j.celrep.2016.10.084

49. Howman, E. V., Fowler, K. J., Newson, A. J., Redward, S., MacDonald, A. C., Kalitsis, P., & Choo, K. H. (2000). Early disruption of centromeric chromatin organization in centromere protein A (Cenpa) null mice. Proceedings of the National Academy of Sciences of the United States of America, 97(3), 1148–1153. 10.1073/pnas.97.3.1148

50. Hu, H., Liu, Y., Wang, M., Fang, J., Huang, H., Yang, N., Li, Y., Wang, J., Yao, X., Shi, Y., et al. (2011). Structure of a CENP-A-histone H4 heterodimer in complex with chaperone HJURP. Genes Dev. 25, 901–906.

51. Huang, H., Strømme, C.B., Saredi, G., Hödl, M., Strandsby, A., González-aguilera, C., Chen, S., Groth, A., and Patel, D.J. (2015). A unique binding mode enables MCM2 to chaperone histones H3 – H4 at replication forks. Nat. Publ. Gr. 22, 618–626.

52. Ikegami, S., Taguchi, T., Ohashi, M., Oguro, M., Nagano, H., And Mano, Y. (1978). Aphidicolin prevents mitotic cell division by interfering with the activity of DNA polymerase-α. Nature 275, 458–460.

53. Ingouff, M., Rademacher, S., Holec, S., Šoljić, L., Xin, N., Readshaw, A., Foo, S.H., Lahouze, B., Sprunck, S., and Berger, F. (2010). Zygotic resetting of the HISTONE 3 variant repertoire participates in epigenetic reprogramming in arabidopsis. Curr. Biol. 20, 2137–2143.

54. Iwata-Otsubo, A., Dawicki-McKenna, J.M., Akera, T., Falk, S.J., Chmátal, L., Yang, K., Sullivan, B.A., Schultz, R.M., Lampson, M.A., and Black, B.E. (2017). Expanded Satellite Repeats Amplify a Discrete CENP-A Nucleosome Assembly Site on Chromosomes that Drive in Female Meiosis. Curr. Biol. 27, 2365–2373.e8.

55. Jansen, L.E.T., Black, B.E., Foltz, D.R., and Cleveland, D.W. (2007). Propagation of centromeric chromatin requires exit from mitosis. J. Cell Biol. 176, 795–805.

56. Jao, C.Y., and Salic, A. (2008). Exploring RNA transcription and turnover in vivo by using click chemistry. Proc. Natl. Acad. Sci. U. S. A. 105, 15779–15784.

57. Jeffery, D., Gatto, A., Podsypanina, K., Renaud-Pageot, C., Ponce Landete, R., Bonneville, L., Dumont, M., Fachinetti, D., and Almouzni, G. (2021). CENP-A overexpression promotes distinct fates in human cells, depending on p53 status. Commun. Biol. 4, 1–18.

58. Jones, K.T. (2008). Meiosis in oocytes: Predisposition to aneuploidy and its increased incidence with age. Hum. Reprod. Update 14, 143–158.

59. Kalitsis, P., Fowler, K. J., Earle, E., Hill, J., & Choo, K. H. (1998). Targeted disruption of mouse centromere protein C gene leads to mitotic disarray and early embryo death. Proceedings of the National Academy of Sciences of the United States of America, 95(3), 1136–1141. 10.1073/pnas.95.3.1136

60. Kalitsis, P., Griffiths, B., and Choo, K.H.A. (2006). Mouse telocentric sequences reveal a high rate of homogenization and possible role in Robertsonian translocation. Proc. Natl. Acad. Sci. U. S. A. 103, 8786– 8791.

61. Kwon, M. S., Hori, T., Okada, M., & Fukagawa, T. (2007). CENP-C is involved in chromosome segregation, mitotic checkpoint function, and kinetochore assembly. Molecular biology of the cell, 18(6), 2155–2168.

62. Larose, H., Shami, A.N., Abbott, H., Manske, G., Lei, L., and Hammoud, S.S. (2019). Gametogenesis: A journey from inception to conception (Elsevier Inc.).

63. Lewis, J.D., Song, Y., De Jong, M.E., Bagha, S.M., and Ausió, J. (2003). A walk though vertebrate and invertebrate protamines. Chromosoma 111, 473–482.

64. Li, Y., Zhu, Z., Zhang, S., Yu, D., Yu, H., Liu, L., Cao, X., Wang, L., Gao, H., and Zhu, M. (2011). Shrna-targeted centromere protein a inhibits hepatocellular carcinoma growth. PLoS One 6, 19–23.

65. Liu, J., Xu, Y., Stoleru, D., and Salic, A. (2012). Imaging protein synthesis in cells and tissues with an alkyne analog of puromycin. Proc. Natl. Acad. Sci. U. S. A. 109, 413–418.

66. Maehara, K., Takahashi, K., and Saitoh, S. (2010). CENP-A Reduction Induces a p53-Dependent Cellular Senescence Response To Protect Cells from Executing Defective Mitoses. Mol. Cell. Biol. 30, 2090–2104.

67. Mahadevaiah, S.K., Turner, J.M.A., Baudat, F., Rogakou, E.P., De Boer, P., Blanco-Rodríguez, J., Jasin, M., Keeney, S., Bonner, W.M., and Burgoyne, P.S. (2001). Recombinational DNA double-strand breaks in mice precede synapsis. Nat. Genet. 27, 271–276.

68. McKinley, K.L., and Cheeseman, I.M. (2014). Polo-like kinase 1 licenses CENP-a deposition at centromeres. Cell 158, 397–411.

69. McKinley, K. L., Sekulic, N., Guo, L. Y., Tsinman, T., Black, B. E., & Cheeseman, I. M. (2015). The CENP-L-N Complex Forms a Critical Node in an Integrated Meshwork of Interactions at the Centromere-Kinetochore Interface. Molecular cell, 60(6), 886–898.

70. McKinley, K.L., and Cheeseman, I.M. (2016). The molecular basis for centromere identity and function. Nat. Rev. Mol. Cell Biol. 17, 16–29.

71. Mellone, B.G., and Fachinetti, D. (2021). Diverse mechanisms of centromere specification. Curr. Biol. 31, R1491–R1504.

72. Milks, K.J., Moree, B., and Straight, A.F. (2009). Dissection of CENP-C-directed centromere and kinetochore assembly. Mol. Biol. Cell 20, 4246–4255.

73. Mitra, S., Bodor, D.L., David, A.F., Abdul-Zani, I., Mata, J.F., Neumann, B., Reither, S., Tischer, C., and Jansen, L.E.T. (2020). Genetic screening identifies a SUMO protease dynamically maintaining centromeric chromatin. Nat. Commun. 11, 501.

74. Moree, B., Meyer, C. B., Fuller, C. J., & Straight, A. F. (2011). CENP-C recruits M18BP1 to centromeres to promote CENP-A chromatin assembly. The Journal of cell biology, 194(6), 855–871.

75. Moritz, L., Schon, S. B., Rabbani, M., Sheng, Y., Agrawal, R., Glass-Klaiber, J., Sultan, C., Camarillo, J. M., Clements, J., Baldwin, M. R., Diehl, A. G., Boyle, A. P., O’Brien, P. J., Ragunathan, K., Hu, Y. C., Kelleher, N. L., Nandakumar, J., Li, J. Z., Orwig, K. E., Redding, S., Hammoud, S. S. (2023). Sperm chromatin structure and reproductive fitness are altered by substitution of a single amino acid in mouse protamine 1. Nature structural & molecular biology, 30(8), 1077–1091.

76. Müller, S., MontesdeOca, R., Lacoste, N., Dingli, F., Loew, D., and Almouzni, G. (2014). Phosphorylation and DNA binding of HJURP determine its centromeric recruitment and function in CenH3CENP-A loading. Cell Rep. 8, 190–203.

77. Nechemia-Arbely, Y., Miga, K.H., Shoshani, O., Aslanian, A., McMahon, M.A., Lee, A.Y., Fachinetti, D., Yates, J.R., Ren, B., and Cleveland, D.W. (2019). DNA replication acts as an error correction mechanism to maintain centromere identity by restricting CENP-A to centromeres. Nat. Cell Biol. 21, 743–754.

78. Ooga, M., and Wakayama, T. (2017). FRAP analysis of chromatin looseness in mouse zygotes that allows full-Term development. PLoS One 12.

79. Oqani, R.K., Kim, H.R., Diao, Y.F., Park, C.S., and Jin, D.I. (2011). The CDK9/Cyclin T1 subunits of P-TEFb in mouse oocytes and preimplantation embryos: A possible role in embryonic genome activation. BMC Dev. Biol. 11, 1–9.

80. Palmer, D.K., O’Day, K., and Margolis, R.L. (1990). The centromere specific histone CENP-A is selectively retained in discrete foci in mammalian sperm nuclei. Chromosoma 100, 32–36.

81. Palmer, D.K., O’Day, K., Trong, H.L.E., Charbonneau, H., and Margolis, R.L. (1991). Purification of the centromere-specific protein CENP-A and demonstration that it is a distinctive histone. Proc. Natl. Acad. Sci. U. S. A. 88, 3734–3738.

82. Pan, D., Klare, K., Petrovic, A., Take, A., Walstein, K., Singh, P., Rondelet, A., Bird, A.W., and Musacchio, A. (2017). CDK-regulated dimerization of M18BP1 on a Mis18 hexamer is necessary for CENP-A loading. Elife 6, 1–25.

83. Pan, D., Walstein, K., Take, A., Bier, D., Kaiser, N., and Musacchio, A. (2019). Mechanism of centromere recruitment of the CENP-A chaperone HJURP and its implications for centromere licensing. Nat. Commun. 10, 1–18.

84. Parashara, P., Medina-Pritchard, B., Abad, M. A., Sotelo-Parrilla, P., Thamkachy, R., Grundei, D., Zou, J., Spanos, C., Kumar, C. N., Basquin, C., Das, V., Yan, Z., Al-Murtadha, A. A., Kelly, D. A., McHugh, T., Imhof, A., Rappsilber, J., & Jeyaprakash, A. A. (2024). PLK1-mediated phosphorylation cascade activates Mis18 complex to ensure centromere inheritance. *Science (New York*, N.Y.), 385(6713), 1098–1104. 10.1126/science.ado8270

85. Pentakota, S., Zhou, K., Smith, C., Maffini, S., Petrovic, A., Morgan, G.P., Weir, J.R., Vetter, I.R., Musacchio, A., and Luger, K. (2017). Decoding the centromeric nucleosome through CENP-N. Elife 6, 1–25.

86. Pesenti, M.E., Raisch, T., Conti, D., Walstein, K., Hoffmann, I., Vogt, D., Prumbaum, D., Vetter, I.R., Raunser, S., and Musacchio, A. (2022). Structure of the human inner kinetochore CCAN complex and its significance for human centromere organization. Mol. Cell 1–19.

87. Ranjan, R., Snedeker, J., and Chen, X. (2019). Asymmetric Centromeres Differentially Coordinate with Mitotic Machinery to Ensure Biased Sister Chromatid Segregation in Germline Stem Cells. Cell Stem Cell 25, 666–681.e5.

88. Ranjan, R. & Chen, X. Quantitative imaging of chromatin inheritance using a dual-color histone in Drosophila germinal stem cells. STAR Protocols 3, 101811 (2022).

89. Raychaudhuri, N., Dubruille, R., Orsi, G.A., Bagheri, H.C., Loppin, B., and Lehner, C.F. (2012). Transgenerational Propagation and Quantitative Maintenance of Paternal Centromeres Depends on Cid/Cenp-A Presence in Drosophila Sperm. PLoS Biol. 10.

90. Schubert, V., Lermontova, I., and Schubert, I. (2014). Loading of the centromeric histone H3 variant during meiosis–how does it differ from mitosis? Chromosoma 123, 491–497.

91. Silva, M.C.C., Bodor, D.L., Stellfox, M.E., Martins, N.M.C., Hochegger, H., Foltz, D.R., and Jansen, L.E.T. (2012). Cdk Activity Couples Epigenetic Centromere Inheritance to Cell Cycle Progression. Dev. Cell 22, 52–63.

92. Smith, B.E., and Braun, R.E. (2012). Germ Cell Migration Across Sertoli Cell Tight Junctions. Science (80-.). 338, 798–802.

93. Smoak, E.M., Stein, P., Schultz, R.M., Lampson, M.A., and Black, B.E. (2016). Long-Term Retention of CENP-A Nucleosomes in Mammalian Oocytes Underpins Transgenerational Inheritance of Centromere Identity. Curr. Biol. 26, 1110–1116.

94. Spiller, F., Medina-Pritchard, B., Abad, M.A., Wear, M.A., Molina, O., Earnshaw, W.C., and Jeyaprakash, A.A. (2017). Molecular basis for Cdk1-regulated timing of Mis18 complex assembly and CENP-A deposition. EMBO Rep. 18, 894–905.

95. Srivastava, S., and Foltz, D.R. (2018). Posttranslational modifications of CENP-A: marks of distinction. Chromosoma 127, 279–290.

96. Stankovic, A., Guo, L.Y., Mata, J.F., Bodor, D.L., Cao, X.J., Bailey, A.O., Shabanowitz, J., Hunt, D.F., Garcia, B.A., Black, B.E., et al. (2017). A Dual Inhibitory Mechanism Sufficient to Maintain Cell-Cycle-Restricted CENP-A Assembly. Mol. Cell 65, 231–246.

97. Stein, P., and Schindler, K. (2011). Mouse oocyte microinjection, maturation and ploidy assessment. J. Vis. Exp. 1–5.

98. Štiavnická, M., Keegan, R. S., & Dunleavy, E. M. (2025). Marking dad’s centromeres: maintaining CENP-A in sperm. Chromosome research: an international journal on the molecular, supramolecular and evolutionary aspects of chromosome biology, 33(1), 8. 10.1007/s10577-025-09766-2

99. Sun, F., Xu, Q., Zhao, D., and Degui Chen, C. (2015). Id4 Marks Spermatogonial Stem Cells in the Mouse Testis. Sci. Rep. 5, 2–13.

100. Swartz, S.Z., McKay, L.S., Su, K.C., Bury, L., Padeganeh, A., Maddox, P.S., Knouse, K.A., and Cheeseman, I.M. (2019). Quiescent Cells Actively Replenish CENP-A Nucleosomes to Maintain Centromere Identity and Proliferative Potential. Dev. Cell 51, 35–48.e7.

101. Tyler-Smith, C., Gimelli, G., Giglio, S., Floridia, G., Pandya, A., Terzoli, G., Warburton, P.E., Earnshaw, W.C., and Zuffardi, O. (1999). Transmission of a fully functional human neocentromere through three generations. Am. J. Hum. Genet. 64, 1440–1444.

102. Van De Werken, C., Van Der Heijden, G.W., Eleveld, C., Teeuwssen, M., Albert, M., Baarends, W.M., Laven, J.S.E., Peters, A.H.F.M., and Baart, E.B. (2014). Paternal heterochromatin formation in human embryos is H3K9/HP1 directed and primed by sperm-derived histone modifications. Nat. Commun. 5, 1–15.

103. Vázquez-Diez, C., & FitzHarris, G. (2018). Causes and consequences of chromosome segregation error in preimplantation embryos. *Reproduction (Cambridge*, England), 155(1), R63–R76. 10.1530/REP-17-0569

104. Xiong, Z., Xu, K., Lin, Z., Kong, F., Wang, Q., Quan, Y., Sha, Q. Q., Li, F., Zou, Z., Liu, L., Ji, S., Chen, Y., Zhang, H., Fang, J., Yu, G., Liu, B., Wang, L., Wang, H., Deng, H., Yang, X., Fan, H., Li, L., Xie, W. (2022). Ultrasensitive Ribo-seq reveals translational landscapes during mammalian oocyte-to-embryo transition and pre-implantation development. Nature cell biology, 24(6), 968–980.

105. Yoshida, K., Muratani, M., Araki, H., Miura, F., Suzuki, T., Dohmae, N., Katou, Y., Shirahige, K., Ito, T., and Ishii, S. (2018). Mapping of histone-binding sites in histone replacement-completed spermatozoa. Nat. Commun. 9, 1–11.

106. Zasadzińska, E., Huang, J., Bailey, A.O., Guo, L.Y., Lee, N.S., Srivastava, S., Wong, K.A., French, B.T., Black, B.E., and Foltz, D.R. (2018). Inheritance of CENP-A Nucleosomes during DNA Replication Requires HJURP. Dev. Cell 47, 348–362.e7.

107. Zelazowski, M.J., Sandoval, M., Paniker, L., Hamilton, H.M., Han, J., Gribbell, M.A., Kang, R., and Cole, F. (2017). Age-Dependent Alterations in Meiotic Recombination Cause Chromosome Segregation Errors in Spermatocytes. Cell 171, 601–614.e13.

108. Zhou, Q., Nie, R., Li, Y., Friel, P., Mitchell, D., Hess, R.A., Small, C., and Griswold, M.D. (2008). Expression of stimulated by retinoic acid gene 8 (Stra8) in Spermatogenic Cells Induced by retinoic acid: An in vivo study in vitamin A-sufficient postnatal murine testes. Biol. Reprod. 79, 35–42.

